# Immunoglobulin switch-like recombination regions implicated in the formation of extrachromosomal circular 45S rDNA involved in the maternal-specific translation system of zebrafish

**DOI:** 10.1101/2020.01.31.928739

**Authors:** Timo M. Breit, Han Rauwerda, Johanna F. B. Pagano, Wim A. Ensink, Ulrike Nehrdich, Herman P. Spaink, Rob J. Dekker

## Abstract

Cellular translation is essential to all life on earth and in recent years we have reported on the discovery of a unique dual translation system in zebrafish. In this system, a maternal-type variant shows absolute expression in eggs and is progressively replaced during embryogenesis by a somatic-type variant. There are several translation system components, all with a non-coding RNA part, that show this dual characteristic: snRNA, snoRNA, rRNA, RNaseP, tRNA, and SRP-RNA.

To produce sufficient ribosomes during oogenesis, zebrafish amplify their 45S locus (18S-5.8S-28S tandem repeat) by means of extrachromosomal circular DNA (eccDNA) organized in extrachromosomal rDNA circles (ERCs). Although this cellular process is discovered quite some time ago, still little is known about the mechanisms involved. Yet, because only the 45S maternal-type (45S-M) rRNA is expressed during oogenesis, the zebrafish genome provides a rare opportunity to compare an ERC 45S locus to a non-ERC 45S locus.

In this study, we analyzed the genomic composition of the 45S-M and 45S-S (somatic-type) loci in combination with ultra-long read Nanopore sequencing of ERCs present in total DNA isolated from zebrafish eggs.

We discovered 45S-M flanking sequences that were absent in the 45S-S locus and showed high homology to immunoglobulin (Ig) switch regions. Also, several other unique G-quadruplex DNA containing regions were found in the 45S-M locus. Some of those auxiliary regions showed different sizes in the sequenced ERCs, although within each ERC they appear to have identical sizes. These results point to a two-step system for ERC synthesis in zebrafish oogenesis: first the 45S-M repeat is excised from the chromosome into an ERC by recombination that uses the flanking Ig switch-like regions, after which the initial ECR is multiplied and extended into many ECRs with a varying number of 45S-M repeats.

## INTRODUCTION

Ribosomes are at the core of the cellular translation system. In eukaryotes a ribosome is made up by many proteins and four non-coding ribosomal RNAs (rRNAs) of different length and function: 5S, 5.8S, 18S, and 28S rRNA. rRNAs function as the scaffold for the ribosomal proteins, but more importantly they carry out the essential peptide bond formation (Nissen *et al*. 2000), which hints to the possibility that the early-evolution ribosome consisted only of rRNAs (Moore *et al*. 2011, Noller *et al*. 2012). Whereas the 5S rDNA is present in the genome as many separate genes in tandem repeats, 18S, 5.8S and 28.S rDNAs are organized in a repeat unit that is transcribed as one continuous transcript (named 45S) and is typically present in the genome as tandem repeats.

Over the last years, we have reported on the discovery of an intriguing dual translation system in zebrafish development (Locati *et al*. 2017a, 2017b and 2018; Pagano *et al*. 2019a and 2019b; Breit *et al*. 2019). The two systems encompass at least all essential non-coding RNA containing components of the translation process: the spliceosomes (snRNA) (Pagano *et al*. 2019b); the ribosomes (5S, 5.8S, 18S, and 28S rRNA) (Locati *et al*. 2017a, 2017b and 2018) plus the pre-rRNA processing snoRNAs (Pagano *et al*. 2019a); the tRNAs (Breit *et al*. 2019) plus the pre-tRNA processing RNaseP (Breit *et al*. 2019); and the Signal Recognition Particle (SRP-RNA) for translocation of the translational complex to the ER (Breit *et al*. 2019). Of this dual translation system, the maternal-type variant (45S-M) is produced during oogenesis and progressively replaced by the somatic-type variant (45S-S) during embryogenesis (Locati *et al*. 2017b, Ortega-Recalde *et al*. 2019).

Zebrafish, much like Xenopus, are thought to amplify their 45S rDNA via so-called extrachromosomal circular DNA (eccDNA) in order to produce sufficient 45S rRNA (Brown & Dawid 1968, Gall 1968, Watson-Coggings and Gall 1972, Thiry & Poncin 2005). Thus far however, little is known about the mechanisms that are involved in the generation of these extrachromosomal rDNA circles (ERCs) other than that in *Xenopus* a rolling circle mechanism appears to be involved (Crippa and Tocchini-Valentini 1971, Hourcade *et al*. 1973, Rochaix *et al*. 1974). Yet, because in zebrafish oogenesis only the 45S-M locus is transcribed (Locati *et al*. 1917b), it is evident that the ERCs are formed from this locus exclusively. Hence, the zebrafish provides an excellent opportunity to analyze the ERC formation process, as the ERC-forming 45S-M locus can be compared to the non-ERC-forming 45S-S locus.

The findings from our genome analyses and ultra-long read Nanopore sequencing point to a two-step ERC production process that starts with excision of an initial ERC by recombination using immunoglobulin switch-like regions flanking the 45S-M locus, followed by circle multiplication, as well as enlargement to achieve sufficient copies of the 45S-M locus. Eventually, this will result in many ERCs that thus can contain several, identical 45S-M repeats.

## RESULTS

### Genomic organization of the 45S-M locus

In zebrafish, the maternal-type 45S locus (or nucleolus organizer region, NOR) is situated close (0.5 Mb) to the 3’ end of chromosome 4, whereas the somatic-type 45S locus is near (0.8 Mb) the 5’ end of chromosome 5 (supplemental File SF3) (Howe *et al*. 2013). Both loci are relatively small in the genome reference sequence (Howe et al. 2013). However, given that these genomic sequences represent ~11.7 kb repeating sequences, it is uncertain how many 45S repeats occur in each locus in the zebrafish genomic DNA. Hence, it could mean that the small 45S loci in reality represent up-to-several hundreds of 45S tandem repeats in the genome (Sakai et al. 1995; McStay 2016).

The maternal-type 45S locus (45S-M) genome reference sequence (GRCz11) contains one complete 45S repeat that extends into another repeat, which abruptly and conspicuously ends at an *Eco*RI restriction site, after which a gap is present in the genome assembly (Figure 1A, Supplemental Files SF1 and SF3).

From our Nanopore ultra-long sequencing experiment of zebrafish DNA, we obtained a ~31.5 kb read spanning the 45S-M locus and, as it contained genomic sequences from up to ~10.2 kb upstream (Figure 1B, Supplemental File SF2), we concluded that this was a read with chromosomal sequences. In this read, the 45S-M genome locus ends 335 bp earlier than in the reference genome (i.e. inside the partial 18S sequence) and continues, after a short intermediate sequence, with an R2Dr retrotransposon sequence (Kojima and Fujiwara 2004) until the end of the read. This is in line with the observations of R2Dr retrotransposon being inserted in the 45S locus (Kojima and Fujiwara 2005) and supported by our finding that all our Nanopore long reads with R2Dr sequences, show integration in 18S or 28S rDNA (results not shown). Unfortunately, we were unable to detect any other read with chromosomal 45S-M sequences that extended beyond the R2Dr sequences. Still, we found many more Nanopore reads with 45S-M sequences that contained up to five 45S-M repeats, yet none with any 45S-M upstream sequences nor R2Dr sequences: hence these are likely ERC sequences. Since no R2Dr sequences were found in these ERC-derived reads, this points towards the possibility that only one complete copy of the 45S-M repeat is present in the zebrafish genome.

**Figure 1.**
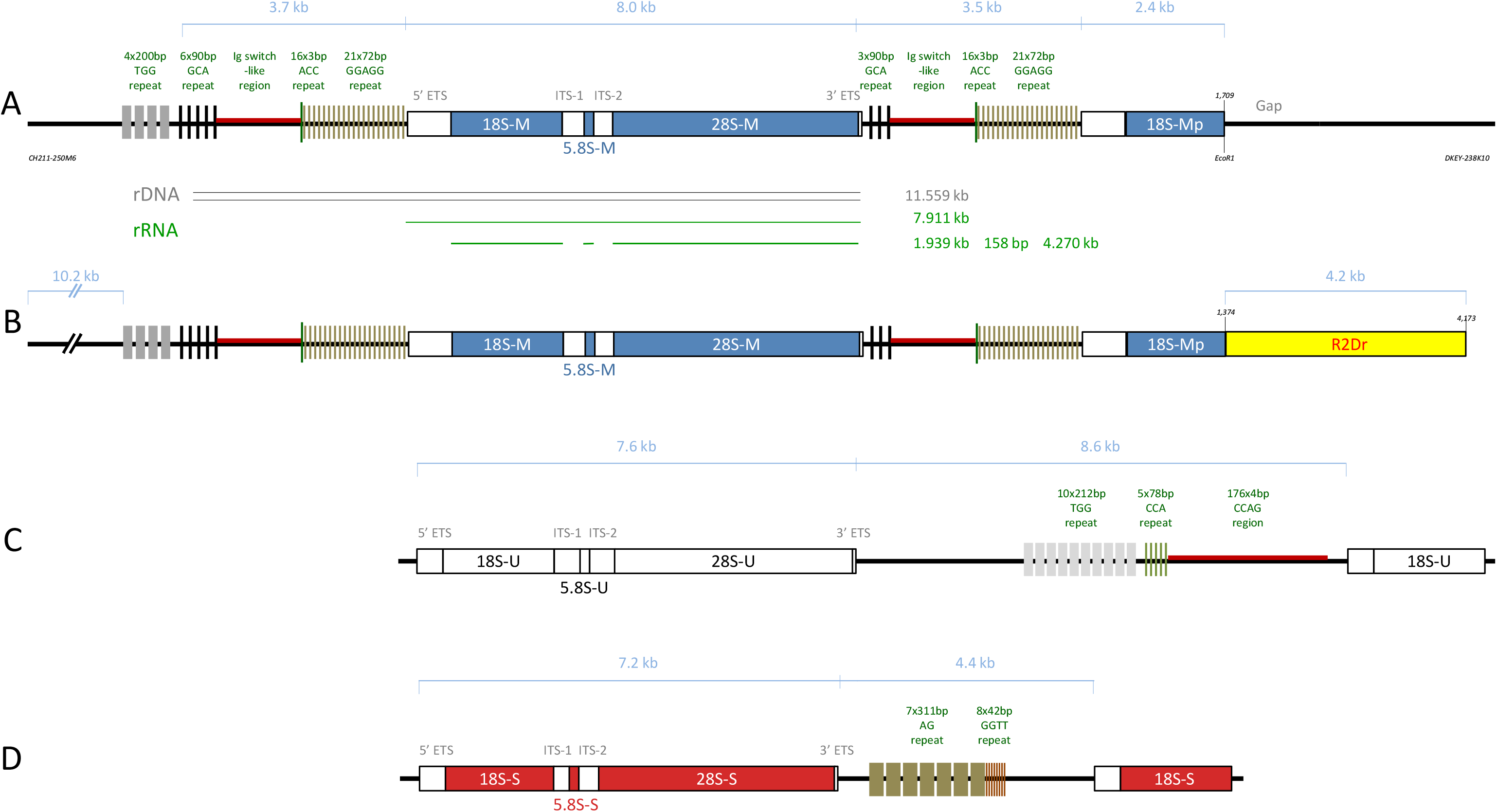
Schematic representations of the three types of 45S repeat regions in the zebrafish genome. Schematic representations of the representative parts of: **A** - the maternal-type 45S locus in the zebrafish genome (blue); **B** - a ~31.5 kb Nanopore sequencing read (blue) including a part of the R2Dr retrotransposon (yellow); **C** - the unexpressed-type 45S gene (white); and **D** - the somatic-type 45S gene (red). For the maternal-type 45S locus the circle rDNA is indicated (double black line) as well as the resulting (pre-)rRNAs (green lines). For details on the genomic locations and repeat sequences see Table 1, plus Supplemental Files SF1 and SF2.

### Distinct auxiliary genomic elements in the 45S-M locus

Comprehensive examination of the 45S-M locus revealed, next to the obvious rDNA parts, several noticeable genomic elements as characterized by their distinct sequences. The inter 45S-M repeat sequence, which is also present in front of the first 45S-M repeat (Figure 1A), contains four clear genomic auxiliary regions (Figure 1A, and Supplemental File SF4):

i. 5’-upstream region GCA-repeat (45S-R1_5’-UR-CGA-R), a 90 nt repeat with a noticeable high GCA trimer presence that sequentially occurs three or six times. The sequence is rather unique to this 45S-M locus as standard BLASTN showed no other genomic occurrence in zebrafish or any other organism in Genbank;
ii. 5’-upstream region Ig switch-like region (45S-M-R1_5’-UR-IgSL-R), a ~1,100 bp region almost entirely made up by CCAG tetramers (Supplemental File SF4). The sequence shows a significant homology (>70%) to (human) immunoglobulin switch regions;
iii. 5’-upstream region ACC-repeat (45S-R1_5’-UR-ACC-R), a 48 bp region consisting of 16 sequential ACC trimers;
iv. 5’-upstream region GGAGG-repeat (45S-R1_5’-UR-GGAGG-R), a 72 nt repeat with a two GGAGG pentamers and a noticeable high of GGG trimers presence that sequentially occurs 21 times. The sequence is rather unique to this 45S-M locus as standard BLASTN showed no other genomic occurrence in zebrafish or any other organism in Genbank.

The most noticeable characteristic is that, besides the relatively small ACC repeat, all other auxiliary regions show various degrees of G-quadruplex DNA (Parkinson 2006), which can lead to intra-molecular secondary structures: the 5’upstream region GCA-repeat contains up to 5 times the well-known TGGGGT sequence (Supplemental File SF4) (Parkinson 2006); the 5’-upstream region Ig switch-like region is similar to the quadruplex-recognized Ig switch regions (Maizels 2006); and the 5’-upstream region GGAGG-repeat contains 41 times the GGAGG structures, 18 times the G_3+_N_4_G_3+_N_3_G_3+_N_4_G_3+_N_3_G_3_ and 20 times the G_3+_N_3_G_3+_N_3_G_3_ putative quadruplex sequences (PQSs) (Huppert 2006), as well as 23 times the vertebrate telomeric G-quadruples sequence TTAGGG (albeit not concatenated) (Supplemental File SF4) (Maizels 2006). Given this overwhelming presence of G-quadruplex sequences, it is to be expected that they play an important role in the formation and use of 45S repeats. This is in line with the fact that G-quadruplex DNA formation occurs on the non-template strand to facilitate rapid 45S rDNA transcription (Maizels 2006).

All these regions are connected by relatively small intermediate sequences (Supplemental File SF3) that, even though they contain no sequences that obviously stand out, could be important for the regulation of the 45S-M locus.

The 45S-U and 45-S loci also have surrounding regions with repetitive sequences (Figure 1B and 1C, Supplemental Files SF3 and SF4). Of these regions only the Ig switch-like region present near the 45S-U locus is similar to that observed in the 45S-M locus. All other 45S-U and 45S-S regions have their own, unique patterns of repeating sequences.

### 45S-M ERC circle formation

During zebrafish oogenesis, amplification of the 45S repeat region DNA occurs by synthesis of extrachromosomal circular DNA (eccDNA) (Thiry & Poncin 2005). From our previous findings that in zebrafish egg only maternal-type rRNA is present (Locati *et al*. 2017b), we inferred that these extrachromosomal rDNA circles (ERC) on zebrafish oocytes originate exclusively from the 45S-M locus. Presently it is unknown how ERCs are produced (Mansisidor *et al*. 2018), however one of the discovered auxiliary genomic elements in the 45S-M locus shows significant homology to specific immunoglobulin switch regions (Bhattacharya *et al*. 2010). For instance, compared to humans, the zebrafish 45S-M Ig switch-like regions have a homology to the switch regions of IgM (~71%), IgA1 (~75%), IgA2 (~77%) and IgE (~71%, just for 260 bp). Given that, similar to the Ig constant exons, the Ig switch-like regions surround the 45S-M rRNA coding sequences (Figure 2A), it seems quite likely that a process using recombination, similar to Ig class switch recombination (CSR) could produce an ERC containing a 45S-M repeat (Figure 2b and 2C). Given the need of ERC during early meiosis this may actually occur during synapsis. Whereas in Ig CSR, the remaining chromosomal recombination product is functional and the Ig-switch circle is discarded, in ERC formation recombination, the circle is functional and the remaining chromosome will remain without a 45S-M locus (Figure 2).

**Figure 2.**
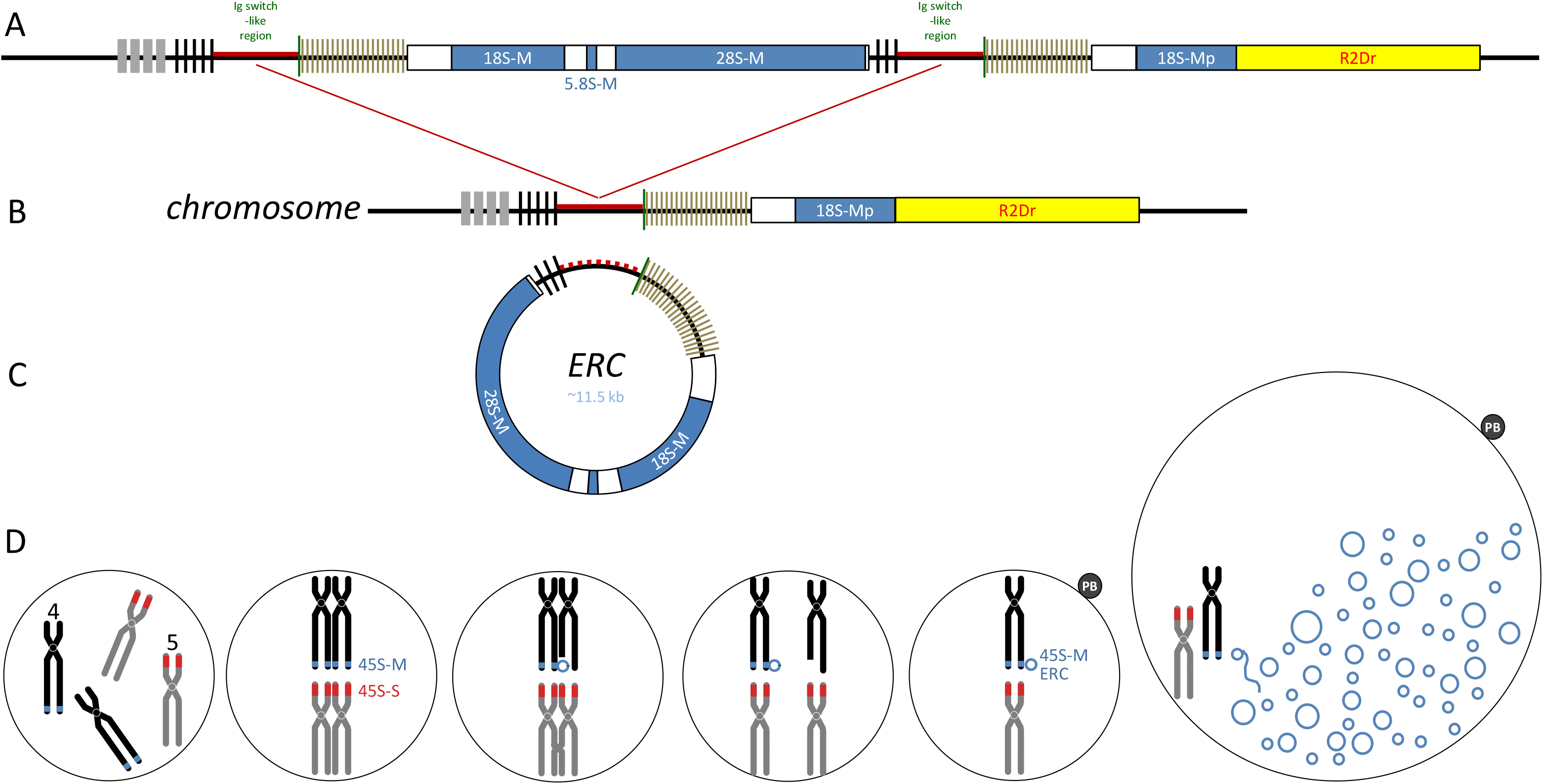
Schematic representations of the assembly and production of rDNA circles in zebrafish oogenesis. Schematic representations of **A** - representative parts of the maternal-type 45S locus in the zebrafish genome (blue) plus the recombination between the Ig switchlike regions (red dashed lines); **B** - the remaining part of maternal-type 45S locus in chromosome 4 after recombination; **C** - the extrachromosomal rDNA circle (ERC) containing a complete maternal type ?5S repeat (blue); and **D** - a hypothetical scheme how during meiosis-I, maternal type recombination might produce an initial 45S-M ERC followed by rolling circle replication resulting in many ERCs in the eventual oocyte. For details confer to the main text.

As second step in the suggested two-step process of ERC formation in zebrafish oocytes, the 45S-M ERC is formed it may be amplified by a rolling-circle process (Figure 2D) (Crippa and Tocchini-Valentini 1971, Hourcade *et al*. 1973, Rochaix *et al*. 1974). This could result in ERCs that contain one to several copies of the 45S-M repeat. In the longest ERC sequencing read (50.3 kb, Supplemental File SF5) we observed parts of five consecutive 45S-M repeats (Supplemental File SF6), but there obviously could be bigger 45S-M ERCs.

To investigate the ERCs for clues about their construction, we analyzed them by ultra-long read Nanopore sequencing of DNA from a large pool of unfertilized zebrafish eggs. This resulted in 125 reads with at least two unique 45S-M elements (Supplemental Files SF6 and SF7). From the analysis of these reads, we were unable to determine an apparent linking sequence where the ERC closes in any of the reads, which was not unexpected given the challenges that on the on hand perfect recombination between virtually identical genomic elements would leave no recognizable mark in the genome and on the other hand the average 10%-15% error rate of Nanopore sequencing, which renders small differences hard to trace. However, after annotating the Nanopore reads for the presence of 45S-M parts with a special attention to the 45S upstream elements, we noticed that several elements occurred with varying sizes. For instance, the 21×72bp GGAGG repeat (Figure 1A), which has an average size of 1,503 in the reference genome, was found in three different size ranges, with the largest about 900 bp longer (Figure 3B and 3D). Similarly, the stretch of ACC trimers (Figure 1A), ranged from 21 bp (7x) to 48 bp (16x) and the Ig switch-like region (Figure 1A) ranged from 1,049 bp to 3,437 bp with an average size in the reference genome of 1.135 bp (Figure 3A, SC, 3D and Supplemental File SF6). These differences compared to the genomic reference sequence, likely are a result from imprecise recombination events during the ERC formation process.

**Figure 3.**
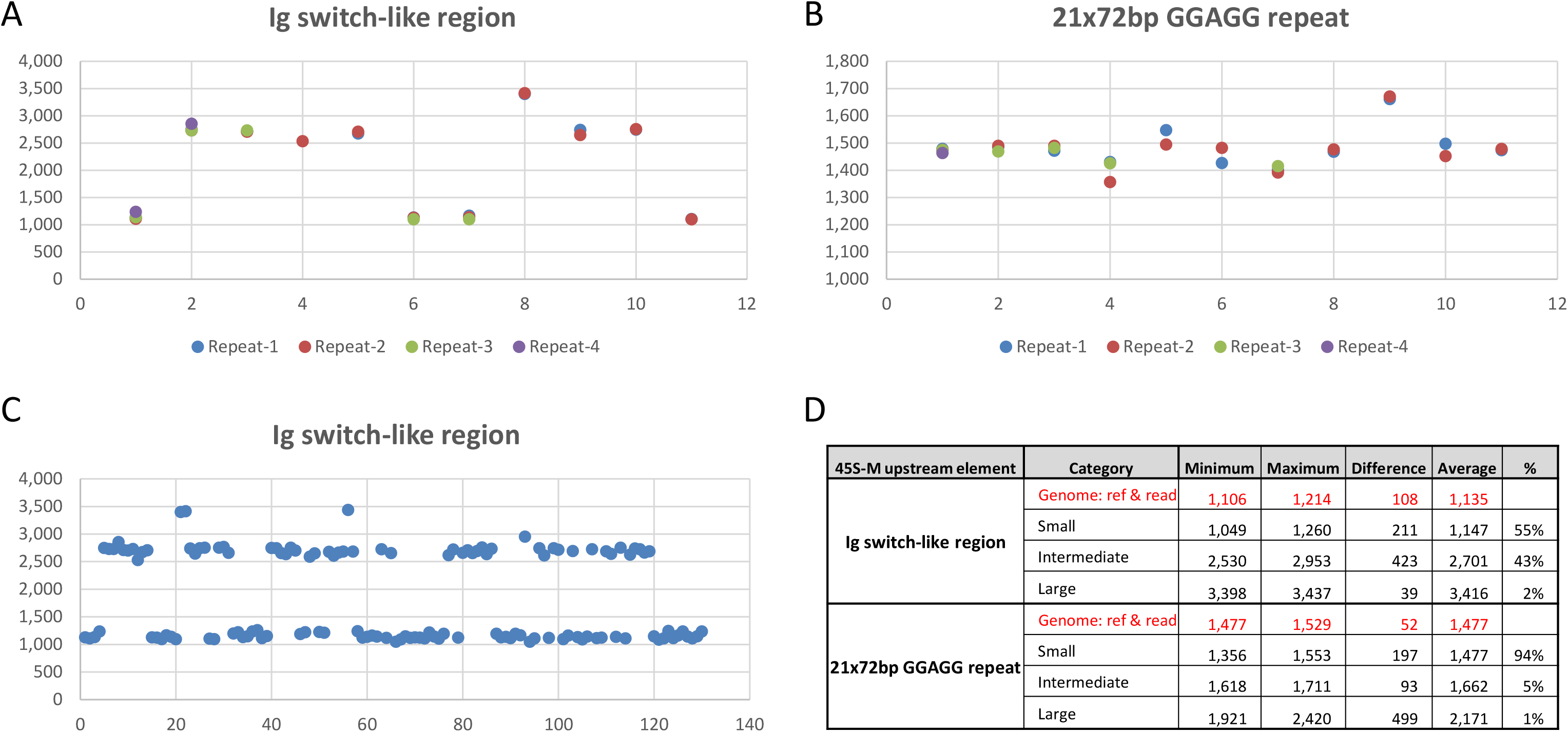
Sizes of auxiliary genome elements of the 45S-M locus in ERC. Dot plot presenting the sizes of two auxiliary genome elements of the 45S-M locus in ERC as determined by Nanopore sequencing (Supplemental File SF6). **A** - the sizes of the Ig switch-like regions in the 11 longest sequencing reads; B - the sizes of the 21×72bp GGAGG repeats in the 11 longest sequencing reads; **C** - the sizes of all Ig switch-like regions in all 125 sequencing reads that contain at least two 45S-M elements; and **D** – the summary of the sizes of all Ig switch-like regions and 21x72bp GGAGG repeats in all 125 sequencing reads that contain at least two 45S-M elements. See the main text for details.

An important observation in the size differences of these elements is that, although there is a substantial size range possible, the size of each element is virtually always the same for each occurrence in the same read. Hence, in every read, the representatives of each element in the successive 45S repeats have the same size (Figure 3A and 3B). This supports the proposed two-step ERC formation model.

## CONCLUSION

The crucial outcome of the here presented study is a hypothetical two-step process for the production during oogenesis of sufficient 45S rDNA copies needed in the zebrafish embryogenesis. It appears that, likely during meiotic synapsis, an initial ERC is excised out of one of the chromosomes 4 by recombination that involves flanking Ig switch-like sequences. The thus formed initial ERC is subsequently multiplied by a rolling-circle process, resulting in larger ERCs, containing one to several 45S-M repeats (up to at least 5) (Figure 2). The remaining affected chromosome 4 might be discharged of via deposition in a polar body (Figure 2).

This two-step 45S-M ERC hypothesis immediately leads to another intriguing observation in that two out of the three complete 45S-U repeats on chromosome 4 are also flanked by Ig switch-like regions almost identical to those of 45S-M (Figure 1, Supplemental Files SF3 and SF4). Thus far, we have been unable to establish where the 45S-U variant is expressed, but by analogy, it seems to be in a situation that a lot of rRNA is needed in a relative short amount of time and a 45-S-U ERC is needed to achieve that, for instance in the production of blood cells (Rochaix and Bird 1974).

A final interesting observation is the fact that the presence of a strict dual translational system thus far only is found in zebrafish where the maternal-type variant seems to employ an Ig switch-like recombination system to achieve sufficient 45S copies. Paradoxically, although zebrafish possess a V(D)J recombination system to produce variable Ig receptors, it has no Ig class-switching recombination system (Stavnezer and Amemiya 2004).

With these findings of our study into zebrafish ERC, we have presented another titillating chapter in the evolving story of the dual translation system in zebrafish development.

## Materials & Methods

### Biological materials

Adult zebrafish (strain ABTL) were handled in compliance with local animal welfare regulations and maintained according to standard protocols (http://zfin.org). The breeding of adult fish was approved by the local animal welfare committee (DEC) of the University of Leiden, the Netherlands. All protocols adhered to the international guidelines specified by the EU Animal Protection Directive 86/609/EEC. Unfertilized eggs (oocytes) were collected by squeezing the abdomen of 8 different spawning females. Eggs were pooled, snap-frozen in liquid nitrogen and stored at −80°C.

### DNA isolation and ultra-long read Nanopore sequencing

DNA was isolated from a collection of frozen egg pools as described previously (Mayonade et al., 2016), using some minor modifications of the original protocol. DNA quality was assessed by spectrophotometry using a Nanodrop 2000 (ThermoFisher Scientific) and by gel electrophoresis using Genomic DNA ScreenTapes on a 2200 TapeStation System (Agilent Technologies).

A long-read sequencing library was prepared from 430 ng of purified DNA using the SQK-RAD004 kit (Oxford Nanopore). The library was run on a MinION 9.4.1 flow cell using a MinION mkIB device (Oxford Nanopore Technologies) according to the manufacturer’s instructions.

### Bioinformatics

Base calling was performed by Guppy Version 3.2.4 requiring a minimum average read quality of 7 and a trim strategy set to ‘DNA’. Next, the Nanopore reads were mapped onto the zebrafish DNA chromosome 4: 77,555,866-77,567,422 (45S-M locus) with minimap2 version 2.12-r836-dirty using the Oxford Nanopore specific preset. The resulting mapping file was filtered using SAMtools version 1.9-74-gf69e678 and only reads that mapped were kept. A BLAST (version 2.7.1+) database was made containing the three 45S-M 5’ upstream region-intermediate sequences: 45S-M-R1_5’-UR-IS-1, 45S-M-R1_5’-UR-IS-2, and 45S-M-R1_5’-UR-IS-3 (Supplemental File SF3). The mapping reads were blasted onto this database using a word size of 13 and a filtering at an e-value of 1e-10. The result was written in a tabulated form which allowed for the determination of the relative distances of the repeat elements (Supplemental file SF3).

## SUPPLEMENTAL MATERIAL

Supplemental File SF1 genomic sequence.docx

Supplemental File SF2 31kb Nanopore read sequence.docx

Supplemental File SF3 genomic parts.xlsx

Supplemental File SF4 repeats.docx

Supplemental File SF5 longest 50 kb Nanopore read.docx

Supplemental File SF6 annotated Nanopore reads.xlsx

Due to its size, Supplemental File SF7 can be downloaded from: http://genseq-h0.science.uva.nl/projects/circular_45S/Supplemental_File_SF7_Nanopore_reads_QS-7+.fastq.gz

## ACKNOWLEDGMENTS

We acknowledge the support of The Netherlands Organization for Scientific Research (NWO) grant number 834.12.003.

## Notes

http://genseq-h0.science.uva.nl/projects/circular_45S/Supplemental_File_SF7_Nanopore_reads_QS-7+.fastq.gz

