## Supplemental File SF1 for "Immunoglobulin switch-like recombination regions implicated in the formation of extrachromosomal circular 45S rDNA involved in the maternal-specific translation system of zebrafish"

### Supplemental File SF1: 45S-M Repeat region, genomic sequence

>-3,680--3,142_5'-upstream-region_5x90-CGA-repeat_(538bp)_(45S-R1_5'-UR-CGA-R)

TGGGTAACCAGCAGCACGCAGTACCTATATGGGGTAACCAGCAGCAGAAGGGATTCAGTCCCAACCCGCACCCCACAGCAGAGCCTTTATTGGGGCACCAGCAGCACGCAGTACCTATGTGGGGTAACCAGCGGCAGAAGGGGTTCTGTCCCGA

CCCGCACCCCACAGCAAAGCCCCTACTGGGGCACCAGCAGCACGCAGTACCTATGTGGGGTAACCAGCGGCAGAAGGGGTTCTGTCCCGACCCGCACCCCACAGCAAAGCCCCTACTGGGGCACCAGCAGCACGCAGTACCCACGTGGGTCGCCAGCGGCAGAAGGGGTCCAGCCTCGCCCCGCACCCCACAGCAAAGCCTTTAGTGGGGCACCAGCAGCACGCAGTACCTACATGGGGTTACCAGCGGCAGAAGAGGTCCGGACCCGACCCACACCCCACAGCAAAGCCTTCGATGGGGTACCAGCAGCACGCAGTACATAAATGGGTCACCAGCGGCGCGTGAGTCTGGCGCGCGCGCCTTACCGGAGTCAGTGTCCCCGGG

>-3,141--3,058_5'-upstream-region_intermediate-sequence-1_(83bp)_(45S-R1_5'-UR-IS-1)

CTTAAGAGTGGGGTGCCCGGGCACGGCTGGTGAACGGGTTCCATGTGGTGGGGTCAGAGGCCGGCCGGTGTCCCCCGGCCTAG

>-3,057--1,951_5'-upstream-region_Ig_Switch-like_region_(1,106bp)_(45S-R1_5'-UR-IgSl-R)

CCAGCCAGCCAGTCAGTCAGCCCGGCCCAGCCCAGCCCAGCCCAGCCAGCCAGCCAGCCAGCCAGCCAGCCCAGCCCAGCGCTGTCCAGCCAGCCCTGTCCAGCCCAGCCCAGGCAAGCCCAGGCCGGCCGGCCAGCCAGCCAGCCAGCAAGCCCAGGCCAGGCCAGGCCCAGCCAGCCCAGCCAGCCCTGCCCTGCCCTGCCCTGCCCTGCCCAGCCCAGCCCAGGCCGGCCAGCCAGCCAGCCAGCCAGCCAGCCAGCCAGCCCAGGCCAGGCCAGGCCAGGCCAGGCCAGGCCAGGCCAGGCCAGGCCAGGCCCAGCCCGGCCAGGCCAGGCCCAGCCCAGCCAGCCCAGTCAGCCAGCCAGCCCAGCCCAGCCAGCCAGCCAGCCAGCCAGCCAGCCCAGCCCAGCCCAGCCCAGGCAAGCCCAGGCCGGCCAGCCAGCCAGCCAGCCAAGGCCAGGCCAGGCCCAGCCCAGCCAGCCAGCCAGCCAGCCCAGGCAAGGCCAGGCCAGGCCAGGCCAGGCCAGGCCAGGCCAGGCCAGGCCAGGCCAGGCCCAGCCCAGCCAGGCCAGCCCAGCCGAGCCCAGCCAGCCAGCCAGCCAGCCAGCCCAGCCCAGCCCAGCCCAGCCCTGCCCTGCCCTGCCCTGCCCTGCCCTGCCCAGCCCAGCCCAGCCCAGCCCAGCCAAGCCCAGGCCGGCCAGCCAGCCAGCCAGCCAGCCAGCCCAGGCCAGGCCAGGCCAGGCCAGGCCAGGCCAGGCCAGGCCCAGCCCAGCCAGCCAGCCAGCCAGCCAGCCAGCCAGCCAGCCCAGGCCAGGCCAGGCCAGGCCATGCCCAGCCCAGCCAGCCAGCCAGCCCAGGCCAAGCAAGCCCTGGCTGGGCAGCAAGCCCAGCTCAGCCAGCCAGCCCAGGCCAGGCAAGCCCTGGCCGGCCAGCCAGCCAAGCCCAGCCAGCCAATCAGTCAGTCAAGCCAAGGCCAGTCAGTCAGGTCAATCAGCCCAGCCCAGCCCAGCCTAGCCTAGCCTAGCCCAGCCCAGCCCAGCCCAGCCCAGCCCAGCCCTGCCCTGCCCAACCTGACAGCCAGGCCTGCTCAGCCAGCCAGCCAGCCA

>-1,950--1,851_5'-upstream-region_intermediate-sequence-2_(99bp)_(45S-R1_5'-UR-IS-2)

TCCTGGTGTTGGAGAGGCTTGGGCTTTAGAGTGGTTGGTCGATAGTCCATGGCCGGAGGGAGAGTGAGCACTATCTCCTCCCACTTCTCCTCTTCCTTT

>-1,850--1,802_5'-upstream-region_16x3-ACC-repeat_(48bp)_(45S-R1_5'-UR-ACC-R)

ACCACCACCACCACCACCACCACCACCACCACCACCACCACCACCACCC

>-1,801--298_5'-upstream-region_21x72-GGAGG-repeat_(1,503bp)_(45S-R1_5'-UR-GGAGG-R)

CCGGAGGCCCGGGGATTGGGCTTGGGGATAGGGTAAGGGTTGCCTGGAGGCTTGGGGGTGGGTTAGGGCTGCCCGGAGGCCCGGGGATTGGGCTTGGGGATAGGGTAAGGGTTGCCTGGAGGCTTGGGGGTGGGTTAGGGCTGCCCGGAGGCCCGGGGATTGGGCTTGGGGATAGGGTAAGGGTTGCCTGGAGGCTTGGGGGTGGGTTAGGGCTGCCAGGAGGCCCGGGGATTGGGCTTGGGCATAGGGTAAGGGTTGCCTGGAGGCTTGGGGGTGGGTTAGGGCTGCCTGGAGGCCCGGGGATTGGGCTTGGGGATAGGGTAAGGGTTGCCTGGAGGCTTGGGGGTGGGTTAGGGCTGCCAGGAGGCCCGGGGATTGGGCTTGGGCATAGGGTAAGGGTTGCCTGGAGGCTTGGGGGTGGGTTAGGGCTGCCTGGAGGCCCGGGGATTGGGCTTGGGCATAGGGTAAGGGTTGCCTGGAGGCTTGGGGGTGGGTTAGGGCTGCCTGGAGGCCCGGGGATTGGGCTTGGGGATAGGGTAAGGGTTGCCTGGAGGCTTGGGGGTGGGTTAGGGCTGCCAGGAGGCCCGGGGATTGGGCTTGGGCATAGGGTAAGGGTTGCCTGGAGGCTTGGGGGTGGGTTAGGGCTGCCTGGAGGCCCGGGGATTGGGCTTGGGGATAGGGTAAGGGTTGCCTGGAGGCTTGGGGGTGGGTTAGGGCTGCCAGGAGGCCCGGGGATTGGGCTTGGGCATAGGGTAAGGGTTGCCTGGAGGCTTGGGGGTGGGTTAGGGCTGCCTGGAGGCCCGGGGATTGGGCTTGGGGATAGGGTAAGGGTTGCCTGGAGGCTTGGGGGTGGGTTAGGGCTGCCTGGAGGCCCGGGGATTGGGCTTGGGGATAGGGTAAGGGTTGCCTGGAGGCTTGGGGGTGGGTTAGGGCTGCCAGGAGGCCCGGGGATTGGGCTTGGGCATAGGGTAAGGGTTGCCTGGAGGCTTGGGGGTGGGTTAGGGCTGCCTGGAGGCCCGGGGATTGGGCTTGGGGATAGGGTAAGGGTTGCCTGGAGGCTTGGGGGTGGGTTAGGGCTGCCAGGAGGCCCGGGGATTGGGCTTGGGCATAGGGTAAGGGTTGCCTGGAGGCTTGGGGGTGGGTTAGGGCTGCCAGGAGGCCCGGGGATTGGGCTTGGGGATAGGGTAAGGGTTGCCTGGAGGCTTGGGGGTGGGTTAGGGCTGCCAGGAGGCCCGGGGATTGGGCTTGAGCATAGGGTAAGGGTTGCCTGGAGGCTTGGGGGGGTTAGGGTTAGGGTTGCCCGGAGGCCTGGGGATTGAGCTTGGGGATAGGGTAAGGGTTGCCTGGAGGCTTGGGTGTTAGGGTTAGGGTTTCCCGGAGGACTGGGAATTGGGTTAGGGTTGCCTGGAGACTTGGGGGTGGGGTTAGGGCTGCCCGGAGGCCTGGGGATTGGGCTTGGGGATAGGGTTAGGGTTGCCCGGAGGCTTAGTTGTTGGGCTTGTG

>-297-0_5'-upstream-region_intermediate-sequence-3_(297bp)_(45S-R1_5'-UR-IS-3)

GAGCTTTGGCTTGCCTGGTTGCCTGAGGCTTGGGAGAAGTGCCACCCCGCGTGCCAGCCAGAAAGGTCAGCTGTGGGTGACCAGCAGCACCGCTCCGGCTGCCGAAGGCAAAGTCAGTCGCGGGTCACCATCGGCACCCGATTCGGCAAGGGAGAAGCAAGAAGCGGCCGCTAACCCTGACTATCCATGGCCCCCGAGCTCAGGTTCACAGCCGGGCGGGACGTGACCTCTTGGGTCTCTCTCTCCCGGAGGTCTGGGCAGCCGTCACTGGTTATCGCTGACCTGCCGTCGAGGTTT

>1-761_45S-5'-external-transcribed-spacer_(761bp)_(45S-R1_5’-ETS)

CGGGGAAGGCGTGATTTAGTGGACTTAGGTTTTGAGCGCGTAACTACGTGCCACCGCCTTCCACCACCTGACCGATGGGAATTGTGCGCGATGCCCCAGGGGCAGGCCCCTTGCCCGGTCCTCCGACAAAGGGGACCGCTGTGGGTGGGGTGGGTTCCTCCCCGTTTTGGGCATTCGAAAGGAGGGCGAGGCGGGCCTTCACCCAGACCTCGACCCTTGCTCCTTTGGGTTAAAGACTCAGATGGCCCGTCGATCTCAGACCGCCCTGTACCGACGGAACAAGTGCGCTAGTCGGTGCTGTGCGTGCGCTCTTACCGGGCAGCATCCCTCCTTGCCCCCCCTCCTCTTCCTCACCACCCGGGTGTAATCCCGGGCGCGTAGGCTGACTTGGGGGTGGGGAAGCTGTGTGGTAGGCGCGCGCGGCATCTGCGGCGCACTCGATCCCGTTGCGAAGCACGATCCTTCCTGGTGATGGCCTCGACCCAGCGGCCAGCCCAACGCCGGCAGACCCCCCTCATTTCCATGTGTGCGGTGGGGGGTGGGTAGGCGGCGGTACCTGGGAGCCCAATCCCCCCCTCCCACTTAGCCCGCGTGCCCCCTCCTGCGAGCAGTCGTGACGCCTCGGCGGAGCGAGGCCCGCTTGTAGGTAGGCGGCGCGGAGCTGTGGTGCGGGCGGGAGGGTTCTCTTCCGCCGAGTCCTTTCCCTCTCTCCTCCGGCACTCTCGGGCCTTACCACCACGCGATTCACCCGAGGCGAGGGC

>762-2,700_45S-18S-coding-sequence_(1,939bp)_(45S-R1_18S)

TACCTGGTTGATCCTGCCAGTAATATATGCTTGTCTCAAAGATTAAGCCATGCAAGTCTAAGTGCACACGGCCGGTACAGTGAAACTGCGAATGGCTCATTAAATCAGTTATGGTTCCTTTGATCGCTCCACCCGGTTACTTGGATAACTGTGGCAATTCCAGAGCTAATACATGCCAACGAGCGCCGACCTGGGCCGCCCTTCTCCCCTCGGGGCGGGGGGTGGGTACCCGGGGACGCGTGCATTTATCAGATCCAAAACCCATGCGGGTGCGCGGGCGGTGGAGAGGGGGGCCTCGCGCCTACCCGCCGCCGTCCGCCCCCGGCCTCGCTTTGGTGACTCTAGATAACCTCGGGCCGATCGCGCGCCCTCGCGGCGGCGACGGTTCATTCGAATGTCTGCCCTATCAACTTTCGATGGTAGGTCCGTCGCCTACCATGGTGACCACGGGTGACGGGGAATCAGGGTTCGATTCCGGAGAGGGAGCCTGAGAAACGGCTACCACATCCAAGGAAGGCAGCAGGCGCGCAAATTACCCATTTCCGACACGGAGAGGTAGTGACGAAAAATAACAATGCAGGTCTCTTTCGAGGCCCTGCAATTGGAATGAGTGCATCCCAAACCCATGGGCGAGGACCCATTGGAGGGCAAGTCTGGTGCCAGCAGCCGCGGTAATTCCAGCTCCAATAGCGTATGCTAACGTTGCTGCAGTTAAAAAGCTCGTAGTTGGATCTCGGGGACCGGGCCGCGCGGTCCGCCGCGAGGCGAGCCACCGCCGGTCCCGGACCCCCAGGCCTCCCGGCGCCCCCCGGATGCCCTTGACTGGGTGTCCTCGGCTTGGGGCCCGGAGCGTTTACTTTGAAAAAATTAGAGTGTTCAAGGCAGGGCCGGCACCGCGCCCCATTGAATACCCCAGCTAGGAATAATGGAATAGGACCCCGGTTCTATTTTCTGTGGGTTTCCGGAACCCGGGGCCATGATCGAGAGGGACGGCCGGGGGCATTCGTATTGCGCCGCTAGAGGTGAAATTCTTGGACCGGCGCAAGACGGACCGGAGCGAAAGCGTTTGCCAAGAACGTTTTCATTAATCAAGAACGAAAGTCGGAGGTTCGAAGACGATCAGATACCGTCGTAGTTCCGACCGTAAACGATGCCGACCCGCGATCCGGCGGCGTTTATTCCCATGACCCGCCGGGCAGCGTTGCGGGAAACCACGAGTCTCTGGGCTCCGGGGGGAGTATGGTTGCAAAGCTGAAACTTAAAGGAATTGACGGAAGGGCACCACCAGGAGTGGAGCCTGCGGCTTAATTTGACTCAACACGGGGAACCTCACCCGGCCCGGACACGGAAAGGATTGACAGATTGACGGCTCTTTCTCGATTCTGTGGGTGGTGGTGCATGGCCGTTCGTAGTTGGTGGAGCGATTTGTCTGGTTGATTCCGATAACGAACGAGACTCTGGCATGCTAACTAGTTACGCGGCCCCGCGCGGTCGGCGTCTGCAACTTCTTAGAGGGACAAGTGGCGTTCAGCCACGCGAGACTGAGCAATAACAGGTCTGTGATGCCCTTAGATGTCCGGGGCTGCACGCGCGCCACAATGGGCGGATCAACGTGTGCCTACCCTGCGCCGACAGGCGCGGGTAACCCGTTGAACCCCGCCCGTGATGGGGACCGGGGATTGAAACTATTTCCCGAGAACGAGGAATTCCCAGTAAGCGCAGGTCATCAGCTTGCGTTGATTAAGTCCCTGCCCTTTGTACACACCGCCCGTCGCTACTACCGATTGAGCGGCTCAGTGAGGTCCTCGGATCGGCCCCGCCCGGGGCTCCCTTACCGGGGGCCCTGGTGGAGCGCCGAGAAGACGATCGAACTCGGTCGTTTAGAGGAAGTAAAAGTCGTAACAAGGTTTCCGTAGGTGAACCTGCGGAAGGATCATTA

>2,701-3,092_45S-internal-transcribed-spacer-1_(392bp)_(45S-R1_ITS-1)

ACGGGGTCGAGGGGATCTCCTCCTCACGCCCAGAGGGCGAAGCCACGGTAAGCTTCCGCGCGGTGCGGGAAGTCCCTACGGGTCTACCTCCCCACCATCGCGCGCGCGAGCGTGATCGGTGATCCAAAGGTTGGCGTCGCGGGCTCCCGGCGGGTACCCGGTTGGTCTCGACCACCCTCGACCTGTCCCCTCTGCGGGGGGGAGAGCGACGGAGGTGCGTGGGGGCCGTGGGTTTAAAAGCACTCTTCGCGTTTCCCCACCCGGGGGGGAAGCAGAGAGGAGACGCCCGTCCCGGGGCCCTGCCGGCCGATGTTTAATTTCCCCACCCCCCCCTCCCGAAGCGTCCTCTGTCTCGGACCGTAACGATTGAAAGAACGAAAACAAGAGTGTAC

>3,093-3,250_45S-5.8S-coding-sequence_(158bp)_(45S-R1_5.8S)

AACTCTTAGCGGTGGATCACTCGGCTCGTGCGTCGATGAAGAACGCAGCTAGCTGCGAGAACTAATGTGAATTGCAGGACACACATTGATCATCGACCTTTCGAACGCACATTGCGGCCCCGGGTCCATCCCGGGGCCACGCCTGTCTGAGGGTCGCC

>3,251-3,580_45S-internal-transcribed-spacer-2_(330bp)_(45S-R1_ITS-2)

TTGCTATCGATCGGACGGGGGAAGAGTCGGTCTTCGTGCCGCCCTGCCCCCTGTCCGCGGCTGGAGCGTCGCAGACCCTGCCCTCCCGCGGTGGCCTACGTCCTCCCAAGTGCAGACCGCCGAACCGTTCGTCCGCCCGCTTGGGGGGCGGCTCCCATTCTCTCCCCCTCGCGCGGCTGCCGGCGGTCTAACAGCTGCCCGCGCGCGGTGGACGGGAGCTGCAGCAAACTCCCCTCGCGTTCCGAGACGACGACAGCGCGATCGGTCGACGCGAGACCGGCGTCCCGCCTCGTGTGGGACGACCGGCCGCCACCACCCGCCTGTTGGCCC

>3,581-7,850_45S-28S-coding-sequence_(4,270bp)_(45S-R1_28S)

ACGACCTCAGCTCAGACGAGAAGACCCGCTGAATTTAAGCATATTACTAAGCGGAGGAAAAGAAACCAACCGGGATTCCCCCAGTAGCGGCGAGCGAAGAGGGAAAAGTCCAGCGCCGAATCCCCGCCCCTCTGCCGAGGGCGAGGGACCTGTGGCGTACGGAGGGCCGCCTCTCTCGGCGCGGGCCGGGGGGCCAAAGTCCTTCTGATGGAGGCTTAGCCCGCGGACGGTGTGAGGCCGGTGTCGGCCCCCGCCCCGCCGGGGTGCGGTTCCTCCCGGAGTCGGGTTGTTTGGGAATGCAGCCCAAAGCGGGTGGTAAACTCCATCTAAGGCTAAATACCGGCACGAGACCGATAGCGGACAAGTACCGTGAGGGAAAGTTGAAAAGAACTTTGAAGAGAGAGTTCAACAGGGCGTGAAACCGTTAAGAGGTAAACGGGTGGGGACCGCACCGTCCGCCCGGTGGATTCAGCCCGGCGGGGCGGGGTCGGCCCGTCCGGTGCGCGCTCCCTTCGCTCCCTCATTCCTGGGGGTGGCGCGGGGGGTTGACGCCCGGGCGAAGGCTCGGCCGCCGCCGGGTGCATTTCCGCCGCGGTGGAGCGCCGCGACCGGCTCCGGTTCGGCTTGGAAGGGTCAGGGGGCGAAGGTGGCCCGTCGGTTCAGGCCGTCGGGCTTTACAGCGCCCTCCCGCCCCGACTTCGCCGCTTGCTCTCCGGGGCCGCGGGTGAGTGTCCTCCGCGCCCTCTCTGCCCTCCCTCCCTGTGGGGGGAGGTGCGGGGACGGGGTCCCCCGCCCCCGGCGTGGCGCGACAGGGGTGGACTGTCCTCAGTCCGCCCACGGCTGCGCCGCGCCGCCCAGGGCGGGGATCCGACCCACGTTCGGGCGCCCGAGGTCCGCGGCGACGCCGGCCTCCCACCCGACCCGTCTTGAAACACGGACCAAGGAGTCCAACGCGCGCGCGAGTCAGAGGGTGGCTCGCGAGCCCCTGCGGCGCAATGAAGGTGAGAACGGGGGTCTCCCCCGGGTGGGATCCCCCCGCCCCGGCGGGGGGCGCACCACCGGCCCGCCTCGCGACCCTCCGGGGGGCAGGTGGAGTTGGAGCGCGCGCGATGGCACCCGAAAGATGGTGAACTATGCCTGGGCAGGGCGAAGCCAGGGGAAACTCTGGTGGAGGCCCGCCGCGGTCCTGACGTGCAAATCGGTCGTCCGACCTGGGCATAGGGGCGAAAGACTAATCGAACCATCTAGTAGCTGGTTCCCTCCGAAGTTTCCCTCAGGATAGCTGGCGCTCGCCGATCAAGCAGTTTTATCCGGTAAAGCCAATGACTAGAGGCCTTGGGGCCGAAACGGCCTCAACCTATTCTCAAACTTTAAATGGGTAAGAGGCCCGGCTCGCTGGCGCTGGAGCCGGGCGTGGAATGCGACGCGCCTAGTGGGCCATTTTTGGTAAGCAGAACTGGTGCTGCGGGATGAACCGAACGCCGGGTTAAGGCGCCCGATGCCGACGCTCATCAGACCCCATAAAAGGTGTTGGTTGATATAGACAGCAGGACGGTGGCCATGGAAGTCGGCACCCGCCAAGGAGTGTGTAACAACTCACCTGCCGAATCAACTAGCCCTGAAAATGGATGGCGCTGGAGCGTCGGGCCCATACCCGGCCGTCGACGGCACAAGGGACACGCGAGCGGTCGCGCGCGCTGCAAGCCTCGACGAGTAGGAGGGCCGCCGCGGTGGCGCCGAAGCCCAGGGCGCGGGCCCGGGTGGAGCCGCCGCGGGCGCAGATCTTGGTGGTAGTAGCAAATATTCAAACGAGAGCTTTGAAGGCCGAAGTGGAGAAGGGTTCCATGTGAACAGCAGTTGAACATGGGTGAGCCGGTCCTAAGGGACGGGCCTACGCCGTTCGGAGGGGAGGGGCGATGGCCTCTGTCGCCCCCGCTCGACCGAAAGGGAGTCGGGTCCAGATCCCCGAGCCCGGAGCGGCGGAGACGGGCGCCGCGAGGCGCCCAGTGCGGTGACGCAAACGAACCCGGAGATGCCGGCGGGTGCCCCGGGAAAGAGTTCTCTTTTCTTTGTGAAGGGCAGGGCGCCCTGGAACGGGTTCGCCCCGAGAGAGGGGCCCGCGCCCTGGAAAGCGCCGCGCTTCTGGCGGCGTCCGGTGAGCTCTCGTCGGCCCTTGAAAATCCGGGGGAGAAGGTGTAAATCTCGCGCCGGGCCGTACCCATATCCGCAGCAGGTCTCCAAGGTGAACAGCCTCTGGCGTGTTGGAACAAGGCAGAGTAAGGGAAGTCGGCAAGTCAGATCCGTAACTTCGGGATAAGGATTGGCTCTAAGGGCTGAGCCGGTCGGGCTGAGGTGCGAAGCGGGCCTGGGCCCGAGCCGCGACTGGGGGAGCGGCCGCCCCGAGGTGCCCTGACCCCCGTTCCCGAGCCGCGGGGGTGCGCGCGCGGTGGCGCGCGGTCCCTCTCTCCCCCCGCCGCGTCCGCCTCGGGCCTTTTTGCGGTCTCGACTCCCCCCGACTCTTCCCCTCCGCCCTTTCCCCGTCCGCGGGGGTCGGGCGGGGGGGTCTCGGAGGAGGGGGGGTGGAGAGGCCGTGGGGAGGAAAGGGGTGGTTCGTGGCGTCGGGGGGCAACGGGAGGGTCCTCGCCTGCCGCGCGATGCCTCCGCTGGCCGGGGCCCGCGGGGGGCGGGATGCGCAGCGGTTGGCGGCGGCGACCCTGGGCGCGCGCCGCGCCCTTCCCGCGGATCTCCGCAGCTACGGCCCCGCGCCGGGGCCCGCGTCCGCGCGTGCGCCCCCCTCCGGGGGAGTGGCGCGCGGCCTCGGCTCCCCCCGGTGCGGGCGCCTCGGCCGGCGGCTAGCAGCCAGCTTAGAACTGTCGCGGACCAGGGGAATCCGACTGTTTAATTAAAACAAAGCATCGCGAAGGCCCTCGGCGGGTGTTGACGCGATGTGATTTCTGCCCAGTGCTCTGAATGTCAAAGTGAAGAAATTCAACGAAGCGCGGGTAAACGGCGGGAGTAACTATGACTCTCTTAAGGTAGCCAAATGCCTCGTCATCTAATTAGTGACGCGCATGAATGGATGAACGAGATTCCCACTGTCCCTACCTGCTATCTAGCGAAACCACAGCCAAGGGAACGGGCTTGGCAGAATCAGCGGGGAAAGAAGACCCTGTTGAGCTTGACTCTAGTCTGGCCCTGTGAAGAGACATGAGGGGTGTAGAATAAGTGGGAGGCCCCCGGGCTTCCCGGGCCGGCCGCCGGTGAAATACCACTACTCTTATCGTTTCCTCACTTACCCGGTGAGGCGGGGAAGCCGGGCGTCCCCCGGCGGGGGCCGCCCCTGCTTCTGGCGTCAAGCGCCCCGGGGCCGTGCGGGGGGTGGGGGCAACCCCCCTTCCCCTCCCGGCCGCCGACCGGGGCGCGACCCGCCCCGGGGACAGCGTCAGGTGGGGAGTTTGACTGGGGCGGTACACCTGTCAAACGGTAACGCAGGTGTCCTAAGGCGAGCTCAGGGGGGACAGAAACCTCCCGTAGAGCAGAAGGGCAAAAGCTCGCTTGATCTTGATTTTCAGTATGAGTACGGACCGCGAAAGCGGGGCCTCACGATCCTTCCGGCTTTTGGGGTTTTAAGCGGGAGGTGTCAGAAAAGTTACCACAGGGATAACTGGCTTGTGGCGGCCAAGCGTTCATAGCGACGTCGCTTTTTGATCCTTCGATGTCGGCTCTTCCTATCATTGTGAAGCAGAATTCACCAAGCGTTGGATTGTTCACCCACTAACAGGGAACGTGAGCTGGGTTTAGACCGTCGTGAGACAGGTTAGTTTTACCCTACTGATGTGAGCCGTTGTTGCAATAGTAATCCCGCTCAGTACGAGAGGAACCGCGGGTTCAGACATTTGGTTCGTGCGCTTGGCTGAGGAGCCACTGGCGCGAAGCCACCATCTGCGGGATTATGACTGAACGCCTCTAAGTCAGAATCCCGCCTAAAAGCAACGATACAGCAGCGCCGTCGGATCTCCGATAGGCCCCGGGTAACCGGGAAGCCCCTCTCCGGGGGGGCCCCCGGCGCGGAGAGCCATTCGTGACAGGAACCGGGGGCGCGGCTAGAACGAGCGCCGCCCCCTCTTCCAGTGACGCACCGCATGTTTGTGGGGAACCCGGTGCTTAAATGACTCGTAGACGACCTGATTCTGGGTCGGGGTGTCGTGCGTGGCAGAGCAGCTCTGTTGCTGCGATCCATTGAAAGTCAGCCCTCGATCCAAGTTTTGTC

>7,851-7,911_45S-3'-external-transcribed-spacer_(61bp)_(45S-R1_3’-ETS)

GGGCCGGACGCAGGCGACAGGGGCCACCGGTCCGACAGACCCGCGGCGTGCGAAGGGGCAC

>7,912-8,039_45S-inter-repeat-untranscribed-spacer_(128bp)_(45S-R1-R2_IRUS)

CAGCTGCGTCGGTGAAGGCTTGCCATCCCCCCTCGCCACTTGCGCGCAGGCCGAGGCACCAGCGGCGGGAGCGTGGCGCGATCCCAATACACCTGGGGCACCAGCGGCAGAGGGAGGCCTAGCCTCGG

>8,040-8,423_5'-upstream-region_3x90-CGA-repeat_(384bp)_(45S-R2_5'-UR-CGA-R)

CCCGCACCCCACAGCAAAGCCCCTACTGGGGCACCAGCAGCACGCAGTACCTATGTGGGGTAACCAGCGGCAGAAGGGGTTCTGTCCCGACCC**TA**ACCCCACAGCAGAGCC**TT**TATTGGGGCACCAGCAGCACGCAGTACCCACGTGGGTCGCCAGCGGCAGAAGGGGTCCAGCCTCGCCCCGCACCCCACAGCAAAGCCTTTAGTGGGGCACCAGCAGCACGCAGTACCTACATGGGGTTACCAGCGGCAGAAGAGGTCCGGACCCGACCCACACCCCACA**A**CAAAGCCTTCGATGGGGTACCAGCAGCACGCAGTACATAAATGGGTCACCAGCGGCGCGTGAGTCTGGCGCGCGCGTCTTACCGAAGTCAGTGTCCCCGGG

>8,424-8,506_5'-upstream-region_intermediate-sequence-1_(83bp)_(45S-R2_5'-UR-IS-1)

CTTAAGAGTGGGGTGCCCGGGCACGGCTGGTGAACGGGTTCCATGTGGTGGGGTCAGAGGCCGGCCGGTGTCCCCCGGCCTAG

>8,507-9,604_5'-upstream-region_ Ig_Switch-like_region_(1,098bp)_(45S-R2_5'-UR-IgSL-R)

CCAGCCAGCCAGTCAGTCAGCCCGGCCCAGCCCAGCCCAGCCCAGCCAGCCAGCCAGCCAGCCAGCCAGCCCAGCCCAGCGCTGTCCAGCCAGCCCTGTCCAGCCCAGCCCAGGCAAGCCCAGGCCGGCCGGCCAGCCAGCCAGCCAGCAAGCCCAGGCCAGGCCAGGCCCAGCCAGCCCAGCCAGCCCTGCCCTGCCCTGCCCTGCCCTGCCCAGCCCAGCCCAGGCCGGCCAGCCAGCCAGCCAGCCAGCCAGCCAGCCAGCCCAGGCCAGGCCAGGCCAGGCCAGGCCAGGCCAGGCCAGGCCAGGCCAGGCCCAGCCCGGCCAGGCCAGGCCCAGCCCAGCCAGCCCAGTCAGCCAGCCAGCCCAGCCCAGCCAGCCAGCCAGCCAGCCAGCCAGCCCAGCCCAGCCCAGCCCAGGCAAGCCCAGGCCGGCCAGCCAGCCAGCCAGCCAAGGCCAGGCCAGGCCCAGCCCAGCCAGCCAGCCAGCCAGCCCAGGCAAGGCCAGGCCAGGCCAGGCCAGGCCAGGCCAGGCCAGGCCAGGCCAGGCCAGGCCCAGCCCAGCCAGGCCAGCCCAGCCGAGCCCAGCCAGCCAGCCAGCCAGCCAGCCCAGCCCAGCCCAGCCCAGCCCTGCCCTGCCCTGCCCTGCCCTGCCCTGCCCAGCCCAGCCCAGCCCAGCCCAGCCAAGCCCAGGCCGGCCAGCCAGCCAGCCAGCCAGCCAGCCCAGGCCAGGCCAGGCCAGGCCAGGCCAGGCCAGGCCAGGCCCAGCCCAGCCAGCCAGCCAGCCAGCCAGCCAGCCCAGGCCAGGCCAGGCCAGGCCATGCCCAGCCCAGCCAGCCAGCCAGCCCAGGCCAAGCAAGCCCTGGCTGGGCAGCAAGCCCAGCTCAGCCAGCCAGCCCAGGCCAGGCAAGCCCTGGCCGGCCAGCCAGCCAAGCCCAGCCAGCCAATCAGTCAGTCAAGCCAAGGCCAGTCAGTCAGGTCAATCAGCCCAGCCCAGCCCAGCCTAGCCTAGCCTAGCCCAGCCCAGCCCAGCCCAGCCCAGCCCAGCCCTGCCCTGCCCAACCTGACAGCCAGGCCTGCTCAGCCAGCCAGCCAGCCA

>9,605-9,703_5'-upstream-region_intermediate-sequence-2_(99bp)_(45S-R2_5'-UR-IS-2)

TCCTGGTGTTGGAGAGGCTTGGGCTTTAGAGTGGTTGGTCGATAGTCCATGGCCGGAGGGAGAGTGAGCACTATCTCCTCCCACTTCTCCTCTTCCTTT

>9,704-9,751_5'-upstream-region_16x3-ACC-repeat_(48bp)_(45S-R2_5'-UR-ACC-R)

ACCACCACCACCACCACCACCACCACCACCACCACCACCACCACCACC

>9,752-11,254_5'-upstream-region_21x72-GGAGG-repeat_(1,503bp)_(45S-R2_5'-UR-GGAGG-R)

CCGGAGGCCCGGGGATTGGGCTTGGGGATAGGGTAAGGGTTGCCTGGAGGCTTGGGGGTGGGTTAGGGCTGCCCGGAGGCCCGGGGATTGGGCTTGGGGATAGGGTAAGGGTTGCCTGGAGGCTTGGGGGTGGGTTAGGGCTGCCCGGAGGCCCGGGGATTGGGCTTGGGGATAGGGTAAGGGTTGCCTGGAGGCTTGGGGGTGGGTTAGGGCTGCCAGGAGGCCCGGGGATTGGGCTTGGGCATAGGGTAAGGGTTGCCTGGAGGCTTGGGGGTGGGTTAGGGCTGCCTGGAGGCCCGGGGATTGGGCTTGGGGATAGGGTAAGGGTTGCCTGGAGGCTTGGGGGTGGGTTAGGGCTGCCAGGAGGCCCGGGGATTGGGCTTGGGCATAGGGTAAGGGTTGCCTGGAGGCTTGGGGGTGGGTTAGGGCTGCCTGGAGGCCCGGGGATTGGGCTTGGGCATAGGGTAAGGGTTGCCTGGAGGCTTGGGGGTGGGTTAGGGCTGCCTGGAGGCCCGGGGATTGGGCTTGGGGATAGGGTAAGGGTTGCCTGGAGGCTTGGGGGTGGGTTAGGGCTGCCAGGAGGCCCGGGGATTGGGCTTGGGCATAGGGTAAGGGTTGCCTGGAGGCTTGGGGGTGGGTTAGGGCTGCCTGGAGGCCCGGGGATTGGGCTTGGGGATAGGGTAAGGGTTGCCTGGAGGCTTGGGGGTGGGTTAGGGCTGCCAGGAGGCCCGGGGATTGGGCTTGGGCATAGGGTAAGGGTTGCCTGGAGGCTTGGGGGTGGGTTAGGGCTGCCTGGAGGCCCGGGGATTGGGCTTGGGGATAGGGTAAGGGTTGCCTGGAGGCTTGGGGGTGGGTTAGGGCTGCCTGGAGGCCCGGGGATTGGGCTTGGGGATAGGGTAAGGGTTGCCTGGAGGCTTGGGGGTGGGTTAGGGCTGCCAGGAGGCCCGGGGATTGGGCTTGGGCATAGGGTAAGGGTTGCCTGGAGGCTTGGGGGTGGGTTAGGGCTGCCTGGAGGCCCGGGGATTGGGCTTGGGGATAGGGTAAGGGTTGCCTGGAGGCTTGGGGGTGGGTTAGGGCTGCCAGGAGGCCCGGGGATTGGGCTTGGGCATAGGGTAAGGGTTGCCTGGAGGCTTGGGGGTGGGTTAGGGCTGCCAGGAGGCCCGGGGATTGGGCTTGGGGATAGGGTAAGGGTTGCCTGGAGGCTTGGGGGTGGGTTAGGGCTGCCAGGAGGCCCGGGGATTGGGCTTGAGCATAGGGTAAGGGTTGCCTGGAGGCTTGGGGGGGTTAGGGTTAGGGTTGCCCGGAGGCCTGGGGATTGAGCTTGGGGATAGGGTAAGGGTTGCCTGGAGGCTTGGGTGTTAGGGTTAGGGTTTCCCGGAGGACTGGGAATTGGGTTAGGGTTGCCTGGAGACTTGGGGGTGGGGTTAGGGCTGCCCGGAGGCCTGGGGATTGGGCTTGGGGATAGGGTTAGGGTTGCCCGGAGGCTTAGTTGTTGGGCTTGTG

>11,255-11,551_5'-upstream-region_intermediate-sequence-3_(297bp)_(45S-R2_5'-UR-IS-3)

GAGCTTTGGCTTGCCTGGTTGCCTGAGGCTTGGGAGAAGTGCCACCCCGCGTGCCAGCCAGAAAGGTCAGCTGTGGGTGACCAGCAGCACCGCTCCGGCTGCCGAAGGCAAAGTCAGTCGCGGGTCACCATCGGCACCCGATTCGGCAAGGGAGAAGCAAGAAGCGGCCGCTAACCCTGACTATCCATGGCCCCCGAGCTCAGGTTCACAGCCGGGCGGGACGTGACCTCTTGGGTCTCTCTCTCCCGGAGGTCTGGGCAGCCGTCACTGGTTATCGCTGACCTGCCGTCGAGGTTT

>11,552-2,312_45S-5'-external-transcribed-spacer_(761bp)_(45S-R2_5’-ETS)

CGGGGAAGGCGTGATTTAGTGGACTTAGGTTTTGAGCGCGTAACTACGTGCCACCGCCTTCCACCACCTGACCGATGGGAATTGTGCGCGATGCCCCAGGGGCAGGCCCCTTGCCCGGTCCTCCGACAAAGGGGACCGCTGTGGGTGGGGTGGGTTCCTCCCCGTTTTGGGCATTCGAAAGGAGGGCGAGGCGGGCCTTCACCCAGACCTCGACCCTTGCTCCTTTGGGTTAAAGACTCAGATGGCCCGTCGATCTCAGACCGCCCTGTACCGACGGAACAAGTGCGCTAGTCGGTGCTGTGCGTGCGCTCTTACCGGGCAGCATCCCTCCTTGCCCCCCCTCCTCTTCCTCACCACCCGGGTGTAATCCCGGGCGCGTAGGCTGACTTGGGGGTGGGGAAGCTGTGTGGTAGGCGCGCGCGGCATCTGCGGCGCACTCGATCCCGTTGCGAAGCACGATCCTTCCTGGTGATGGCCTCGACCCAGCGGCCAGCCCAACGCCGGCAGACCCCCCTCATTTCCATGTGTGCGGTGGGGGGTGGGTAGGCGGCGGTACCTGGGAGCCCAATCCCCCCCTCCCACTTAGCCCGCGTGCCCCCTCCTGCGAGCAGTCGTGACGCCTCGGCGGAGCGAGGCCCGCTTGTAGGTAGGCGGCGCGGAGCTGTGGTGCGGGCGGGAGGGTTCTCTTCCGCCGAGTCCTTTCCCTCTCTCCTCCGGCACTCTCGGGCCTTACCACCACGCGATTCACCCGAGGCGAGGGC

>12,313-14,021_45S-18S-coding-sequence-(partial)_(1,709bp)_(45S-R2_18Sp)

TACCTGGTTGATCCTGCCAGTAATATATGCTTGTCTCAAAGATTAAGCCATGCAAGTCTAAGTGCACACGGCCGGTACAGTGAAACTGCGAATGGCTCATTAAATCAGTTATGGTTCCTTTGATCGCTCCACCCGGTTACTTGGATAACTGTGGCAATTCCAGAGCTAATACATGCCAACGAGCGCCGACCTGGGCCGCCCTTCTCCCCTCGGGGCGGGGGGTGGGTACCCGGGGACGCGTGCATTTATCAGATCCAAAACCCATGCGGGTGCGCGGGCGGTGGAGAGGGGGGCCTCGCGCCTACCCGCCGCCGTCCGCCCCCGGCCTCGCTTTGGTGACTCTAGATAACCTCGGGCCGATCGCGCGCCCTCGCGGCGGCGACGGTTCATTCGAATGTCTGCCCTATCAACTTTCGATGGTAGGTCCGTCGCCTACCATGGTGACCACGGGTGACGGGGAATCAGGGTTCGATTCCGGAGAGGGAGCCTGAGAAACGGCTACCACATCCAAGGAAGGCAGCAGGCGCGCAAATTACCCATTTCCGACACGGAGAGGTAGTGACGAAAAATAACAATGCAGGTCTCTTTCGAGGCCCTGCAATTGGAATGAGTGCATCCCAAACCCATGGGCGAGGACCCATTGGAGGGCAAGTCTGGTGCCAGCAGCCGCGGTAATTCCAGCTCCAATAGCGTATGCTAACGTTGCTGCAGTTAAAAAGCTCGTAGTTGGATCTCGGGGACCGGGCCGCGCGGTCCGCCGCGAGGCGAGCCACCGCCGGTCCCGGACCCCCAGGCCTCCCGGCGCCCCCCGGATGCCCTTGACTGGGTGTCCTCGGCTTGGGGCCCGGAGCGTTTACTTTGAAAAAATTAGAGTGTTCAAGGCAGGGCCGGCACCGCGCCCCATTGAATACCCCAGCTAGGAATAATGGAATAGGACCCCGGTTCTATTTTCTGTGGGTTTCCGGAACCCGGGGCCATGATCGAGAGGGACGGCCGGGGGCATTCGTATTGCGCCGCTAGAGGTGAAATTCTTGGACCGGCGCAAGACGGACCGGAGCGAAAGCGTTTGCCAAGAACGTTTTCATTAATCAAGAACGAAAGTCGGAGGTTCGAAGACGATCAGATACCGTCGTAGTTCCGACCGTAAACGATGCCGACCCGCGATCCGGCGGCGTTTATTCCCATGACCCGCCGGGCAGCGTTGCGGGAAACCACGAGTCTCTGGGCTCCGGGGGGAGTATGGTTGCAAAGCTGAAACTTAAAGGAATTGACGGAAGGGCACCACCAGGAGTGGAGCCTGCGGCTTAATTTGACTCAACACGGGGAACCTCACCCGGCCCGGACACGGAAAGGATTGACAGATTGACGGCTCTTTCTCGATTCTGTGGGTGGTGGTGCATGGCCGTTCGTAGTTGGTGGAGCGATTTGTCTGGTTGATTCCGATAACGAACGAGACTCTGGCATGCTAACTAGTTACGCGGCCCCGCGCGGTCGGCGTCTGCAACTTCTTAGAGGGACAAGTGGCGTTCAGCCACGCGAGACTGAGCAATAACAGGTCTGTGATGCCCTTAGATGTCCGGGGCTGCACGCGCGCCACAATGGGCGGATCAACGTGTGCCTACCCTGCGCCGACAGGCGCGGGTAACCCGTTGAACCCCGCCCGTGATGGGGACCGGGGATTGAAACTATTTCCCGAGAACGAG**GAATTC** *(EcoR1)*
