## Supplemental File SF2 for "Immunoglobulin switch-like recombination regions implicated in the formation of extrachromosomal circular 45S rDNA involved in the maternal-specific translation system of zebrafish"

### Supplemental File SF2: 45S-M Repeat region, genomic Nanopore read sequence

>8c84d04e-8f27-4b43-8ca4-c568310da34d_(31,519bp)

**>45S-M-Upstream_region_(10,297bp)**

CAATGCATACTTCGTTCGGTGCCCCTCGGAGAGCTGAGTGTTTAACCATTTCGCATTTATCGTGAAACGCTTTCGCGTTTTCGTGCGCCGCTAAGAAGCTTTATTAGTGCATTGCTGCTGGCGCGCAGCGAAATATAGGCAATATTTTATTACGTGCTGCTCTTACGTGCTGAAACACTGAACACTTTCCTTTACTTAAGATATAATAAGCAACATCTTTAAATAAAGGGAATCACCTATTAAGTGTCATAAAGCCTATAGGGAAAAATAATCATGTTTCAGTTCTCACGGGCATTTAGGATGGATCTTGGACAGATTCTTGTCTAGAGGATGTTATAAAAAGCATTTTATGTCGAGTTATTAACGGAGAATTTTATATATGTGGTGTTATATTATGGATGTGTCGCTCCCTGCGCTCCGGGTCCCTCCGTTGGTGGCAAGACCACTGTGGATTTCGATGCCAGTCGATTACGGCGAGTAAATGAATATGATGAATGAAAACACACAGGCCTGTGCTTTATAGACATGCGTTGTACTGCTGTACAGTAGCGTGTCATTTGTTTGTTTTGGTTGTCCTATTTTAATAGTTTCGTGATGATGTGATGGTGTGCTTTATCTGCATGATTCTCGCTGAAGTGGGCGGTTGGTGACGTTGATTCGGGGATGTTCTGGGAGCGGGGTTGCGTGACCCGCGCTGCGGTATACTAGCCCCACAGTGGAAACACCATGATTTCGGCCTCACTGCTCAACCTCCCGGGCGATTGGCCGGCTGGCTGATATAAGCCCTGGCTCGCACTGGCCTGATAGTGGAAATGCGGCTAGTGACACCCTCTGCCCAGTGCATTGATGAAAATGAACAGTTGGATGTGCCAAAACTTTCTTCAGTTAAACAAGAGAAAACTGAAGTCACTGCGTTGGGAACAGACATGAGCTTCTCAAGGTGAATGTGTACCTTGGCACTAAAGAGTCAAACAACAAAAAATAAGGTCAAGAATCTTGGTGTGACTCTGGAGTCAGATCTGAGTTTCAACAGTCATGTCAAAGCAGTCAGTAAATCAGCATACTATCATCTCAAAAACATAGCAAGAATCCAGATGCTGTGTTTCCAGTGAAGACTTAGAGAAACTTGTTGATGCTTTTATCAGCAGGGTCCAAATCCAACTCACTCTTTAAGATCTCAAAACTCAGAGCTTCTGGTAGTACCTAGAATAGCAAAAATCAAATAAAGGAGGTTGAGCCTTCTCCTTTATGGCTCCTACACTCTGGAATAGCCTTCCTGATAACGTCCGAGGCTCAGACACACTCTCCCAGTTCAAAAACTAGATTAAAGACCTATCTGTTTAGTAAAGCATACCTCAGTGCATCACCTAGCGGAGTTCCACACTGGCTTCTGCATCTTGCTTATATACACTATGAACAGCAGCTACGCTAATTATTCTCTTTATTCTCTGGCCTCACCTGGGGATACTCATCCCGAGACCCTCCAGATTATGCAGAGTCACTGATTGGATCCAAGACTATTGTGAAGAGATGATCCCAAGGTTTCCATATCCGGGACCAGGCCATCTGAGCTGCTGCTGCGCTGATGGTCATGGAGGAGTGGAGAACATGAGTCTGATTCCAGCGGCGCTCCAGGGACAGACGAGTCTTCACTGAGGCCATCTTCCAGCCTCCACCGCTGTGATTGAAGCTCTGCACAAGACTTTTGGCCAGCTTAAAAGGCTGTGCCCAAATTAAAATAGCCGTGCCCACCTGAGTCTGGTTCTCTTAAGGTTTATTTTCTTCACTCCCTATCAGGTGAAGTTTTCCCCTCTCCGCTGTCGCCATGGCTCGCATGGTTCAGGACTGGCTACGCATCGATGAATTAGCTCTTCAGTGTTTGAACTCTCAGTAATGATTAGAAATCACACTGAACTGGGCTAAACTGAACTGAACTGAACTTAAACACTAAAAGCTGAACCACACTGTTCCAGTTATTATGACTATTTCCATGTGAGCTGCTTTGACACAATCTACATTGTAAAAAAGGCTATGCAAATAAAGCTGGGTGAATTGAATTATAGTAACGGCCTTCTCACTGGCCTTCCCAAAAGACAGTCAGACGATTTGTGCAGCTCATAAGTTATTGTATAACGGTTTGTTTTGTAGAGTATCAAGATATTTAGCTTAAAGGGGCTAATAATATTGAGTTTAAGCTCTTAAGGCTCTTAATTTTATTTCTTTTTGCTATTTAGGCTTATTAGAACCTAATTAGAATAAAACCGTATCATCTTTTGATGGCAGATGCATTTTAGAGAAAATGATGTCCATAAATGTGGACCTAGTGGTCTGAATGTCCATAAAACACAATAAAGTCACCTTTGTGGAAAAGAGTGAAAAACTCTTTATTTCTGCCTTGAACCCAGCGTCTTGTCCTGACTGTTTTTATGCATTAACAACACATACACACATATTTTTGAACATGCTTTGACTATCAGTAAAAATATTTTTGCCCTTTCTGGACAGTTAAAATAGTGTTTGGGACATTTCATATGCTACAGGATAACACTGGATAACACATATGTCTGCAACAAAATATAAAACATAAGACAAAAGAGCAAAATGTGCTGTCCATGTATGTGGCCAGGAGGAAAATGTTTTAAAATTAAAACTGATTTTATTCTAGCCGATATAAACAAATAAGACTTTCTCCCAAAGAAGAAAAAAAAAAAACATTATCCGACATACTGTGAAAAATTTTCTTGCTCTGTTAAACATCATTTAGGAAATATTTAAAAAATTAAAAAAAAAAAAAATAAAAGGAGCTAATAATTCTGACTTCAACTATAAATAAATATATATATATATATATATGTGTGTGTGTGTGTGTGTCTGTTTTAAGTCATGCAATTAATTACCCAGTTACAATGTAGACTTTAAACTTTTGAGAAACACCTGATTTGCTTTAAAAAATTGATGCAGTTAGGAATAGAATTGTATTATTAGAGGAGTCATATTTATTGCATGTAGCATAATTTATTTAGATTTTGAATATTTACTATTGTAGAAAAATAAACAATCTGTAACACTGTTCTCATCTTCAAATGTATAACTACTCCTTGTTATTCTGTTCTGTGATTTATTTTTAAGATTTATATGTGAAAATATATTTTTAGCTAATAACTTGGAGTGTAGTTGGAAATAACGTGCTTCAACTTAGGTGCTCCCTGTTATATCAAATCTGCATCTTTACCATCAGGAGATAACCACCTTTGTTTCATAAATGTTTAACAAACTGTCAAGATGTGTCAACCTTTATTTTGTTTTTTTACTTTCGATGAAAACCGCAGAATGGTGGATTACATAATAAAACCATGATTAAATATGAATATAAAAAACAATTAGGCATAAACATAGACCCTAACTAATGAATAATTAAATAACAAAAAAGATCAGCTAAATATTTTTTTAACTTAGTACTTTAAATGTAATATTTGCTAAGTAAAAATAATAATAATTAATCAGTTTTATTAGGCATAATAACCCTAAACTGGGCAGTCGAATATAATTTTACTGATTAGATGTAAATTAAATGATAACATATTTATCAACACCAAATAATCTCCAATAAAAATGTAAAAACTACATAATGCAGCATTAACCTACACGTATTGTTGCCACGTTTCATTTTAGGAGAGTTAAACTCATGAGAGAACTCACCGGTATTACATTTTGACCAATAGCTGGCTATTAAACTCACCGTTGAATGAAAGTCACATTGACAGAAGTATATTCTTAGTTACCCTGTTTACAGATGCTTTTCATGAACAAAACAAATCACATATTTAAAATAAAAAACTGTGATTATAGGCGTAAGCAAGGTTTAAAAAAATATTGCGTTGACATCTATTGGTAATTGCTTATGCGAAGCTTTGAGCGTCTCTATATGACGTCTGATGCTACCTATTAGAAAGCAATAGAATTCCCTTTGCACGTCATTATAGTGACCCCAGGGGCACCTTTGGCGTGCTTATACTTTGTGGCATAAGGTGCTTCTACCATGCTTCAGATGTTGATGCCTTGGGGTAAGAATGGGTTGCATACTAGCTTAGGCTTACCACTTAACAAGCTCACTGTTGTTTGTCTACCACTTTTAAATGTTGCCTGATTAGTGGTGTTAAGTATTACAATTTTACAATAATACAATAATTTGTTAAATTTCAATATATAATTCAGAAAAATCATAGAAGCAAATCAATTTCAAATAGGAATGGGTGATTAAAATACCTGCGTATCCCTATACATGTATGGTAAAACTAAGCTCCCAAACAAGCAGTCGGCATGCGCCTTTGCGGAAAGACCTAATTAAAGATAATAAAATGATAAACTGTTAATGCTTACCTCTTAAAATGCTATGAAGGTAAATCAGTAGCAAAACTCGAGAGCTTGCAAATGTAAATGTGAGCACTTGTGATAATAATATTTGTATGTGTTGGTTGAAATGTACACTCACAGCTCTGATGTGAACAGCTGAATCCCTCGTTGCACACACACACACACACACACACACACACACACAAAACACACTTGTGTTTTGAATTGGAAGATTGAATTAGTTGTGTATTTGTAAAATGTCAATCACAAATACAAATGACAGACTTCTGCGATGTGCACCGCAGAATTTGATTGCATTTCGCTCTGCAACCACAGGTCGGCCTCACAACCTCTCAGATTGCGTCCTTCAAATACAAATCTCTTGCGCTCCTGACTTTGAGTAACCATTTCTCGCAATCTCTGCTTAGCAGTGAAGTGCGGATCGATTCTAAAAATGTCGGCATCTCCAATTCCAGCTTTGTATATTCTAAGAATGAGTATCTTATCAAAATATCGGCGCTAGCTTTCTGCAACTCTCACATGATTGCCCACTGAAGCTATGCAGGGCTGCGCCGGTCAGTACCTGGGTGGGAGACCGCATGGGAAACTAGATTGCTGCCGGAAGAGGTGTTAGTGAGGCCAGCAGGGCACTCAGCCTGTGGTCTGTGGATCCTGCCCCAGTATAGTGACGGGGACTCTATACTGCTCAGTAGAAGTGCTGCTGTCTTTCGCATGGGAGCGTTAAACCCAGGTCCTGGCTCCAGGATCGTTAAAAATCCCAGGATGTCTTTCGAAAGAGCGAGAGTTGCCCAGCATCCTGTCCAAATCCGCAATCTGGTCTCTGTCCATCGTGGCCTCCTAACCATCCCATTTCATAACTGGCTTCATCACTCTGTCTCCTCTCCAGGCTCAGCTGGTGTGTGGTGTGTGGTCTGGCGCAAAATGTCTGCCGTCGCGCGTCATCCAGGTGGATGCTGCACACTGGTGCTGGATGAGGAGATTCCCCCCATTGTGTAAAGCGCTCTGAGTGCCCAGAGAAGCGCTATATAAGTGTAAGGAATTAATAATAATTCTTCTTCTTCTTATTGTTATTATTATAAAATATCGACATTTCAATCATTTGGACCAAATATGGTTTTTTTTTGGAAGTAGAAACATAGTGGAAAGTAACCACGATTACCTAGTCTGAATAATGCCAACAGGTCTGACTACTTGTTCTTCCGCCTGCAAAGTCATGTGCGTAATATGGCACTGTACAACTGTAGACTCAGAGCACAGCTAGCCACTTAGTCAGCTGGCCTGCTTTTATGCAGAATTATGTGTGGAGCTACTTCACTTTTATCAAGTCATACTGCAAGTCAAATGATGCGGTTTATGACACACGTGTAACAAAAAACAGTGAGGTACTGTGGTAATACTAAAAACCTAACCAAACATCTACATAGAAATCACGAGACAGAATATGACAACGTCATGACTAGGCGGTCAGACGAGAGGAAATAAGAGGTACCCAGGTGCTGCTAGGCAGACAGACATCGTTTTCACAGTCCTTAGGAGTGCCTTGAGCAGGCTCATCCAGGTACAGGGCAAGAAAAATGCTTCCGATGTGGCTGTAATAGGGTCTTAGGTGTTTGTGACTGATAAAACCACACAAAAAAAAGGTTCAAAACACTTTCATAGAAACCAGATTAACATGTCACTAAATGTAGAAGTGTGAATGTTTTTAAATGTTCAGGCAGGCGTTTTAAGATATACACGCCCAGATGCTGTCAATAAATATGCAAGTTGGCTTTTTTTGTATGATAATTCTTCTTTAGCTAAAAACCTAAGAATAACCCGTAATATGGATTAGAGTATTCGGTATAGGTGACACTTTTAAGTACACTACTTGTTACTTTAATTTCCCAAACAGTCATTTTGGGCAGTATAGGGTTAGTTGGTATCGCCGATACTCGCCTTCATTATACATGGTATCGGAGTGGATAGGGAAATCAGCGGTATCGCACATCACTACTCTGTGGCAGAACTCATCTGTGCACTCGCAGATGCAGGCGAAGCAGCGCGAGGCGGGGCAGGTGTAAAGCTGTTTGATTGGCTGATGCCTGACCAATGAACAATGCCCAGCAGCCAACAGGACTGAGGTGCAAAAGAGTAATTTCCACCGTTATTTTATTGTTTAGGGTTTATTTCAACTATTTTCTGAAAATATTTTTGTTGAATAGCATTAATTCTACGGAAATGTACATACCAGTAAAGAGCAGCAAAGCCGGTTTGTTGGCTGAGAGCCGAAAAGCCAATGAGATGGGACGTTTCTGTACGTCAGCACACAGGGCAGCTCACGTTGTTGGAAGAATCAAGCAATCAGGCGGGTAAGTTATTATTTTTAAAAATATCTATATGTTCAATACTGTTAGCTATGCTAAGGACTGTGCTAGGTCATACCTGTAGTTTGTTTTTATATTAGAAATGGATTTAGGAGCTGCTTTCAATGATGTTATACGGTTCTAAAGATACAATTTAGCAGATGAAATGAAATTGTAATAGTATTTATAGTAAACAATATTACGTTTTTGAACCATACTAACTATAGTAAACTGTATTATACTGTATACTGCGAGTATCTACAACTTTAATAAATTGCAGTATACTTATATATATACAGACACACAAACACACACACACACACACAGTTTAAAGTTGAATAAATTGAAATATAGCTATCTGTTTAAAACTAAACATTCTAATGTATTCATTTGTAAACCTAATACATTTTAAAGCAGTTTCATTTCTTTCTGATTTATACATGTACATACATTACATCTGATTTAGGTCCACAACTTGCATCCTTCAACATGATACGTGTGTCCAAAGGAGCAACTAGGTGAGAGGCGGTGTCCATTCCGCTCGGCTGAGGGTCTACACATTGAGCATCTTATCTGCAGGGTTAAGCAACATTAGCTGGGCCAGTCATGTCTCTGTACAGCGAGCATCAACCAGCTTTACACAACAGCCTGCTTTCTGCCCTGTAGTGAAGGCTTGAAGGGTCACCAAGTTGTGAATTTAATCAAAGATGCTAATACATGTACATGTATAAATCAGAAAGAAATGAAACTGCTAAAATGTATTGGGTTTTTTGCAAATGAATACATTAAATATTTAGTTTAAACAGATAGCTATATTTCAATTTATTCAACTTTACAAGCAGTGTGTGTGTGTGTGTTTGTGTGTCTGTATATATATAAGTATACTGCAATTCTCAAGTGGAAGCTGCTACGGTATACAGTACTCAGCACAAATTTACTATAGTTGAGCATAGTTCAAGCATGTGTGTGACATAAATACTATTACAATTTGGAACAGAAAATGCTATTTCTTCTAACATAAAAAACAAATACAGAAGCACTAGCACAGTCATAACGCCTTTAACAGTATTGAGTATATAGATATTTTAAAATAATAACTTACCACTTCTGATGTGATTTCTTCCAACAGCGTGAGCCTGTGCGGCATTACAGAAACGGCTCGGCTGGCTCTAATAGAGCCGGCTGTGCACAGCATGGCACATTTCCGTAAATTAATACATTCCAACAAAAACTATTTTTCAGAAAATGGACTCGACCTAGGCAATAAAAATAACGGTGAGGAAATACTCTTTGCACCTCAGTCTGTTCAGCTGCTGGCGGCCATTGGTCAGGCATCGGCCAATCAGCCAGCCACACCTGCCCCTGGCTGCGCACCATCTGGAGTGCACAGGCTAATGCTACAGGCTCTTGAGATGTGTATGCCCAAAAGTCAGGGCGAGAGATTTGTAGTTGTACTGAAAAATCTGAGAGGTTGTGAGTGTCGGCCTGTGCTTGTAGATCGAAGATGCAATCAAATTTGCGGTGCACAGAGTGTGTCTGTCATTTTGTATTTATGAATTGACATTTACAAATACACAACGCTATTAATCTGCAACTTACAATTAAAACACAAGTGTGTGTGTGTGTGTGTGTGTGTGTGTGTGTGTGTGTGTGTGTGTGTGTGTGTGTGTGTGTGTGTGTGTACCGTCTGCTGCGACGGGAGGAGTGACAGATACAGCTGAGAGTAAAGATCAATATGCAGGTTTATTCAGCGTCAGGCAGCCAATGGTCAAATCAGGTACAATGAGTTTATGTGCAGGCAAATCCAAAATCATAGTCGTATTAACAGGCAACAGATCAAAAGGCAGGTGGCTAAACAGGAGAACAGGGAATACAGGGCTAGGTCTAACACGGAGAAAACAAGACAGGGAAACGCGTTGTAAAGTCACAATAAAGCAAACAAGACTCGGCCAACAGGATGGGTCAATGAATGGCTTATGTATGGTGTGTTATCAGTCTTTGACAAGGTTCAGCTGGTGAGTGTAAACAAGCTAGATGAATGAGCAGGCGTGAGTGTCTGGGAGAAATTGCAGTATGCAAATGTAGTCAGAGGAAAGGCGGAAGTGTAGTTCATGGTTATAGTGAAACCAGCGATCTCCAGGAGAGCGATCACTGGTGCTCGTAACAAAATGTCAGGATTCAGCTGTTTACATCAGAGCTGTGATGTATTTCAACCTTCACCTACAAAATATTATTATCACAAGTGCTCACATTTACATTTGCAAGCTCTCAGTTTGCTACTGATTTACCTTCATAAAATGCTCACTGCCGTTGTTGAGATTCAAATGCAAATATCTCAATCAAATCCAACGCAATGGAGTGAACTTTGTTAAATAGGTATCCGAGCCATCTTTTTCTAAATTTTATTAGCTTAATTGCTTTCTCCTTGTTAATTGTGGGTCACCAGCGGCACCGCTACAGAAGAGGTGTTAAAATTTCAGTCTGTGGCAAAAGCATGACCGGTCATCTCTCCTCACATGCGGCCTGCGCGACTGACCGATCACAGCCAGGCAGAGTTGTGCCTTTTGGGCACAGATTCCGAGCAGCACCGCTTGGAAGTTCT

GACTCTCATCACCAGTTATTGACCTGCCATCGAGATTCGTGGAAACGCCATTTAATGGACTCAGGTTTTGAGAGCTTGAATGTGTTTCAAACTATTGTCTTTAGTGGCAATCAAAGTGATTTCCCCTGTGGGCTCCAGGTTGGAGTGCTTGGGAGCTTGGCTTCTCACTGTCTGTCTGCAAGCGGGGACATGTCCTAGCCCGCCTGCCTGCCTGCTCTCCTCCCGTCCGTTTCTGTGTTTAATGGTTGGTTGTGGTATGGATTTGGGTCTTTCCTTTAAAACGCAGTCAGATGTGACTTGGCCTGCCCAGGCCCCTCCCTGAAGTTCCCACCCTCCTCCGGGGGGCTCCAGAGTTGGTGATTGGCGGGTGTATGCCACTGGCTCTCGTGCGAGGACATGTCCTGGCCTGCTTGCCCCCTGCCTGCTGTCCCTGGTCTGTTTCAGTGTAATGGTTGGTGGTGGTGGGGTTTTACCCTTCTTTTGACTACGGGCGCTGTGTGACCTGGCCTGCCAGGCCTTTCTGAAGCCCTCTCTTCGTGTTGTTCTTGAGTTGGTGCTTAGGGTGCTTGGCTTCCCACTGGCTGTCTGCATGCAGAAACAAATCCTGCCTACCTGCTTGCCTGCTATCCTTGGTCCGTTTTGGTGTTTAGTGGTGGTGGTTTTTATTGGTCTTTCGCTTTGCCCGTGGTCAGCTGTGTGACCTGGCCTGGGACCCCTCTGAAGCCCCTCCCTCGTGGTGCTCCAGAGTTGGTGCTTAGGTTGTTTGGTTTTCCACCGGTTGTCTGTGAGCAGGGATGTGCCTCGGTTTACCTACCTGCCTGCTCTTCCTATTATTAATTTTCAGTGTTTAAAGTTGACGGTGGTAAAGGTGATGAGGGAGTGTTTGTAGTTTCCTTTGCCTGTTGTCTGCCCACTATTGCTAGTGGTAATAAATGGAGTAAGAACTGTGGAGTTTGCGGGAAATTTCATTACTGTCAGG

**>45S-M-R1_5'-UR-CGA-R**

TGCAGTAACCAGCAGCACGCAGTACTTATATGGGTAACCAGCAGCAGAAGGATTCAGTCCCAACCCCGCACCCCAGCAGAGCCTTTATTGGGGCACCAGCAGCCGCGTACCCACATGGGTCGCCAGCGGCAGAAGGGGTCAGCCTGGGCCCGCACCCCACAGCAAAGCCTTTAGTGGGGCACCAGCAGCACACAGTACCTACATGGGTTACCATGGCAGAAGAGGTCCGGACCGACCCACACCCCACAACAAAGCCTTCGATGGGGTACCAGCAGCCACGCAGTACATAAATGGGTCACCAGCGGCGCGTGAGTCTGGCGACGCGCGTCTTACGAAGTCAGTGTCCCCGGG

**>45S-M-R1_5'-UR-IS-1**

CTTAGTGGGAGGGTGCCCGGGCACGGCACAGTGAACGGGTTCCATGTGGTGGGGTCAGAGACGTTCGGTGTCCCCGGCCTAG

**>45S-M-R1_5'-UR-IgSL-R_(1,118bp)**

CCAGCCAGCCAGTCAGTCAGCCCAGGCCCAGCCCAGCCCAGCCCAGCCAGCCAGCCAGCCAGCCAGCCAGCCCAGCCAGCCGTCAGCCAGCCCAGCCAGCCAGCCAGCCAGCCAGCCAGCCCAGCCCAGCGCTGTCCAGCCCAGCCACACCCAGCCCAGCCCAGGCAAGCCCAGGCGGCCCGGCCAGCCAGCCAGCCAGCAAGCCCAGGCCAGGCCAGGCGCCAGCCCAGCCAGCCACCCTGCCCTGCCCAGCCCAGCCCAGGCCGGCCAGCCAGCCAGCCAGCCAGCCCAGGCCAGGCCAGGCCAGGCCAGGCCAGGCCAGGCCCAGCCCGGCCAGGCCAGGCCCAGCCCAGCCAGCCCAGTCAGCCAACTGTCCCAGCCCAGCCAGCCAGCCAGCCAGCCCAGCCCAGCCCAGGCAAGCCCAGGCCGGCCAGCCAGCCAGCCAGCCAAGGCCAGGCCAGGCCCAGCCCAGCCAGCCAAGGCCAGGCAGGCCAGGCCCAGCCAGCCAGCCAGCCAGCCAGCCCAGGCAAGGCAGGCCAGGCCAGGCCAGGCCCAGCCCAGCCAGGCCAGCCCAGCCGAGCCCAGCCAGCCAGCCAGCCAGCCCAGCCCAGCCCAGCCCAGCCCTGCCCTGCCCTGCCCTGCCCCTGCCCTGCCCTGCCTGCCCAGCCCAGGCAGCCCAGCCAGCCCAGCCCAGGCCGGCCAGCCAGCCAGCCAGGCCCAGGCCAGGCCAGGCCAGGCCAGGCCCGGCCCAGCCCAGCCAGCCAGCCAGCCAGCCAGCCAGCCCAGGCCAGGCAGGCCAGGCCAGGTGAGCCATGCCACCAACCCAGCCAGCCAGCCAGCCAGCCCAGGTAAGCAAGCCCTGGCTGGGCAGCAAGCCCAGCTCAGCCAGCCAGCCCAGGCCAGGCAAGCCCTGGCCGGCCAGCCAGCCAAGCCCAGCCAGCCAATCAGTCAGTCAAGCCAAGGCCAGTCAGTCAGGTCAATCAGCCCAGCCCAGCCCAGCCTAGCCTAGCCTAGCCTAGCCCAGCCAGCCCAGCCCAGCCCAGCCCAGCCCAGCCCAGCCCAGCCCAGCCTGCCCTGCCAACCTGACAGCCAGGCCTGCTCAGCCAGCCAGCCAGCCA

**>45S-M-R1_5'-UR-IS-2**

TCCTGGTGTTGGAGAGGCTTGGGCTTTAGAAGTGGACTGACGATAGTCCATGGCCGGAGAAGTGAGCACTATCTCCTCCCTCCCTCCCTCTTCCTTT

**>45S-M-R1_5'-UR-ACC-R_(24bp)**

**ACCACCACCACCACCACCACCACC**

**>45S-M-R1_5'-UR-IgSL-R_(1,492bp)**

CCGGAGGCGGGGAATTGGGCTTGAGGATAGGGTAGGGTTGCCTGGAGGCTTGGGGGTGGAGTTAGGGCGCCCGGAGAACCTGGGGATTGAAGCTGGGTATGAGGTAGAAGTTGTAAGGGGCGGGTTAGGGTGCCCGGAGGCCTGGGGATTGGGCTTGGGGATAGATGCAGGGTTGCCTGGAGGCTTGGGGGAGTGGGTTAGGAGCTGCCCGAGCCTGGGGATTGGAGGCTTGGGGATAGGAAATGAAGTTGCTTGGAGGCTTGGGGGGCGGGTTAGGGGTGTCCCCGGAGGCCCGGGGATTGGGCTTGAGCATGGGTAAGGGTTGCCTGGAGGCAGGGGTGAGTTAGAGGCCTGGAGGAGCCCGAAGTTGGGCTTGGGGATAGGGTAAGGAGTTGCCTGGAGGCTTGGGGGTGGGTTAGGGCTGCGCGGGGACCCGGGGATTGGGATTGGGCTTGGGCATAGGGTAAGGGTTGCCTGGAGGCTTGGGGGTGGGTTAGGGCTGCCTGGAGGGCTTGGGAGTGGGTTAGGGCTGCCAGGAGGCGGGGATTGGGCTTGGGCATAGGGTGGGGTTGCCTGGGAGGCTTGGGGGGGTGGGTTAGGGTGTGCCTGGAGGCTTGGGGGTGGAGTTAGGGCTGCCCAGGAGGCCCGGGGATTGGAGCTTGGGCATAGGGTGGGGTTGCCTGGAGGCTTGGGGGTGGGACAGGGCTGCCTGGAGGCCCGGGGGATTGGGCAGGGATAGGGTAAGGGGTTGCCTGGAGGCCCAGGGGTGGGTTAGGGCTGCCCAGGAGGCCGGGGATTGGGCTTGGGCATAGGGAGTAGGGTTGCCTGGAGGCTTGGGGGTGGGTTAGGGGCTGCCGGAGGCCCGGGGATTGGGCTTAGGGATAGGGTAGGGTTGCCCTGGAGGCTTGGGGGTGGGTTAGGGCTGCCAGGAGGCCCGGGGATTGGAGGCTTGGGCATAGGGTAAGGGTTGCCTGGAGGCTTGGGGGTGGGTTAGGGCTGGGAGGCCGGGGATTGGGCTTGGGGATAGGGTAGGTTGCCTGGAGGCTTGGGGGTGGAGTTAGGGCTGCCCAGGAGGCCGGAGGGGTGGAGACGGAAACGCCAAGGTAAGGTTGCCTGGAAGGCTTGGGGTTAGGGCTGCCTGTGGAGGCCCGGGGATTGGGGCTTGAGGATAGGGTAAGGTTGCTAGGCTTGGGGGTGGGTTAGGGCTGCCAGAGGCCCGGGGATTGGAAAGCTTGAGCATAGGGTAAGGGTTGCCTGGAGGCTTGGGGGTTAGGTTAGGGTTGCCCGGAGGCCTAGGGATTGAGCTTGGGAGATAGGAGTAGGGTTGCCTGGAGAAGCTTGGGTGTTGGGTTGGGGTTTCCCCGGAGGACTAGGAACTGGGTTAGGGTTGCCCTGGAGACTTAGGGGTGAGGTTAGGGCTGCCGGAGGCCTGGGGATTGGGCTTGGGGATAGGGTTAGGGTTGCCCGGAGAGGCTTAGTTGTTGGGCTTGTG

**>45S-M-R1_5'-UR-IS-3**

GAGCTTTGGCTTGCCTGGGTTTGCCTGAGGCTTGGGAGAAGTGCCACCCAGCGTGCCAGCCAGAAAGGTCAGCTGTGGGTGACCAGCAGCACCGCTCCGGCTGCCGAAGAGCAAAGTCCAGTCGCGGGTCACCATCGGCACCCGGTCGCGGAAGCAAGAGCGGCCGCTAACCTGACTATCCATGGCCCCCGAGCTCAGGTTCACAGCCGGGCGGGACGTGACCTCTTGGGTCTCCTCTCTCCCGGAGGGTCTGGGCAGCCGTCACTGACAGTTATCGCCAGCCTGCCGTCGGAGGATTTC

**>45S-M-R1_5’-ETS**

GGGAAGGCGTGGTTAGTGGACTTAGGTTTTGACGCGTAACTATGTGCCACCGCCTTCCACCACCTGACCGATGGGAATTGTGCGCGTACCCCAGGCAGGCCACCTTGATCCTCCGACAAAAGGGGACCGCTGTGGGTGGGGTGAGTTCCTCCATTTGGCATTCGAAAGGAGGGCGAGCGGGCCTTCACAGACCTCGGCCCTTGCTCCTTTCGATTTAAGGCTTTCAAGATGGCCAGTCGATCTCAGACCGCCCTGTACAGCTGGAACAGTGCGCTAGTCGGTGCTGTGCGTGCGCTCTGCAGAACCATCCTCCTTCCCCCCTCCTCTTCCTCACCACCGGGGTAATCCGGGCGCGTGAGCTGACTTGGGGGTGAAGCTGTGTGGCCAGGCGCGCGGCCATCTGCAGCCTTTCGATCCGGTGCAACACGATCCTTCCTGGTGATGGCCTCGACCAGCGGCTGGCCACGCCCGGCAGACCCCCTCATTTCCATGTGTGCGGTGGGGGGGTGGGTAGGCGGCGGTACCTGAGCCCAATCCCCCCTCCCGCCCAGCCCGCGTGCCCCCTCCTGCGAGCAGTCGTGACGCCTCGCGGGAGCGAGACCCGCTGCCAGTGGCGGCGCGGGCTGTGGTGCGGGCGGGGAGGGTTCTCTTCCGCCGAGTCCTTTCCCTCTCTCCTCCGGCACTCTCGGAGAAAGGTGCCACCACCACGCGATTCACCCGGCGAGGC

**>45S-M-R1_18S**

TACCTGGTTGATCACTGCCCGGTAATATGCTTGTCTCAAGGTAAGCCATGCAAGTCTAAGTGCACACGGCGGTACAGTGAAACTGCGAAATGGCTCATTAAATGGTCTGGTTCCTTTGATCGCTCCACCCGGTTACTTGGATAACTGTGAATTCCAGAACTATGCAGCTGCCAGCGCCGACCTAGGCCGCCCACTTCTCCCCTCGGGGCGGGAGGTGAGTACCCAGGGGACGAAGCGTGCATTTATCAGATCCAAAACCCATGCGGGTGCGTGGGCGGTGAAGAGGGGGCCCTCGCGCCTACCCGCCGCCGTCGCCCCCGGCCTCGCTTTGGTGACTCTGGGCCTGCGGGCGATCGCGCGCCTCGCGGCGGCGACGGTTCGGCGTCGAATGTCTGCCCTATCAACTTTGATGGCCCGGTCATTACGCCTACCATGGTGACCACGGAGTGACGGGAATCAGGGTTCGATTCCGGAGGAGGAGCCTGGCTTGGCTGCCACCACGTCAAGGAAGGCAGCAGGCGCGCAAAATTGCCCATTTCCGACACGGAGAGGTAGTGACGAAAAATAACAATGCAGGTCTCTTTCGAGGCCCTGCAGGTGGAGTAAGGTGCATCCCAAACCCATGGCGAGGACCCATTGGAGGGCAAGTCTGGTGCCAGCAGCGCGGTAATTCCAGCTCCAATAGCGTATGCTAACGTTGCTGCAGTTAAAGCCTAGTAGTTGGATCTCGGGGACCGGGCGCGCAGTCGCCTTGTGAAGCGAGCCACACGCCCGGTCCCGGACACAGGCCTCCAGCCGGTGCCTTGACTGGGTGTACCGGCTTTAGGGCCGGGCGTTTACTTTGAAAGGGTAAGTGTTCAAGGCAGGGCGGCACCGCGCTTCATTGAATACCCCAGCTAGGAATAATGGAATAGGACCCGGTTCTATTTTCTGTAGGTTTCGGAGCGGGGGCCATGATCGAAGGGGCGGCCGGGGGCATTCGTATTGCGCCGCTAGAGTGAAATTCTTGGACCCGGCGCAAGACGGACCCGGAGCGAAAAGCGTTTGCCAAGAAACGTTTTCATTAATCCAAGAAGCGAAAGTCGGAGGTTCGAGAAGACGATCAGATGCACGTCGTAGTTCCGGCCGTAAAACGTGCCGACCCGCGATCCGGCGCGTTTATTCCCATGACCCGCCGGGCAGCGTTGCGGGAAACCACGAGTCTCTGGGCTCGGGGAGTATGGTTGTGGCAGAAACTTAAAGGAATTGTGAGGAAGGGGCACCACCAGGAGTGGAGCCTGCGGCAATTTGACTCAACACGGGAACTCACCGGCCCCCGGACACGGAAAGGATTGACAGATTGACGGCTCTTTCACCGATTCTGTGGGTGGTGGTGCATGGCCGTTCGTAGTTGGTGGAGCGATTTGTCTGGTTGATTCCGATAACGAACGAGACTCGGCATGCTAACTGAATTCTGGCCTTGATCGGCGTCTGCAACTTCTTAGAGGGACAAGTGGCGTTCAGCCACGCGAGACTGAGCAATAGCTGGAGTCTGTGATGCCCTTAGATGTCGGGGGGCTGCACGCGCGCCACAATGGGCGGATCACGTGTGCCCTGCGCCCGACAGGCGCGGGTACTCCCGTTGAACCCCGCCCGTATTAAGGGCCGAAGTTAAACTATTTCCACACGAGAGCAGGAATTCCCAGTAAGCGCAGGTCATCAGCTTGCGTTGATTAAGTCCCTGCCCTTTGTACACACCGCCCGTCGCTACTACCGATTGAGCGGCTCAGTAGAGTCCTGGGATCGGCCCCGCCAGCGGCTCCCTTACCGGGAGCCCTGGTGGAGCGCCGAGAAGGCAGTCAACTCGGTCGTTTAGAAGGAAGTAAAAGTCGTAACAAGGTTTCCGTAGGTAGACCTGCGGAAGGATCGTGG

**>45S-M-R1_ITS-1**

GAATTGAGGGATCTCCTCCTCACGCCCAGAGGCGAAGCCACGGTAAGCTTCCGCGCGGTGGGAAGTCCCTACGGGTCTACCTCCCCACCATCGCGCGCGCGGCGTGATCCCGAGTGATCAAAGGTTGGCGTCGCGGGCTCCCGGCGGGTACCCGGTTGGTCTCGACCACCTCGGCCCTGTCCCCTCTGCGGGGGAGGCGCGAGTGCGTGGGGGCCGTGGGTTTAAAAAGAAAAGCACTCTTCGCGTTTCCCCACCCGGGAGGAAGCAGAGAGGAGACGCCCGTCCCGGGGCCTGCCGGCCCGTGTTTAATTTCCCCACCACCCCCCTCCCGAAAGCGTCCTCTGTCTCGGACCGTAACGATTGAAAACGAAAACAAGAGTGTAC

**>45S-M-R1_5.8S**

AACTCTTAGCGGTGGATCACTCGGCTCGTCGTCGATGAAACGCAGCTAGCTGCGGAACTAATGTGAATTGCAGGACACACATTGATCATCGGCCTTTCGCGAACGCACATTGCGACGGGTCCATCCCGGGCCACGCCTGTCTGAGG

**>45S-M-R1_ITS-2**

TAGCCTTGCATCGATCGGACGGGAAAGAGTGGAGTCTTCGTGCCGCCCTGCCCCCTGTCCGCGGCTGGGGCGTCGCAGGCCCTGCCCTCCCGCGGTGGCCTACGTCCTCCCAAGTGCAGACCGCCCGGAGCCGTTGATCCGCCCGCTTGGGGGCGGCTCCCATTCTCTCCCCCTCGCGCGGCTGCCGGCGGTCTGCTGCAGCTGCCCGCGCGCGTGGGCAGGAGCTGCAGCCTGAAGCTCCCTCGCGTTCGGGAACGGCGACGGCGCGTGGTGGCGCGGGCCGGCGTCCCGCCTCGTGTGGGGCGACCGCGCCACCACCCGCCTGTTGGTCCC

**>45S-M-R1_28S**

ACGACCTCAGCTCAGACGAGAAGACCCGCTGAATTTAAGCATATTACTAAGCGGAGGAAAAAGAAACCAGCAGGATTCCCCCAGTAGCGGCGGCGAAGAGGAAAAGTCCAGCGCCGAATCCGCCCCTCTGCCGAGGCGAGGACCTGTGGCGTACGGAGGGCCGCCTCTCTCTCGAGCGCGGGCCGAGGGCCAAGTCCTTCTGATGGAGGCTTAGCCCGCGGACGGTGTGAGCCGGTGTCGGCCCCTGCCGCCGAGTACGGTTCCTCCCGGAGTCGGGTTGTTTTGGGAATGCAGCCCAAAACGTTGTGGCTCATCTAAGGCTAAATACCCGGCTAGGGCCGATAGCGGACAAGTACCGTGAGAAGAAGTTGAAAAAAAGAACTTTGAAGAGAGAGTTCAACAGGGGCGTGAAACCGTTAAGAGGTAAACGGGTGGGGACCGCGTCCGCCCGGTGGATTCAGCCCGGCGGGGGGGTCGGCCCGTCCGGTGCGCGCTCCCTTCGCTCCCCTCATTCCTGGGGGTGGCGCGGGGATTGGCCGCCCGAAGCGAGCTCGCCGCCGCAGGTGCATTTCGCGCGGTGGGCGCCGCGACCCGGCTCAGTTCGGCTTGGAAAGGTCAGGGGAGAGAAAGAGTGGCCCGTCGGTTCAGGCCGTCGGGCTTTACAGCGCCCTCCCGCCGACCGCGCTTGCTCTCCCGGGGCCGCGGGTGAGTGTCCTCCGCGCCTCTCTGCCCTCCCTCCCTGTGCGTGGGGGGAGGTGCGAGGCAGAGTCTCCCCCGCCCCCGGCGTGGCGACACCCAGGGTGGACTGTCCTGATCCGCCTGGCTGCGCCCAGCGCCGCCCAGAGAGGCGAGATCCGACCCCACGTTCGGGCGCCCGGAGTCCCGCGGCGACGCCGGCCTCCCACCCGGCCACGTCTTGAAACGGACCAAAAGAGTCCAACGCGCGCGCGAGTCAGAGGTGGCTCGCGAGCCCTGCGGCGCAATGAAGGTGAAGCAGGGGTCTCCCGGGTGGGATCCCCCGCCGCGGGCTGCAGGCCGCCTCGCGCGACCCTCCGAGGGCAGGTAGGTGGAGCGCGCGCGATGGCACCCGAAAGATGGTGAACTATGCCTGGGCAGAGAGCGAAGCCAGAAACTCTGGTGGAGGCCGCGCGGTCTGACGTGCAAATCGGTCCCGTCCGACCTGGGCATAGGGCGAAAGACTAATCGAACCATCTAGTAGCTGGTTCCCTCCGAGAAGTTTCCCTCAGGATAGCTGGCGCTCGCCCAGTCAAAAAGCAGTTTTATCCGGTAAAGCCAATGACTAGAGGCCTTGGGGCGAAACGGCCTCAACCTATTCTCAAGCTTTAAATGGGTAAGAGGCCCGGCTCGCTGGCGCTGGGCCGGGCGTGGAATGCGGCGCGCCCTGGTGGGCCATTTTTGGTAAGCAGAACTGGTGCTGCGGGATGAACGAACGCCGGGTGGCGCCCGATACCCGGCGCTCATCAGACCCCATAAAAGGTGTTGGTTGATATAGACAGCAGGGCGGTGGCCATGAAGGCTGGCACCCGCCAAGGGAGTGTGTAACACCAGCTCTGCCGAATCAACTAGCCCTGAAAATGGATGGCGCTGGGCGTCGGGCCCATACCCGGCAGTCGGCGGCACAAGGGACACGCGGCCGGTCGCGCGCTGCAAGCCACTCGGCGAGTAGGAGGGCCGCGGTGGCGCGAGAAGCAGGCGGGCCCGGGTGGAGCCGCCGCGGGCGCAGATCTTGGTGGTAGTAGCAAATATTCAAGCGAGGCTTTGAAGGCCGAAGTGGAGAAGGGTTCCATGTGAACAGCAGTTGAACATGGGTGAGCGATCTAAGGGGCGGGCCTACGCCGTTCGGAGGAGGGCGATGGCCTCTGTCGCCCCGCTCGACCGAAAGGGAGTCGGGTCCAGATCCCAGACGGAGCGGAGAGCAGGCGCCCGCGGCGCCCAGTGCGTGGCATAAACGAACCGGAGATGCCGGCGGGTGCCCGGGAAAAAGTTCTCTTTTCTTTGTGAAGGGCAGGCGCCTGGAGCAGGTTCGCCCGAGAAGGGCCCGCCTGGAAGCGCCCGGCTTCTGGCGGCGTCCGGTGAGCTCGTCGGCCTTGAAAATCCGGGGGAGAAAGGTGTAAATCTCGCCGGGCCGTACCCTTATCCGCAGCAGGTGCTCCAGGTGAACAGCCTCTGGCGTGTTGGAACAAGGCAGAGTAAGGAAGTCGGCAAGTCAGATCCCGTAGCTTCGGGGATGGATTGGCTCTGAGGCTGAGCCGGTCGGGCTGAGGTGCGGCAGGCGCTGGGCCGAGCGCGACTGGGGGAGCGGCCGCCCGGGTGCCCTGACCCCGTTCCCGAGCCGCGGGGGTGGCAGCCTTCGGTGGCGCGCGGTCCCTCTCTCCCCCCGCCGCGCGTCCGCCTCGGGCCTTTGCGGTCTCGGCTCCCCCGGCTCTTCCCCTCCGCCCTACGTCCGCGGGGGTCGGGCGGGGTCTCGGAAGAGGGGGTGGAAAGAGGCCGTGGGGAGAAAAGGGTGGATGGGTGGCGTCGGGGGCAACGGGAGGTCTCGCCTGCCGCGCGATGCCTCCGGCGGCCGGGGCCCGCAGGCGAGATGCGCGCGGTGCGCGCGGCCCTGGCGCGCGCCCGTCACTCCATGGTCTCCGCAGCTACGGCCCGCACCGGGAGCCGCGACCGCGCGTGCGCCCCCTCCCGGGAGTGGCGCGCGCGGCCTCGCTCCCGGTGCGAAGCCTCGGCTGGCGGCTGGCAGCCAGCTTAGAACTGTCGCGGACCAGGGGAATCCGACTGTTTAATTAAAACAAAGCATCCTTGAAAGGCCCTCGGCGGAGTGTTGACCTTGATGTGATTTCTGCCTAGTGCTCTGAATGTCAAAGTGAAAAATTCAACGAAGCGCGGGTAAACGGCGGGAGTAACTATGACTCTCTCCTGAGTAGCCAAATGCCTCGTCTAATTAGTGACCGCATAGATGTTAAGATGGCAAGATTCCCACTGTCCCTACCTGCTGCCTCTGCAGAAACCACAGCCAAAGGCGAGGCTTGGCAGAATCAGCGGGAAAAGAAGACCCTGTTGAGCTTGACTCTAGTCTGGCTGTGAAGAGACATGAGGGGTGCTGTAGAATAAGTGGGAGGCCCCAGGGCTTCCCGGAGCCGGCCGCGGTGAAATACCACTACTCTTGCCGTTTCCTCACTTACCCGGTGAGCGGGGAAGCCGGGCGTCCCCCAGCGCAGGGGCGCCCTGCTGCGTCAAGCGCCCCGGGGCCGTGCGGGGGTGGGAACCGCAGCCCCCTTCCTCCCGGCCGCCCCGACCGGGCGCGGCCCGCCCGGGGGACAGCGTCAGGTAGGGAGTTTGACTGGGCGGGCCATTTCCTGTCAGTGGTCACGCAGGTGTCTAGGCGAGCTCAGGGGGACAGAAACCTCCCCGTAAAGAGCAGAAGGGCCAAAAGCTCGCTTGATCTTGATTTTCAGTATGGTGCACGGACGCGCGGCAGGCCTCACGATCCTTCGGCTTTTGGGGTTTTAAGCGGGAGGTGTCCAGAAAAGAGATTACCACAGGGATAACTGGCTTGTGGCGGCCAGCGTTCATAGCGACGTCGCTTTTTGATCCTTGATGTCGGCTCTTCTATCATTGTGAAGCAGGAATTCTGTTAGCATTGATTGTTCACCACTAACAGGGAGCGTGAGCGTAGGTTTGGGCAAAGTCGTGAGACAGGTTAGTTTTACCCTACTGATGTGAGCCGTTGTTGCAATAGTAATCCCGCTCAGTACGAGAGGAACCGCGGGTTCAGACATTTGGTCCGTGCGCTTGGCTGAGAGCCACTGGCGCGAAGCCTGCATCTGCGGGATTATGACTGAGCGCCTCTAACAAGTCAGAATCCCGCCTAAAAGCTGCTTGATACAGCAGCGCCGTCGGATCTCCGATAGGCCCCGGAGTAGCAGGAAAGCCCCTCTCGGGGGGCCCCGGCGCGGAAGAGCCGTATTCGTGACCAGACAAAGCGCGGCTGAAGGCTTCGCCCCTCTTGTGGCGCTACCGCATGTTTGTGGGAACCGGTGCTTAAATGACTCGTAAACGACCTGATTCGGTAGGTATGCGTGCGTGGAAGAAGCAACTCTGTTTGCTGCGATCCATTGAAAGTCAGCCCTCGATCCAAGTTTTGTC

**>45S-M-R1_3’-ETS**

GGGCCGGACGCAGGCGACAGGGCCACGGTCCGACAGACCCGCGGCGTGCGAAGGGGCA

**>45S-M-R1-R2_IRUS**

CCAGCTGCGTCGGTGAAGGCTTGCCATCCCCCTCGCCATAACGCCCAGGCCGAGGCACCAGCGGCGGGGAGCGTGTCGCGATCCCAGCCTGGGGCACCAGCGGCAGGGGAGGCCCTGGCCTCGG

**>45S-M-R2_5'-UR-CGA-R**

CCCACCCCCACCCAGCAAAGCCTCCCTGCCTGAGGCACCAGCAGCACGCAGTACCTATGTGAGACTTGGCCAGAAGGGTTCTGTCCGACCCTAACCCCCACAGCAGAGCCTTTATTGGGGCACCAGCAGCAGTACCCACGTGGGTCGCCAGCGGCAGAAGGGGTCCAGCCTCGACCCGCACCCCACAACAAAGCCTTTAGTGGGGCACCAGCAGCACGCAGTACCTACATGGGGTTACCAGCGGCGAAGAGGTCCGGACCCGACCCACACCCCACAACAAAGCCTTCGATGAGGTACCAGCATGCCACTGACCCATAAATGGGTCACCAGCGGCGCGTGAGTCTAGCGCGCGTCTTACGAGAATCCAGTGTCCCCCGGG

**>45S-M-R2_5'-UR-IS-1**

CTTAAGAGTGGGGTGCCCGGGCACGGCCTGTGAGCAGTTCCATGGTGGGGTCAGAGGCCGGCCGGTGTCCCCCGGCCTAG

**>45S-M-R2_5'-UR-IgSL-R_(1,212bp)**

CCAGCCAGCCAGTCAGTCAGACCCAGCCCAGCCAGCCCAGGCCTTGGCCAGGCCAGGCAAGCCCTGGCGGCCAGCCAGCCGCCCAGCCAGCCAGCCAGCCAGCCCAGCCCAGCGCTGTCCAGCCAGCCTGCCCAGCCCAGCCCAGGCAAGCCCAGGCCGGCCGGCCAGCCAGCCAGCCAGCAAGCCCAGGCCAGGCCTTCACTCCAACCAGCCCTGCCCTGCACCACTGCCCAGCCCAGCCCAGGCCGGCCAGCCAGCCAGCCAGCCCAGGCCCAGGCCAGGCCAGGCTGAAGCCAGGCCAGTGAGCGAGCCAGGCCAGGCCCAGCCCGGCCAAGTTGAGCCCCAGCCAACCAGCAGTCAATGGCCAGCCAGCCCAGTCTTCCAGCCAGCCGCCCAACCCAGTCAGCCAGCCAGCCCAGCCCAGCCGCCCAGCCCAGCCCAGGCAAGCTGACCAGCCAAGCCTGCAGCCAGCCAGCCAAGAGCCAAGAGCCAGTGAGCCCAGCCCAGCCAGCCAGCCAGCCAGCCAGCCAGCCAGCCAGCAGCCAGCCAGCGGCAAATAGCCAGCCAGCCAGCCCAGGCCAGGCCCAGCCCAGCCAGCCCAGTCAGCCAGCCAGCCAGCCAGCCAGGCCAGCCCAGCCGAGCCCAGCCAGCCAGCCAGCCCAGCCCAGCCCAGCGTTGCCCAGCCAGCCCAGCCCTGCCCTGCCCTGCCCTGCCCAGCCAGCCCAGCCCAGCCAGCCAGCCAGCCACTTGTTGTAAGCCAAGCCCAGGCCGGCCAGCCAGCAGCAGCCAGCTGAGCCCAGGCTTGAGGCAGGCCAGGCCAGGCCAGGCCAGGCCCAGCCCAGCCAGCCAGCCAGCCAGCCAGCCAGCCCAGGCCAGGCCAGGCCCGGAGGCCAGGCCAGGCCAAGGCCAGGCCATGCCCAGCCAGCCAGCTTCCAGCCCAGGCCAAGCAAGCCTGGCTGGGCAGCAAACTTAGCTCAGCCAGACCAGCCCAGGCCAGGCAAGCCTGGCCGGCCAGCCAGCCAAGCCCAGCCAGCCAATCAGTCCAGTCAAGCCAAGGCCAGTCGGTCAGGTCAATCAGCCCAGCCCAGCCCAGCCTAGCCTTCCCAACCCTAGCCTAGCCTAGCCCAGCCAGCCAGCCCAGCCCAGCCCAGCCCAGCCCAGCCTGCCCTGCCCAACCTGACAGCCAGGCCTGCTCAGCCAGCCAGCCAGCCA

**>45S-M-R2_5'-UR-IS-2**

TCCTGGTGTTGGAGAGGCTTGGGCTTTGAAATGGATGTGATAGTCCATATGGAGGGAGAGTGAGCACTATCTCCTCCCCCTCCTCCTCTTCCTTT

**>45S-M-R2_5'-UR-ACC-R_(21bp)**

**ACCACCACCACCACCACCACC**

**>45S-M-R2_5'-UR-IgSL-R_(1,554bp)**

CCGGAGGCCGGGGATTGGGCTTGGGGATAGGGTAAGGGTTGCCTGGAGGCTTGGGGTGGAGTTAGGGCTGCCGGAGAGCCTGAGGGATTGGGCTTGGGTATATAAGCAAGTTGCTTGAGGGGCGGGTTAGGGTTGTCGGAGGCCTGGGGATTGGGCTTGGGGATGGGGGTACGGGTTGCCCTGGAGGCTTGGGGAGTGGGTTAGGGCTGCCCGGAGCAGACCTGAGTTGGGCTTAGGGATAGGGTAAGAGTTGCTTGGAGGCTTGGGGGCGGGTTAGGAGTGTCCCGGAGGCCCGGGGATTGGAGGCTTGAGCATAGGGTAGGGTTGCCCCTGGAGGCTTGGGGTGAGTTAGGAGCTGCCTGGAGGGCCCGAGTTGGGCTTGGGGATAAGGGTAAGGGTTGCCTGGAAGGCTTGGGAGCAGGTTAGAGGGCTGCCAGGAGGCCCGGGGGATTGGAGGCTTGGGCATAGGGTAGGGTTGCCTGGAGGGAAGCAGGGTGGTTAGGGCTGCCTGGAGGGCTTGGGAGTGGAGTTAGGGCTGGAGGCCCGGGGGGATTGGGCTTGGGCATAGGAGTACGGGTTGCCTGGAGGCTTGAGGGTGAGTTGGGCTGCCTGGAGGCTTGGGGTGGAGTTAGGGCTGCCAGGAGGCCGGGGATTGGGCTTGGGCATAGGGTAAGGGTTGCCTGGAGGCTTGGGGTGGAGTTAGGAGCTGCCTGGAGGCCCGGGGGATTGGGCAGGGATAGGGTAGGGGTTGCCTGGAGGCTTGGGGGTGGAGTTAGGGCTGCCAGGAGGCCCGGGGAGTTGGGGCTTGGGCATAGGGTAAGGGTTGCCTGGAGGCTTGGGGTTAGGGCTGCCCTGGAGGCTGGGAGTGGGTTAGGGCTGCCAGGAGGCGGGGATTGGGCTTGGGCATAGGGTGAAGTTTGCCTGGAGGCTTGAGGGGTGGGTTAGAGTTTGCCTGGAGGCTTGGGGAGTGAGTTAGGGCTGCCAGGAGGCGGGAGTTAGGAGTTAGGGTTAGGGTAAGGGTTGCCTGGAGGCTTGGGGGTGGAGTTAGGGCTGCCTGGAGGCCCGGGGGGGTGGAGGCTTGGGGATAGGGTGGGGTTGCCTGGAGGCTTGGGGGTGGGTTAGGGCTGCCAGGAGGCCCGGGGATTGGGCTTGGGCATAGGGTCTGCGGTTGCCTGGAGGCTTGGGGTGGGTTAGGGCTGCCTGGAGGCCCGGGGATTGGGCTTGGGGATGGGGTAAGGGTTGCCTGGAGGCTTGGGGGTGGAGTTAGGGCTGCCAGGAGGCCCGGGGGATTGGGCTTGAGCATAGGGTAAGGGTTGCCTGGAGGCTTAGGGGGGGTTAGGGTTAGGGTTGCCCCGGAGGCCTGGGAGGTTGAGCTTGAGGGATAGGAGTAAAGGGTTGCCTGGAGGCTTGGGGGTTAGGGTTTCCCGGAGGACTGGGAATTAGGTTAGGAGTTGCCTGGAGACTTGGGGGTGGGGTTAGGGCTGCCCGGAGGCCTAGGGATTGGGCTCGGGGATAGGGTTCAAGATTGCCCGGAGGCTTAGTTGTTGGGCTTGT

**>45S-M-R2_5'-UR-IS-3**

GGGCTTTGGCTTGCCTGGTTGCCTGGAAGCAGGAGAAGTGCCGCCGCGTGCCAGCCAGAAAGGTCAGCTGTGGGTGACCAGCAGCACCGCTCCGGCTGCCGAGAAAGGCAAAGTCCAGTCGCGGGTCACCATCGGCACCCGGTCGGCAAGGGAGAAGCAAGCGGCCGCTAACCCTGACCATCCATGGCCCCCGAGCTCCCAGGTTCACAGCCGGGCGGGACGTGACCTCTTGGGTCTCTCTCCCGGAGGTCTGGGCAGCCGTCACTGGTTATCGCTGACCTGCCGTCGAGGTTCC

**>45S-M-R2_5’-ETS**

GGGAAGGCGTGATTTAGTGAACTTAGGTTTTGAGCGCGTAACTACGTGCCACCGCCTTCCACCACCTGACCGATGGGAATTGTGCGCGATGCCCCAGGGGCAGGCCTTGCGGTCCTCGACAAAGGGACCGCTGTGGGTGGGTGGGTTCCTCCCGTTTTGGGCATTGAAAGGAGGGCGGCGGGCGCTTCCCCCAGGCCTCGACCCTTCTTGCTCCTTTGGGTTAAAGACTCAGATGGCCCGTCGATCTCAGACCGCCCTGTACCGACGGAACAAGTGCGCTGAGTCGGTGCTGTGCGTGCGCTCTTACCGGGCAGCATCCCTCCTTGCCCCCTCCTCTTCCTCACCACCCGGAGTGTAATCCGGGCGCGTAGAGCTGACTTGAGGTGGGGCTGTGTGGTAGGCGCGCGTGGCATCTGCGGCGCACTCGTCCCGTTGCGAAGCACGATCCTTCTAAGTGATGGCCTCGACCCAGCGGCCAGCCCGCGCCGGCGGAGCCCCCTCATTTCCATGTGTGCGGTGGGGGTGGTGGCGGTACCTGGGAGCCCAATCCCCCTCCCACTTAGCCCGCGTGCCCCCTCCCTGCGAGCAGTCGTGACGCCTCGGCGGAGGGCGAGACCACGCTTGTAGAGTAGCGGCGCGAGCTGTGGTGCGGGCGGGAGGGTTCTCTTCCGCAGTCCTTTCCCTCTCCTCCGGCACTCTCGGGCGCCACCACCCACACGGGTCACCCGAGCGAGG

**>45S-M-R2_18Sp_1,374bp**

CTACCTGGTTTGATCCTGCCAACTAAGCCTGCTTGTCTCAAGAATTAAGCCTATGCAAGTCTAAATTGCACCCACGGCGGTACAGGTAAACTGCGAATGGCTCATTAAATCAGTTGGTTCCTTTGATACGCTCCCTGGTTTACTTGGATAACTGTGGCAATTCCAGAGCTAGCCCATACTCCTTGGCCGACCTGGGCCGCCTTCTCCCCTCGGGGCGGGGTGGGTACCCGGGGACGCGTGCATTTATCAGATCCAAAACCCATGCGGGTGCGCGGGCGGTGGAGAGGGGGCCTCGCGCCTGCCCGCCGCCGTCCGCCCCCGGCCTCGCTTTGGTGACTCTGGTGCCTCGGGCGTCGCGTAGCCTCGCCGGCGCGGCGCCCGGTTCATTCGAATGTCACTGCCCTATCAACTTTCGATGGTAAGGTCAAGTCGCCTACCATGGTGACCACGGGTGACAGGGAATCAGGAGTTCGATTCCGGAGAGGGAGCCCTGAGAAACGGCTACCACATCCAAGGAAGGCAGCAGGCGCGCAAATTACCCATTTCGACACGGAGGGGAGTAGTAATGAAAAATAACAATGCAGGTCTCTTTCGAGGCCCTGCAGGTGGAATGAGTGCATCCCAACCCATGGCGAGGACCCATTGGAGGGCAAGTCTGGTGCCAGCAGCGCGGTAATTCCAGCTCCAATAGCGTATGCTTGCGTTGCTGCAGTTAAAAGCTCGTAGTTGGATCTCGAGGGGACCGGAAGGACCGCGCAGTCCCCGCGAGCGAGCCACCCCGCGGTCCGGACCCCAGGCCTCCAGGCGCCCCCGGATGCCCTTGACTGGAGGTGTCCTCGGCTTGAGACGGAGCGTTTACTTTGAAAAAATTAGAGTGTTCAAGGCAGGAGCCGGCACCGCGCCCCATTGAATACCCCCAGCTAGGAATAATGGGAATAGGACCCGGTTCTATTTTCTGTGGTTTCGAACCCGGGGCCATGATCGAGAGGACGGCCGGGGGCATTCGTATTGCGCCGCTAGAGGCAGAAATTCTTGGACCCGGCGCAAGCGGACGGAGGGCGAAAGATTTGCCAGAACGTTTTCATTAATCAGAGCAGTCGGAGAGTTCGAAGACGATCAGATACCGTCGTAGTTCCGACCGTAAACGATGCCGACCCGCGATCCGGCGGCGTTTATTCCATAACCCGCCCGGAGCAGCGTTGCGGGAAACCACGGGGAGTCTCTGGGCTCCGGGGAGTATGGTTGCAAAGCTGAAACTTAAAGGAATTGACGGAAGGGCACCAGGAGTGAGCCTGCGGCTTAATTTGACTCAGCACGGGAACCTCACCGGCCCGGACACGGAAAGGATTGACAGATTGATAGCTCTT

**>R2Dr-Retrovirus_upstream-part**

CGGAGGATGACCCGTGGATGGGTGAAATATAGAGTCGGGGTCACTCTGGTGAGGCCTGGCTAAAACTGATCGATAGGATGATTATGCCCCTGGGGTAGGCAGCTTCCCGGATGACGTCAAACTCATCCGTTATGGGATGATCCAGAACCTCCCGATTTAGGCCCCTGCTGAGC

**>R2Dr-Retrovirus_(AB097126.1)_4,000bp**

AATCCCCCTACCCAATCCCCCGTCGTGACCTCCAGGCCAGGAATCACGGGCGTCGACAGTGGCCATCCGGCAATGACAATAAACGTGACTAACGACAATGAGTTAGATCCATGACCCTTGGAGTGGGTTTAACCTCCGCCTCTTTAAAAACATGGAAAGTACAGCAAAAAGGAAAGTCATACTGGATGACCCCCGTCGCCCAGTAGAAAGGTGCCACGGAGGGATCTTTGGGTCGGGTCCCTTTCGTAACGCGAGATCTAAGCGCAAACCAGAGGCTAAACGAACACTACATGGCTTAGGACTACGAGAATGCTCGGTTGTCTTGACGCGCCTCAGCGGGGCGTCGAGGTCGTGATCAGACACCAGCAGGATGGAACGCACAACGCGGCAGCCCAAACGACGAGCTCGGTTGAAGGCCCAATGGGCCGATACCGTAACCCCATACCAACGGGCACCCAAGCCCTGCCTGAACCTATGGCGGGCCCGGGAGCAGGGGAGCACCCGGGGAGTGGTGGATGATCCTGCCGCTCCGGGACTTCAACTGCCCCCTATGTGGCGGGTCGGCGAAACGCAGCGGTGGAAGTGCAAAGACACTTTGCATTTCGCCACGGAACAGTGCCCGTTAGATTCCAGCTGTGAATCATGTGGAAAAACCTCTCGGTTGCCATTCCGTCCTCTGTCACATTCCCGAAATATCGCGGACCGACAGGCGAGCCGCCTGAAAGTGGTTAGTGCGGGGATGCAATAGGACATTTGACACAAGGAGGCGTGGTAGCCACATGAGATGCACGTCCACTCAGAAATCCGCAATAGGAAAAGAATTGCTCAAGACAGGCAAGAAAAGGGACCTCGACAGATGGAGAGGAGAGCTATGGGAGAATCGAAAGGGCTGACGCTGAGGAAGGTCCCTCTGGGGAAGGGATCCCCCTAACGTCCCAGGCGTGCGAGAACGCCCAGAGAACCGTCCGAGCCCCAGCAAATCCACCGATCCTCTCACCACAGCCCAATCTGCCGGGAGGAGCCTCCCGGGACCAGCTCCGGGAGGTGGCCAGTAGATGGGTAAGGGCGACGAGACGGGTACGGTGATCGACAGCGTTCTCGCTGCATGGTTGGATGGCAACGATCGGCTCCCTGAGCTGGTTGGCGCGACGCAAAAAGGACACTGCAGGGCCTACCTGCAGGAGGTTGACCCGAAGGCCCGTAGCTTTGTTGCGCCTAAGCGGAGGAGAGGCAGAGTGGGGACGCCGGCTCAAATTACTCGGCGCTAACGCCGCGCCTGCACCACAACTGCCCAGAGTCGGTTCCGGAAAGGCCCAGCCCGCCTGGCCGCGAACATCCTAGACGGCAAAAGTGAAACAAAGTGCCCAATCACAGGAGAGGTGATTTACGAACACTTTAAAAGCAAATGGCGAATCCAAGACCCTTTGCCGGGCTGGGGCGATTTGGGGCGGAAGAAAGGGCAAACAACGCCCACCTCCTCGGAGATCTCCATAGCGAGGTCCAAAATAGCCTCCGACATACATCGAATACATCCGCACGGGCCCAGATGGCGTTGGAAGGACATTTCCAGCTGGGACCCCGAATGCGAGACCCTCCTTTGGCTGTTTCAACATGTGGTGGTTCACAGGTGTCATTGAGCAGCTTGAAGAAAGGGTCGAACGGTGCTACTGCCCAAGACCTCTCGGACCAGGGGCGCGATGGAGGGTGGCTGGAGACCAATCACCATCGGGTCGATGGTCTTGCGGCTTTTCACGAGGGTGATCAACACAAGATTGACAGAGGCGTGCCCGTTGCACCCAGACAGAGAGGGTTTCGGCAGAGCCCCGGGTGCTCAGAAAGAGGCCTGAAGCTGCTCCAATCGTTCTCCGTCACTCCAAAAGAAAAGAGCGCAGCCAACTGGCGGTAGTGTTTGTCGATTTGCACAAGCGTTCGACACCGTCTCTCATGAACACTTGCTGTCGGTTCTTGAACGGATGAGCGTGGACCCCACATGGCTAAATCTGATCCAGATTTCCAAACAGCTGCACAAGTGTCGAGTTAGGCCGAAGGGAGGGACCAGATATCCCAGTGAGGGTTGGTGTCAAGCAGAAGTCCACTGTCCCGTTGCTTTTCAACCTGGCTTTGGATCCCTCGGCTCAAAGTCACCGAGCGCGCACAGGCAAGGTATGAGGTGGAAAGTGCAAAGTGACAGCTTTAGCGTTCGCGGATGACCTGGCACTGGTTGCGAACTCGTGGGAGGGAATGGCACACAACCTTGCGCTTGTAGACGAATTCTGCCTAACCACCGGCCTCACAGTCCAACCCAAAAAGTGCCACAGTTTCATGGTCAGGCCCTGCAGAGGTGCCTTCACAGTGAGCGACTGCCCCCATGGGTTCTGGGGGCAAGGCCCTGCAGCTAACGAACATCGAAAACTCCATCAAATATCTGGGAGTAAAAGTCAATCCTTGGGCGGGGATTGAAAAGCCTGACCTTACAGTGGCAACTGGGCCGATGGTGCAGCGCATTGGGAAGTCACTGCTCAAACCCTCACAGAAGGTATACATTCTCAATCAGTTTGCCATCATGACTCTTCTACCTGGCTGATCACGGTGGGGCGGACGTCATGCTCAGAACCTGGATGGGACAATCAGGAAGGCGGTGAAGAAATGGCTGCATCTTCCGTCAGCCTGGGCTGTTGTATGCAGGGACTGTAATGGTGGCCTCGGTATATGCAAGCTCACTTTCATTACATCCCATCAATGCAGGCGAGGCAGATATTCCGCTTGGCCAACTCCGCGGACAACCCGTTGATGAAGGCCATGATGCGCGGCTCCCGGAGTCGAACAGAAATTCAAAAAAGGCCTGGATGCGGGCCGGGGAGAGGAGAGTGCGCTCCACGGGTGTTCGGGGCGGAATCCAGTACCAGGAAGGGAGGAGGGTCGCTAACGATCTGGTACCTCGCTGCCCAATGCCGGCGATTGGAGACTGGAAGAATTCCAGCACTGGATGGGCCTGCGGAGTCCAGGGTGTGGGGTATGCTTGGCTTCTTCAGAAACAAGGTGGCTAACGGATGGCTCAGGAAGCCGGCAGGGTTCAAGAGCGGCACTACATCGCCGCTCTACAACTGCGAGCATGTGTATACCCCACCCTGTGAATTCCAGCAAAGGGGCAGGAGCAAAGCGGGTGCGACCTGCAGGCGGTGCTCATCCCGGTTAGGTCCAGCTCTCACATCCTCGGCAAATGTCGGTGCAGGGAGCCAGAATCAAGGCGTCACAACAAAATATGCGACCTCCTGAAGGCGAAGCGAAACCCGGGGTTGGGAAGTACGCCAGGAATGGGCCTTGGAACTCCGGCTAGGGAACTGAAGGCTCGACCTGGTACTCATCCTCGGGGATGAGGCATTAGTCATTGACGTCACAGTAGGTACGAGTTGTAGGATACCCTCCAGAATGCCGGAAAGGACAAGGTCCCAGCTACTACGGCCCGCACAAAGAAAGCGATCGCTCAGGCTGGGCGTAGAAAAGGTCGACATACATGGGTTCCCGTTGGGTGCACGCGGACTTTGGCTGCACTAGCAACTCCAAAGTGCTGGAACTGATGGGATTAGCAGGAAAGAGTGAAGGTCTTCTCAGACTCTTGAGTCGGAGGGTGCTCCTGTACTCTATCGACATCATGAGGACATTTTACGCATCCCTGCAATGAAAATCCCAGCGGGATACAGCAAGAAGGTATCGGATCTAATAAGGTTGAGCGAGGAGAGGTGGAGATCCTTTGGGGGTCGGGCGGATTCCCTCTCGGGTCCTCCCCACGGTGACGCTCTGCCCCTCCTCTCGCTCGTAGGAGCCCAGCGGTGAACACGGTTGGCAGGATGAGTGACGTGAGGGGTAAGACATGCGTACGTGAGCGCGCATTTTGCTGTTCTCTGGACTGGGTTTCGTCTCCCCCTCACAACCATCTTACACTATAGGGGCACAGCGGCTCCTACCTCCTCCCTATGACCCCCTCCCATACCGATCCATG
