## Supplemental File SF4 for "Immunoglobulin switch-like recombination regions implicated in the formation of extrachromosomal circular 45S rDNA involved in the maternal-specific translation system of zebrafish"

### Supplemental File SF4: 45S-M Repeat region, genomic sequence

**45S-M: 5x90nt GCA repeat**

***Repeat 1***

TGGG**TA**ACCAGCAGCACGCAGTACCTAT**A**TGGGGTAACCAGC**A**GCAGAAGGG**A**TTC**A**GTCCC**A**ACCCGCACCCCACAGCA**G**AGCC**TT**TA**T**

TGGGGCACCAGCAGCACGCAGTACCTATGTGGGGTAACCAGCGGCAGAAGGGGTTC**T**GTCCCGACCCGCACCCCACAGCAAAGCC**CC**TA**C**

TGGGGCACCAGCAGCACGCAGTACCTATGTGGGGTAACCAGCGGCAGAAGGGGTTC**T**GTCCCGACCCGCACCCCACAGCAAAGCC**CC**TA**C**

TGGGGCACCAGCAGCACGCAGTACC**C**A**C**GTGGG**-**T**CG**CCAGCGGCAGAAGGGGTCC**A**G**C**C**T**CG**C**CCCGCACCCCACAGCAAAGCC**TT**TA**G**

TGGGGCACCAGCAGCACGCAGTACCTA**CA**TGGGGTTACCAGCGGCAGAAG**A**GGTCC**G**G**A**CCCGACCC**A**CACCCCACAGCAAAGCC**TTCGA**

TGGGG**T**ACCAGCAGCACGCAGTAC**A**TA**AA**TGGG**-**T**C**ACCAGCGGC**GCGT**G**A**G**TCTGGCG**C**G**CG**CG**CC**TT**ACC**GG**A**GTCAGTGT**CC**CC**G**GG**

***Repeat 2***

```
                                                                CCCGCACCCCACAGCAAAGCCCCTAC
TGGGGCACCAGCAGCACGCAGTACCTATGTGGGGTAACCAGCGGCAGAAGGGGTTCTGTCCCGACCCTAACCCCACAGCAGAGCCTTTAT
TGGGGCACCAGCAGCACGCAGTACCCACGTGGG-TCGCCAGCGGCAGAAGGGGTCCAGCCTCGCCCCGCACCCCACAGCAAAGCCTTTAG
TGGGGCACCAGCAGCACGCAGTACCTACATGGGGTTACCAGCGGCAGAAGAGGTCCGGACCCGACCCACACCCCACAACAAAGCCTTCGA
TGGGGTACCAGCAGCACGCAGTACATAAATGGG-TCACCAGCGGCGCGTGAGTCTGGCGCGCGCGTCTTACCGAAGTCAGTGTCCCCGGG
```

***Red****;* *non-consensus nucleotides*

***Blue****; nucleotides without consensus
Yellow: GCA trimer*

***Green****: GGG trimer*

***-****: missing nucleotide*

*Boxed: different nucleotide as compared to other repeat*

**45S-M: Ig switch-like region** (176x4nt CCAG repeat)

***Region 1***

CCAGCCAGCCAGTCAGTCAGCCCGGCCCAGCCCAGCCCAGCCCAGCCAGCCAGCCAGCCAGCCAGCCAGCCCAGCCCAGCGCTGTCCAGCCAGCCCTGTCCAGCCCAGCCCAGGCAAGCCCAGGCCGGCCGGCCAGCCAGCCAGCCAGCAAGCCCAGGCCAGGCCAGGCCCAGCCAGCCCAGCCAGCCCTGCCCTGCCCTGCCCTGCCCTGCCCAGCCCAGCCCAGGCCGGCCAGCCAGCCAGCCAGCCAGCCAGCCAGCCAGCCCAGGCCAGGCCAGGCCAGGCCAGGCCAGGCCAGGCCAGGCCAGGCCAGGCCCAGCCCGGCCAGGCCAGGCCCAGCCCAGCCAGCCCAGTCAGCCAGCCAGCCCAGCCCAGCCAGCCAGCCAGCCAGCCAGCCAGCCCAGCCCAGCCCAGCCCAGGCAAGCCCAGGCCGGCCAGCCAGCCAGCCAGCCAAGGCCAGGCCAGGCCCAGCCCAGCCAGCCAGCCAGCCAGCCCAGGCAAGGCCAGGCCAGGCCAGGCCAGGCCAGGCCAGGCCAGGCCAGGCCAGGCCAGGCCCAGCCCAGCCAGGCCAGCCCAGCCGAGCCCAGCCAGCCAGCCAGCCAGCCAGCCCAGCCCAGCCCAGCCCAGCCCTGCCCTGCCCTGCCCTGCCCTGCCCTGCCCAGCCCAGCCCAGCCCAGCCCAGCCAAGCCCAGGCCGGCCAGCCAGCCAGCCAGCCAGCCAGCCCAGGCCAGGCCAGGCCAGGCCAGGCCAGGCCAGGCCAGGCCCAGCCCAGCCAGCCAGCCAGCCAGCCAGCCAGCCAGCCAGCCCAGGCCAGGCCAGGCCAGGCCATGCCCAGCCCAGCCAGCCAGCCAGCCCAGGCCAAGCAAGCCCTGGCTGGGCAGCAAGCCCAGCTCAGCCAGCCAGCCCAGGCCAGGCAAGCCCTGGCCGGCCAGCCAGCCAAGCCCAGCCAGCCAATCAGTCAGTCAAGCCAAGGCCAGTCAGTCAGGTCAATCAGCCCAGCCCAGCCCAGCCTAGCCTAGCCTAGCCCAGCCCAGCCCAGCCCAGCCCAGCCCAGCCCTGCCCTGCCCAACCTGACAGCCAGGCCTGCTCAGCCAGCCAGCCAGCCA

***Region 2***

CCAGCCAGCCAGTCAGTCAGCCCGGCCCAGCCCAGCCCAGCCCAGCCAGCCAGCCAGCCAGCCAGCCAGCCCAGCCCAGCGCTGTCCAGCCAGCCCTGTCCAGCCCAGCCCAGGCAAGCCCAGGCCGGCCGGCCAGCCAGCCAGCCAGCAAGCCCAGGCCAGGCCAGGCCCAGCCAGCCCAGCCAGCCCTGCCCTGCCCTGCCCTGCCCTGCCCAGCCCAGCCCAGGCCGGCCAGCCAGCCAGCCAGCCAGCCAGCCAGCCAGCCCAGGCCAGGCCAGGCCAGGCCAGGCCAGGCCAGGCCAGGCCAGGCCAGGCCCAGCCCGGCCAGGCCAGGCCCAGCCCAGCCAGCCCAGTCAGCCAGCCAGCCCAGCCCAGCCAGCCAGCCAGCCAGCCAGCCAGCCCAGCCCAGCCCAGCCCAGGCAAGCCCAGGCCGGCCAGCCAGCCAGCCAGCCAAGGCCAGGCCAGGCCCAGCCCAGCCAGCCAGCCAGCCAGCCCAGGCAAGGCCAGGCCAGGCCAGGCCAGGCCAGGCCAGGCCAGGCCAGGCCAGGCCAGGCCCAGCCCAGCCAGGCCAGCCCAGCCGAGCCCAGCCAGCCAGCCAGCCAGCCAGCCCAGCCCAGCCCAGCCCAGCCCTGCCCTGCCCTGCCCTGCCCTGCCCTGCCCAGCCCAGCCCAGCCCAGCCCAGCCAAGCCCAGGCCGGCCAGCCAGCCAGCCAGCCAGCCAGCCCAGGCCAGGCCAGGCCAGGCCAGGCCAGGCCAGGCCAGGCCCAGCCCAGCCAGCCAG**--------**CCA GCCAGCCAGCCAGCCCAGGCCAGGCCAGGCCAGGCCATGCCCAGCCCAGCCAGCCAGCCAGCCCAGGCCAAGCAAGCCCTGGCTGGGCAGCAAGCCCAGCTCAGCCAGCCAGCCCAGGCCAGGCAAGCCCTGGCCGGCCAGCCAGCCAAGCCCAGCCAGCCAATCAGTCAGTCAAGCCAAGGCCAGTCAGTCAGGTCAATCAGCCCAGCCCAGCCCAGCCTAGCCTAGCCTAGCCCAGCCCAGCCCAGCCCAGCCCAGCCCAGCCCTGCCCTGCCCAACCTGACAGCCAGGCCTGCTCAGCCAGCCAGCCAGCCA

*Yellow: CCAG tetramer*

***Blue****: CCCTG aptamer*

***-****: missing nucleotide*

*Boxed: different nucleotide as compared to other region*

**45S-M: 16x3nt ACC repeat**

***Repeat 1***

ACCACCACCACCACCACCACCACCACCACCACCACCACCACCACCACC

***Repeat 2***

*100% identical to Repeat 1*

*Yellow: ACC trimer*

**45S-M: 21x72nt GGAGG repeat**

***Repeat 1***

C**CGGAGG**CCCGGGGATTGGGCTTGGG**G**ATAGGGTAAGGGTTGCCT**GGAGG**CTTGGGGGTGGGTTAGGGCTGC

C**CGGAGG**CCCGGGGATTGGGCTTGGG**G**ATAGGGTAAGGGTTGCCT**GGAGG**CTTGGGGGTGGGTTAGGGCTGC

C**CGGAGG**CCCGGGGATTGGGCTTGGG**G**ATAGGGTAAGGGTTGCCT**GGAGG**CTTGGGGGTGGGTTAGGGCTGC

C**AGGAGG**CCCGGGGATTGGGCTTGGG**C**ATAGGGTAAGGGTTGCCT**GGAGG**CTTGGGGGTGGGTTAGGGCTGC

C**TGGAGG**CCCGGGGATTGGGCTTGGG**G**ATAGGGTAAGGGTTGCCT**GGAGG**CTTGGGGGTGGGTTAGGGCTGC

C**AGGAGG**CCCGGGGATTGGGCTTGGG**C**ATAGGGTAAGGGTTGCCT**GGAGG**CTTGGGGGTGGGTTAGGGCTGC

C**TGGAGG**CCCGGGGATTGGGCTTGGG**C**ATAGGGTAAGGGTTGCCT**GGAGG**CTTGGGGGTGGGTTAGGGCTGC

C**TGGAGG**CCCGGGGATTGGGCTTGGG**G**ATAGGGTAAGGGTTGCCT**GGAGG**CTTGGGGGTGGGTTAGGGCTGC

C**AGGAGG**CCCGGGGATTGGGCTTGGG**C**ATAGGGTAAGGGTTGCCT**GGAGG**CTTGGGGGTGGGTTAGGGCTGC

C**TGGAGG**CCCGGGGATTGGGCTTGGG**G**ATAGGGTAAGGGTTGCCT**GGAGG**CTTGGGGGTGGGTTAGGGCTGC

C**AGGAGG**CCCGGGGATTGGGCTTGGG**C**ATAGGGTAAGGGTTGCCT**GGAGG**CTTGGGGGTGGGTTAGGGCTGC

C**TGGAGG**CCCGGGGATTGGGCTTGGG**G**ATAGGGTAAGGGTTGCCT**GGAGG**CTTGGGGGTGGGTTAGGGCTGC

C**TGGAGG**CCCGGGGATTGGGCTTGGG**G**ATAGGGTAAGGGTTGCCT**GGAGG**CTTGGGGGTGGGTTAGGGCTGC

C**AGGAGG**CCCGGGGATTGGGCTTGGG**C**ATAGGGTAAGGGTTGCCT**GGAGG**CTTGGGGGTGGGTTAGGGCTGC

C**TGGAGG**CCCGGGGATTGGGCTTGGG**G**ATAGGGTAAGGGTTGCCT**GGAGG**CTTGGGGGTGGGTTAGGGCTGC

C**AGGAGG**CCCGGGGATTGGGCTTGGG**C**ATAGGGTAAGGGTTGCCT**GGAGG**CTTGGGGGTGGGTTAGGGCTGC

C**AGGAGG**CCCGGGGATTGGGCTTGGG**G**ATAGGGTAAGGGTTGCCT**GGAGG**CTTGGGGGTGGGTTAGGGCTGC

C**AGGAGG**CCCGGGGATTGGGCTTG**A**G**C**ATAGGGTAAGGGTTGCCT**GGAGG**CTTGGGGG**GG**T**TA**GGGTTAGGGTTGC

C**CGGAGG**CC**T**GGGGATTG**A**GCTTGGG**G**ATAGGGTAAGGGTTGCCT**GGAGG**CTTGGGTGT**--TA**GGGTTAGGG**T**T**T**C

C**CGGAGGA**C**T**GGG**A**ATT**-------------**GGGT**T**AGGGTTGCCT**GGAGA**CTTGGGGGT**---G**GGGTTAGGGCTGC

C**CGGAGG**CC**T**GGGGATTGGGCTTGGG**G**ATAGGGT**T**AGGGTTGCC**CGGAGG**CTT**A**G**TT**G**T---T**GGG**C**T**T**G**T**G

***Repeat 2***

*100% identical to Repeat 1*

***Red****;* *non-consensus nucleotides*

***Blue****; nucleotides without consensus
Yellow: GGAGG pentamer*

***Green****: GGG trimer*

***-****: missing nucleotide*

**45S-U: 10x212nt TGG repeat**

***Repeat U1 upstream***

*100% identical to Repeat U1-U2, but only the first 7 short repeats*

***Repeat U1-U2***

CTG**C**GCCCCGCAAGCCGGCTC**C**GGTCGTCCGTTTTGGGCATGGTGC**---**ATAGATTGAGTGTGGGTTGGTGGTGTCCGGTCAACTCCGTGGCTATAGATCA**T**CGGGACATGTCTGC

CAATCCCTGCCCTGGCTTGCACACTTTCCCCCGGGTATGCATAGCCCCGTGGCTCGGTGAAGTGGAAGTTCTGGTGCTTCAATAGCGACGTTGTGG

CTGGGCCCCGCAAGCCGGCTCTGGTCGTCCGTTTTGGGCATGGTGCTGCATAGATTGAGTGTGGGTTGGTGGTG**A**CCGGTCAACTCCGTGGCTATAGATCA**C**CGGGACATGTCTGCCAATCCCTGCCCTGGCTTGCACACTTTCCCCCGGGTATGCATAGCCCCGTGGCTCGGTGAAGTGGAAGTTCTGGTGCTTCAATAGCGACGTTGTGG

CTGGGCCCCGCAAGCCGGCTCTGGTCGTCCGTTTTGGGCATGGTGCTGCATAGATTGAGTGTGGGTTGGTGGTGTCCGGTCAACTCCGTGGCTATAGATCATCGGGACATGTCTGCCAATCCCTGCCCTGGCTTGCACACTTTCCCCCGGGTATGCATAGCCCCGTGGCTCGGTGAAGTGGAAGTTCTGGTGCTTCAATAGCGACGTTGTGG

CTGGGCCCCGCAAGCCGGCTCTGGTCGTCCGTTTTGGGCATGGTGCTGCATAGATTGAGTGTGGGTTGGTGGTG**A**CCGGTCAACTCCGTGGCTATAGATCA**C**CGGGACATGTCTGCCAATCCCTGCCCTGGCTTGCACACTTTCCCCCGGGTATGCATAGCCCCGTGGCTCGGTGAAGTGGAAGTTCTGGTGCTTCAATAGCGACGTTGTGG

CTGGGCCCCGCAAGCCGGCTCTGGTCGTCCGTTTTGGGCATGGTGCTGCATAGATTGAGTGTGGGTTGGTGGTG**A**CCGGTCAACTCCGTGGCTATAGATCATCGGGACATGTCTGCCAATCCCTGCCCTGGCTTGCACACTTTCCCCCGGGTATGCATAGCCCCGTGGCTCGGTGAAGTGGAAGTTCTGGTGCTTCAATAGCGACGTTGTGG

CTGGGCCCCGCAAGCCGGCTCTGGTCGTCCGTTTTGGGCATGGTGCTGCATAGATTGAGTGTGGGTTGGTGGTGTCCGGTCAACTCCGTGGCTATAGATCATCGGGACATGTCTGCCAATCCCTGCCCTGGCTTGCACACTTTCCCCCGGGTATGCATAGCCCCGTGGCTCGGTGAAGTGGAAGTTCTGGTGCTTCAATAGCGACGTTGTGG

CTGGGCCCCGCAAGCCGGCTCTGGTCGTCCGTTTTGGGCATGGTGCTGCATAGATTGAGTGTGGGTTGGTGGTGTCCGGTCAACTCCGTGGCTATAGATCATCGGGACATGTCTGCCAATCCCTGCCCTGGCTTGCACACTTTCCCCCGGGTATGCATAGCCCCGTGGCTCGGTGAAGTGGAAGTTCTGGTGCTTCAATAGCGACGTTGTGG

CTGGGCCCCGCAAGCCGGCTCTGGTCGTCCGTTTTGGGCATGGTGCTGCATAGATTGAGTGTGGGTTGGTGGTGTCCGGTCAACTCCGTGGCTATAGATCATCGGGACATGTCTGCCAATCCCTGCCCTGGCTTGCACACTTTCCCCCGGGTATGCATAGCCCCGTGGCTCGGTGAAGTGGAAGTTCTGGTGCTTCAATAGCGACGTTGTGG

CTGGGCCCCGCAAGCCGGCTCTGGTCGTCCGTTTTGGGCATGGTGCTGCATAGATTGAGTGTGGGTTGGTGGTGTCCGGTCAACTCCGTGGCTATAGATCATCGGGACATGTCTGCCAATCCCTGCCCTGGCTTGCACACTTTCCCCCGGGTATGCATAGCCCCGTGGCTCGGTGAAGTGGAAGTTCTGGTGCTTCAATAGCGACGTTGTGG

CTGGGCCCCGCAAGCCGGCTCTGGTCGTCCGTTTTGGGCATGGTGCTGCATAGATTGAGTGTGGGTTGGTGGTG**A**CCGGTCAACTCCGTGGCTATAGATCA**C**CGGGACATGTCTGCCAATCCCTGCCCTGGCTTGCACACTTTCCCC**T**GGGT**G**TGCATTGCCCCGTGGCTCGGTGAAGTGCAAGTTCTGGTGCTTCAATAGCG**G**CGTTGTGG

***Repeat U2 downstream***

*100% identical to Repeat U1-U2, but only the first 7 short repeats*

***Red****;* *non-consensus nucleotides*

***Yellow****: TGG trimer*

***-****: missing nucleotide*

**45S-U 4x78nt CCA repeat**

***Repeat U1 upstream***

AAACTCCAGTAGTGACAGCCAGGTGGGTCACCAGCAGCACACAATACTCAATTGGGCC

ACCAGCAGCACGCAATACTCAATTGGGCCACTAGCGTCACACAATACTCGATTGGGTCACCAGCGGCACGCAGCCCCC

AAATCCAAATAGTAAAGGCAAGGTGGGCAACCAGCGTCAAACAATACTCGATTGGGTCACCAGCGGCACGCAGCCCCC

AAATCCAAATAGTAAAGGCGAGGTGGGCAACCAGCGTCAAACAATACTCGATTGGGTCACCAGCGGCACGCAGCCCTC

AAATCCAAATAGTAAAGGCCAAGAGGGCCACCAGCGGCACGCATTACTGAAATGGGTCACCAGCAGCACGTGAGTCTG

***Repeat U1-U2***

*100% identical*

***Repeat U2 downstream***

*100% identical*

*Yellow: CCA trimer*

*Green: GGG trimer*

**45S-U: Ig switch-like region**

***Repeat U1 upstream***

CAAGCCCAGGCCGGCCAGCCAGCCAGCCAGCCAGCCAGCCACCCCCAATCCAGCCAGGCCAGGTCAGGTCAGGCAAGCCCAGGCCGGCCAGCCAGCCAGCCAGCCAGCCACCCCCAGCCCAGCCAGGCCAGGCCAGGTCAGGCAAGCCCAGGCCGGCCAGCCAGCCAGCCAGCCAGCCACCCCCAGCCCAGCCAGGCCAGGCCAGGTCAGGCAAGCCCAGGCCGGCCAGCCAGCCAGCCAGCCAGCCAGGCCAGGCCAGGTCAGGCAAGCCCAGGCCGGCCAGCCAGCCAGCCAGCCAGCCACCCCCAATCCAGCCAGGCCAGGCCAGGTCAGGCAAGCCCAGGCCGGCCAGCCAGCCAGCCAGCCAGCCAGCCACCCCCAATCCAGCCAGGCCAGGCCAGGTCAGGCAAGCCCAGGCCGGCCAGCCAGCCAGCCAGCCAAGTCCAGCCAAGCCCACCCCCAGTCAGCCAGCCAGCCAGCCAGCCAGCCAGCCAGGCCAGGCCAGCCAAGCCCAGCCCAGCCCACCCCCAGTCAGCCAGCCAGCCAGGCCAGGCCAGGCCAGCCAAGCCCAGCCCAGCCCACCCCCAGTCAGCCAGCCAGCCAGCCAGCCCAGGCCAGCCAAGCCCACCCCCAGTCAGCCAGCCAGCCAGCCAGCCAAGCCCACCCCCAGTCAGCCAGCCAGCCAGCCAGCCAGGCCAGGCCAGGCCAGGCCAGCCAAGCCCAGCCCAGCCCAGCCCAGCCCACCCTCAGTCAGCCAGCCAGCCAGCCAGGCCAGGCCAGGCCAGCCAAGCCCAGCCCCAGTCAGCCAGCCAGCCAGCCAGCCAGCCCAGGCCAGCCAAGCCCACCCCCAGTCAGCCAGCCAGCCAGCCAGCCAGCCCAGGCCAGCCCAGCCCACCCCCAGTCAGCCAGCCAGCCAGCCAGCCAGCCCAGGCCAGCCAAGCCCACCCCCAGTCAGCCAGCCAGCCAGCCAGCCAGCCAGCCAGGGCAGGCCAGGCCAGCCCAGCCCAGCCCAGCCCACCCCCAGTCAGCCAGCCAGCCAGCCAGGCCAGGCCAGGCCAGGCCAGCCAAGCCCAGCCCACCCCCAGTCAGCCAGCCAGCCAGCCAGCCAGCCCAGGCCAGCCAAGCCCACCCCCAGTCAGCCAGCCAGCCAGCCAGCCAGCCCAGGCCAGCCAAGCCCACCCCCAGTCAGCCAGCCAGCCAGCCAGCCAGGCCAGGCCAGGCCAGCCAAGCCCAGCCCACCCCCAGTCAGCCAGCCAGCCAGCCAGCCAGCCAAGCCCACCCCCAGTCAGCCAGCCAGCCAGCCAGGCCAGGCCAGGCCAGGCCAGGCCAGCCAAGCCCAGCCCACCCCCAGTCAGCCAGCCAGCCAGCCAGCCAGCCCAGGCCAGCCAAGCCCACCCCCAGTCAGCCAGCCAGCCAGCCAGCCAGGCCAGGCCAGGCCAGGCCAGGCCAGCCAAGCCCAGCCCAGCCCAGCCCAGCCCAGCCCAGCCCAGCCCACCCTCAGTCAGCCAGCCAGGCCAGCCAAGCCCAGCCCACCCCCAGTCAGCCAGCCAGCCAGCCAGCCAGCCCAGGCCAGCCAAGCCCACCCCCAGTCAGCCAGCCAGCCAGCCAGCCAGCCAGCCAAGCCCACCCCCAGTCAGCCAGCCAGCCAGCCAGCCAGCCAGCCAGGCCAGGCCAGGCCAGGCCAGGCCAGGCCAGCCCAGCCCAGCCCAGCCCACCCCCAGTCAGCCAGCCAGCCAGCCAGGCCAGGCCAGGCCAGCCAAGCCCAGCCCACCCCCAGTCAGCCAGCCAGCCAGCCAGCCAGCCAGCCAAGCCCACCCCCAGTCAGCCAGCCAGCCAGCCAGCCAGCCAGCCAAGCCCACCCCCAGTCAGCCAGCCAGCCAGCCAGGCCAGGCCAGGCCAGGCCAGGCCAGCCAAGCCCAGCCCAGCCCAGCCCAGCCCAGCCCAGCCCACCCCCAGTCAGCCAGCCAGCCAGCCAGGCCAGGCCAGGCCAGCCAAGCCCAGCCCACCCCCAGTCAGCCAGCCAGCCAGCCAGCCAGCCCAGGCCAGCCAAGCCCACCCCCAGTCAGCCAGCCAGCCAGCCAGCCAGCCAGCCAGGCCAGGCCAGGCCAGCCCAGCCCAGCCCACCCCAGTCAGCCAGCCAGCCAGCCAGGCCAGGCCAGGCCAGGCCAGGCCAGGCCAGGCCAGGCCAGGCCAGCCAAGCCCAGCCCACCCCCAGTCAGCCAGCCAGCCAGCCAGCCAGCCAGCCAGCCCAGGCCAGCCAAGCCCACCCCCAGTCAGCCAGCCAGCCAGCCAGCCAGCCAGCCAGGCCAGGCCAGGCCAGGCCAGGCCAGCCCAGCCCAGCCCAGCCCAGCCCCAGTCAGCCAGCCAGCCAGCCAGCCAGCCAGCCAGGCCAGGCCAGCCCAGCCCAGCCCAGCCCAGCCCAGCCCAGGCCAGCCAAGCCAGCCCACCCCCAGTCAGCCAGCCAGCCAGCCAGCCAGGCCAGGCCAGGCCAGCCAAGCCCAGCCCAGCCCACCCCCAGTCAGCCAGCCAGCCAGCCAGCCCAGGCCAGCCAAGCCCACCCCCAGTCAGCCAGCCAGCCAGCCAGGCCAGGCCAGCCCAGCCCAGCCCAGCCAGCCAGCCAGCCAGCCCAGGCCAGCCAAGCCAGCCCAGGCCAACCCAACCAGCCCAGGCCAGCCAAGCCAGCCCACCCCCAGTCAGCCAGCCAGCCAGCCAGCCAGGCCAG

***Repeat U1-U2***

*100% identical to Repeat U1 upstream*

***Repeat U2 downstream***

*100% identical to Repeat U1 upstream*

*Yellow: CCAG tetramer*

**45S-S: 7x300nt TGGG repeat**

***Repeat 1 upstream***

TGGGTTTTATCTTGCTCTTTGGAACACTGCACAAGCCTCGTCAAGGCCCAGTCCTCCAACTCTCACGCTGGGGGATGGCTTTCCTGCTCTCTGTCTAGGGGTGGTGGGTTTCCTCCATTTTTGTCAAAGTCAAGCCGATTCAGCTCACCCTGTAAGAAAGGGGGGGTTAGAAAGAGAGAATAGGCAAAGAGTCACTGGTCGTTCCAACACTCACTTGGAGGATGGTGGGTCTGTCCTTTTTTAGAGAGTCAAGCCGATTCCCCTCACTCTCAAGAGAGGCACTGTACAACCAATAG

TCACTTGGGGGATGGTGGGTCTTATCTTGCTCTTTGGAACAGTGCACAAGACTCGTCGAGGCCCACTCCTCCAACTCTCACGCTGGGGGATGGCTTTCCTGCTCTCTGTCTAGGGGTGGTGGGTTTCCTTCATTTTTGTCAAAGTCAAGCCGATTCAGCTCACCCTGTAAGAAAGGGGGGGGTTAGAAAGAGAGAATAGGCAAAGAGTCACTGGTGGTTCCAAGACAAGCCCCCAACACTCACTTGGGGGATGGTGGGTCTGTCCTCTTTTTGTAGAGTCAAGCCGATTCCCCTCACTCTCAAGAGAAAGCGAGCCACTGGTCATCCACAC

TCACTTGGGGGATGGTGGGTCTTATCTTGCTCTTTGGAACAGTGCACAAGACTCGTCGAGGCCCACTCCTCCAACTCTCACGCTGGGGGATAGGTTTCCTGCTCTCTGTCTATGGGTGGTGGGTTTCCTCCATTTTTGTCAAAGTCAAGCCGATTCAGCTCACCCTGTAAGAAGGAGAGAGAAGAAAGAGAAAGAGAGAAAAGGCAAAGAGTCACTGGTGGTTCCAACACTCACTTGGGGGATGGTGGGTCTGTCCTCTTTTTGTAGAGTCAAGCCGATTCCCCTCACTCTCAAGAGAAAGCGAGCCACTGGTCATCCAACAC

TCACTTGGGGGATGGTGGGTCTTATCTTGCTCTTTGGAACACTGCACAAGTCTCGTCAAGGCCCACTCCTCCAACTCTCACGCTGGGGGATGGCTTTCCTGCTCTCTGTCTATGGGTGGTGGGTTTCCTCCATTTTTGTCAAAGTCAAGCCGATTCAGCTCACCCTGTAAGAAGGGGGGGGGGGGGGGGGTTTAGAAAGAGAGAAAAGGCAAAGAGTCACTGGTGGTTCCAACACTCACTTGGGGGATGGTGGGTCTGTCCTCTTTTTGTAGAGTCAAGCCGATTCCCCTCACTCTCAAGAGAAAGCGAGCCACTGGTCATCCACAC

TCACTTGGGGGATGGTGGGTCTTATCTTGCTCTTTGGAACAGTGCACAAGACTCGTCGAGGCCCACTCCTCCAACTCTCACGCTGGGGGATAGGTTTCCTGCTCTCTGTCTATGGGTGGTGGGTTTCCTCCATTTTTGTCAAAGTCAAGCCGATTCAGCTCACCCTGTAAGAAAGGGGGGGTTAGAAAGAGAGAATAGGCAAAGAGTCACTGGTGGTTCCAACAC

TCACTTGGGGGATGGTGGGTCTTTTCTTGCTCTTTGGAACACTGCACGAGCCTCGTCGAGGCCCACTGCTCCAACTCTCACTCGGCGCATGACTTTCCTGTTCTCTGTTTAGGAGTGGTGGGTTTCCTTCGTTTTTGTCAAAGTCAAGCCGATTCAGCTCATCCTGCAAGAAAGAGAGAGGAGGGGAAGGGAGGGAGAGACAGAGAGAGAGAGAGAGAGCGAGAGAGAGAGAGAGAGAGAGGCACAGAGGCCATGGTTGTCCAACAC

TCAATTGGGGGATGGTGGGTCTTTTGCTGCTGGGTGGAACACTGCACCATGCCACAGTCGTCCTACTCTCGCTGGGGGATGGCTTTCCTGCTCTCTGGCTAGAGGTGGTGGTTTTCCTTCGTTTTTGTCAGTCAAGCCGATTCAGCTCACCCTGCAAGAGAGAGAGAGAGAGAGAGAGAGAGAGAGAGAGAGAGAGAGAGAGAGAGAAAAGGCACAGAGACTATGGTCGTCCAACACTCAATTGGGGGATGGT

***Repeat 1 downstream***

*100% identical to Repeat 1 upstream*

*Yellow: TGGG tetramer*

*Green: CTC trimer*

**45S-S: 8x46nt GGG repeat**

***Repeat 1 upstream***

-GGGTCTTTTCTTAAAGGATTAGGGTTAGGCTTGGTAGGGTTTGAGGTTATGATT

TGGGGTTTAGGGTTAGATTTGGGTTTAGGGTAGGGTT

TGGGTTGGGGTTAGGTGTGGGGTTAAATTTGGGTTTAGGGTTGGGGTAGGGTT

TGGGTTGGGGTTAGGTGTGGGGTTAAAGTTGGGGGTAGGGTTTGGAGT

TAGGTTGAGGGGTAAAGTTGGATTTAGGGTTGGGGGTAGGGTT

TGGGTTGGGGTTAGGGGTGGAGTTAAAATTGGGTTTAGGGTTAGGT

TGAGGTTTAGAGTTAGGGGTTTATGGTTGTATGGT

TGGGTTAAGGTTAGGGCACACTTGTCCAACTCTCACTTGGGGGATGGTGGGT

***Repeat 1 downstream***

*100% identical to Repeat 1 upstream*

*Yellow: GGG trimer*

*Green: CTC trimer*
