## Supplemental File SF5 for "Immunoglobulin switch-like recombination regions implicated in the formation of extrachromosomal circular 45S rDNA involved in the maternal-specific translation system of zebrafish"

### Supplemental File SF5: 45S-M Repeat region, longest genomic Nanopore read

>98fea6ad-7872-40b2-9e43-d07918e20948 REVERSE COMPLEMENT (50,267)

CACCCCCTCCTGAAGCGTCCTCTGTCTCGACCGTAACGATTGAAAGAACGAAAACAAGAGTGTACAACTCTTTTAGCGGTGGATCACTCGGCTCGTGCGTCGATGAAGAACGCAGCTAGCTGCGAGAACTAATGTGAATTGCAGGACACACATTGATCATCGACCTGAAGCACAGGCCCTGGTCCATCCCGGGGCCACGCTCTGTCTGAGGGTCGCCTTTGCTATCGATCGGACGGGGAAGAGTCGGTCTTCTCGTGCCGCCCTGCCCCCCTGTCCGCGGCTGGAGCGTCGCAGACCCTGCCCTCCCGCGGTGGCCTACGTCCTCCCAAGTGCAGACCGCCGAACCGTTCGTCCGCCCGCTTGGGGGCGGCTCCATTCCTTCCGCGCGGCTGCCGGCGGTCTAACAGCTGCCCGCGCGCGGTGGACGGGAGCTGCAGCAAACTCCTCGCGTTCCGAGACGACGACAGCGCGATCGGTCGACGCGAGACCGGCGTCCCGCCTCGTGTGGGACGACCGGCTGCCACCACCCGCCTGTTACCCACGACCTCAGCTCAGACGAGAAGACCCGCTGAATTTAAGCATATTACTAAGCGGGAGGAAAAGAAACCAACCGGGATTCCCCCAGTAGCGGCGCGGAGCGAAGAGGGAAAAGTCCAGCGCCGAATCCCCGCTCTGCCGAGGGCGAGGGACCTGTGGCGTACGGAGGGCCGCCTCGGCGCGGGCCGGGGGGCCAAAGTCCTTCTGATGGAGGCTTAGCCCGCGGACGGTGTGAGGCCGGTGTCGGCCCCCGCCCCGCCGGGGTGCGGTTCCTCCCGGAGTCGGGTTGTTTGGGAATGCAGCCCAAAGCGGGCGGGTAAACTCCATCTATACAACACCGGCACGAGACCGATATGACAAGTACCGTGAGGGAAAGTTGAAAAGAACTTTGAAGAGAGAGTTCAACAGGGCGAAACCGTTAAGAGGTAAACGGGCGGGGACCGCACCGTCCGCCCGGTGGATTCAGCCCCGGCGGGGCGGGTCATCTGTCCGTGCGCGCTCCCGCTCCCTCATTCCTGGGGTGGCGCGGGGGTTGGACGCCCGGCGAAGGCCCGGCCGCCGCCGGGTGCATTTCCGCTGCGGTGGAGTTACGCCGCGACCGGCTCCGGTTCGGCTTGGAAGGGTCAGGGGCGAAGGTGGCCCGTCGGTTCAGGCCGTCGGGCTTTACAGCGCCCTCCCGCTTCCCGACCTCGCCGCTTGCTCCGGGGCCGCGGGTGAGTGTCCTCCGCGCCCTCTCTGCCCCTCCCTCCCTGTGGGGAGGTGGGGACGGGGTCCCCGCCCCCGGCGTGGCGCGACAGGTGGACTGTCCTCAGTCCGCCCACGGCTGCGCCGCGCCGCCCAGGCGGGGATCCGACCCACGTTCGGGCGGCTTGAGTGTCCGCGGCGACGCCATCTTCCCACCTGACCCGTCTGCGAAACACGGACCGCGAGTCCAACGCGCGCGCGAAATGAGGGTGGCTCGCGAGCCCTGCGGCGCAATGAAGGTGAGAACGGGTCTCCCCTGCGGGGATCCCCCTGCCCCGGCGGGGCGCACCACCGGCCCGCCTCGCGACCCTCCGGGGGCAGGTGGAGCTGGAGCGCGCGGCGATGGCACCCGAAAGATGGTGAACTATGCCTGGGCAGGGCGAAGCTAGGAAACTCTGGTGGAGGCCCGCCGCGGTCCTGACGTGCAACCGGTCGTCCGACCTGGGCATAGGGCGTAGGTAAGACTAATCGAACCATCTAGTAGCTGGTTCCCTCCGAAGTTTCCTCAGGATAGCTGGCGCTCGCCGATCAAGCAGTTTTATCCGGTAAAGCCAATGACTAGAGGCTCGGGGCCGAAAAACGGCCTCAACCTATTCTCAAACTCTTAAATGGGTAAGAGGCCCGGCTCGCCGGCGCTGGAGCCGGGCGTGGAATGCGACGCGCCTATGGGCCATTTTGCAAGCAGAACTGGTGCTGCGGATGAACCGAACGCCGGGTTATGCCCGATGCCGACGTCATCAGACCCCATAAAGGTGTTGGTTGGATACAGACAGCAGTGACGGTGGCCATGCAAGTTTGGCACCCGCCATGAGTGTAACAACTCACCCGCCGAATCAACTAGCCCTGAAAATGGATGGCGCTGAGCGTTTCGGGCCCATACCTGCCGTCGACGGCACAAGGGACAAAGGAGCGGTCGCGCGCGCCGCAAGCCCGACGAGTAAGGGAGGGCCGCTGGTGGTGGCGCCGAAGCCCAGGCGCGGCCCGGGTGGAGCGCAGTGGGCGGCAGATCTTGGTGGTAGTAGCAAATATTCAAACGAGAGCTTGAAGGCCGAAGGGAGAAGGGTTCCATGTGAACAGCAGTTGAACATGGGTGAGCCGGTCCTAAGGGACGGGCCTACGCCGTTCGGAGGGGAGAGGGCATGGTCCTGTCGCCCTGCTCTCCGACCGAAAGGGCGCCGGTCCAGATCCCGAGCCCGAGCGGTAAGGAGACGGCGCCGCGAGGCGCTCCAGTGCGGTGACGCAAACGAACCCGGAGATGCCAGCCGGCGGCGCCCCGGGGAAAGAGTTCTTTTCTTGTGAAGGGCAGGGCGCCCTGGAACGGGTTCGCCCCGAGAGAGGGGCCCGCGCTCCCTGGAGCGCCGCGCTTCTGGCGGCGTCCGGTGAGCTCGTCGGCCCTGAAAATCCGGGAGAAGGTGTAAATTCGCGCCGGGCCGTACCCATATCCGCAGCAGGTCTCCAGCGTGGAATGCTTCTGGCGTGTTGGAACAAGGCAGAGTAAGGAAGTCGGCAAGTCAGATCCGTAACTTCGGGATAAGGATTGGCTCTAAGGGCTGAGCCGGTCGGGCTGAGGCGAAGCGGGGCCTGGGCCCGGAGCCGCGGGACTGGGGAGCGGCCGCCCTGAGGTGCCCTGGACCCCTGTTCGAGCCGGGCAGCGCGGTGGCGCGCGGTCCCTCCTCTCCCCGCCGCGTCCGCCCGGGGCCTTTGCGGAACCCCGACCCTCCTCCGCCCTTCCCCGTCCGGGGCGGGGTCGGGCGGGGGTCTCGAGGAGGGTGGAGAGGCCGTGGGGGAGGAAAGGGGTGGTTCGTGGCGTCGGGGGCAACGGGTGGTCCTCGCCTGCCGCGCGGATGCCTCCGTTGGCCGGGGCCCGCGGGGGGCGGGATGCGCAGCAGGTTGGCGGCGGCGACCCTGGGGCGCGCGCCGCGCCCTTCCCGCGGGATCTCCGCAGCTACGGCCCCGCGCCGGGGCCCGCGTTCGCGCGTGCGCCCCCTCCGGGGAGTGGCGCGCGGCCCGGCTCCCCCGGCGCGGGCGCTGGCCGGCGGCTACAGCCAGCTTAGAACTGTCGCGGACCAGGGGGAACTGGACTGTTTAATTAAAACAAAAGCATCGCGAAGGCCCTCGGCGGGTGTTGACGCGATGTGATTTCTGCCCAGTGCTCTGAATGTCAAAGTGAAGAAATTCAACGAAGCGCGGGTAAACGGCGGGAGTAACTATGACTCTCTTAATAGTAGCCAAATGCCTCGTCATCTAATTAGTGACGCGCATGAATGGATGAACGAGATTCCCACTGTCCCTACCTGCTATCTAGCGAAACCAATAGCCAAGGGAACGGGCTTGGCAGAATCATGGGGAAAGAAGACCCTGTTGGAGCTTGACTCTAGTCTGGCCCTGTGAAGACATGAGGGGTGTAGAATAAGTGGGAGGCCCTGGCTTCCCGGGCCGGCCGCCGGTGAAATACCACTACTCTTATCGCTCTCTCACTTACCTGGTGAGGCGGGGAAGCCGGGCGTCCCCCGGCGGGGCCGCCCTGCTTCTGGCGTCAAGCGCCTCGGGGCCGTGCGGGGTGGGGCAACCCCCTTCCCCTCCTGCCGCCGACCGGGGCGCGACCCGCCCCGGGGACAGCGTCAGGTGGGGGAGTTTGACTGGGGCGGTACAACCCGTCAAACGGTAACGCAGGTGTCCTAAGGCGAGCTCAGGGGACAGAAACCTCCCGTAGAGCAGAAGGCAAAAGCTCGCTTGATCTTTGATTTTCAGTATGAGCACGGACCGCGAAAGCGGCCTCACGATCCTTCCGGCTTTGGGGTTTTGGGCGGGAGGTGTCAGAAAAGTTACCAGGGATAACTGGCTTGTGGCACAAGCGTTCATAGCGACGTCGCTTTTTGATCCTTCGATGTCGGCTCTTCCTATCATTGTGAAGCAGAATTCACCAAGCGTTGGATTGTTCACCCACTAACAGGGAACGTGAGTTGGCTTAGACCGTCGTGAGACAGGTTAGCTTTACCCTACTGATGTGAAGCCGTTGTTGCAATAGTAATCCCGCTCAGTACGAGAGGAACCGCGGTTCAGACATTTGCTCGTGCGCTTGGTTGAGGAGCCACTGGCGCGAAGTTGTTCATCTGCGGGATTATGACTGAACGCCTCTAAGTCAGAATCCCGCCTAAAAGCAACGATACAGCAGCGCTGCCGGATCTCCGATAGCCCTGGTAACCGGGAAGCCCTCTCCGGGGGCCCCCGGCGGAGAGCCATTCGTGACAGGAACCGGGGCGCGGTTAGAACGAGCGCCGCCCTCTTCCAGTGACGCACCGTACGTTTGTAAAGAAACCCGGTGCTTAAATGACTCGTAGACGACCTGATTCTGGGTCGGGGTGTCGTGCGTGGCAGAGCAGCTCTGTTGCTGCGATCCATTGAAAGTCAGCCCGATCCAAGTTTTGTCGGGCCGGACGCAGGCGACAGGGGCCACCGGTCCGACAGACCCGCGGCGTGCGAAGGCACCAGCTGCGCCGGTGAAGGCTTACATCCCCCGCCACTTGCGCGCAGGCCGAGGCACTGCATGCGGGAGCGTGGCGCGATCCCAATACACCTGGGGCACCAGCGGCAGAGGCCTAGCTCGGCCCGCACCCCACAGCAAAGCCCCTACTGGGGCACCAGCAGCACGCAGCACCTATGGGGTAACCAGCAGAAGGTTCCGTCCCGACCCTAACCCCACAGCAGAGCCTTTATTGGGGCACCAGCAGCACGCAGTACCCACGTGGGTCGCGCTACAGAAGGGGTCCAGCTCGCCCCGCACCCCACAGCAAAGCCTTTAGTGGGGCACCAGCAGCACGCAGTACCTACATGGGGTTACCAGCGGCAGAAGAGGTCCGGACCCGACCCACACCCCACAACAAAGCCTTCGATGGGGTACCAGCAGCACGCAGTACATAAATGGGTCACCATGGCGCGAGTCTGGCGCGCGCGCCTTACCGAAGTCAGTGTCCCTGCTTAAGAGTGGGTGCCCGGGCACGGCTGGTGAACGTTCCATGTGGTGGGGTCAGAGGCCGGCCGGTGTCCCCCGGCCTAGCCAGCCAGCCAGTCAGTCAGCCCGGCCCAGCCCAGCCCAGCCCAGCCAGCCAGCCAGCCAGCCAGCCAGCCCAGCCCAGCGCTGTCCAGCCAGCCCTGTCTCAGCTCCAGCCCAGCAAGCCCAGGCCGGCCGGCCAGCCAGCCAGCCACAGCAAGCCCAGGGGCCAGGCCAGGGCCCAGCCAGCTCCAGCCAGCCCTGCCCTGCCCTGCCCTGCCCTGCCCAGTCTGTGCCCAGGCCGGCCAGCCAGCCAGCCAGCCAGCCAGCCAGCCAGCCCAGGGCCAGGCCAGGCCAGGCCAGGCCAGGCCAGGCCAGGCCAGGCCAGCCCAGCTCGGCCAGGCCAGGCCCAGCCCAGCCAGCCCAGTCAGCCAGCCAGCCCAGCCTGTTGCAGCCAGCCAGCCAGCCAGCCAGCCCAGCCCAGCCCAGCCCAGGCAAGCCCAGGCCGGCCAGCCAGCCAGCCAGCCAAGGCCAGGCCAGCTCCAGCCCAGCCAGCCAGCCAGCCAGCCCAGGCACAGGCCAGGCCAGGCCAAGGCCAGGCCAGGCCAGGCCAGGCCAGGCCAGCTCCAGCCTGGGCCAGGCCAGCCCAGCCGAGCCCAGCCAGCCAGCCAGCCAGCCAGCTCAGCTCCAGCCCAGCCCAGCCCTGCCCTGCCCTGCCCTGCCTCTTGCCCCGCCCAGCCCAGCCTAGCCCAGCCTGTGCCAAGCCCAGGCTGGCCAGCCAGCCAGCCAGCCAGCCAGCCCAGGCCAGGCCAGGCCAGGCCAGGCCAGGCCAGGCCAGGCCAGGCCCAGCCCAGCCAGCCAGCCAGCCAGCCAGCCAGCCCAGGCCAGGCCAGGCCAGGCCATGCCCAGCCCAGTTGTGGTTAGCCAGCTCCAGGCTGGGCAAGCCCTGGCTGGGGCAGCAAGCCCAGCTCAGCCAGCCAGCCCAGGCCAGGCAAGCCCTGCCGGCCAGCCAGCCAAGCCCAGCCAGCCAATCAGCCGCCCGGCTGGGCGGCCAGCCTGGCCAGGTCAATCAGCCCAGCCCAGCCCAGCCTAGCCTAGCCTAGCTCCAGCCCAGCTCCAGCCCAGCTCCAGCCCAGCTCCTGCCCTGCCCAACCTGGACAGCCAAGTTCTGCCTCAGCCAGCCAGCCAGCCATCCTGGTGTTGAGAGGCTTGGGCTTTAGAGTGGTTGGTCGATAGTCCATGGCCGAGGGAGAGTGAGCACTATCTCCTCCCACTTCTCCTCTTTCCTTACCACCACCACCACCACCACCACCACCACCACCACCACCACCACCACCCCGGAGGCCCGGATTGGGCTTGGGGATAGGGGTAAGGGTTGCCTGGAGGCTAGTGGAGTTAGGCTGCCCGGGAGGCCCGGGGATTGGGCTTGGGGATAGGTGTTGTCAGTCGCTGGAGGCCGGGGGGGTGGCTAGGGGCTGCCCTTGGAGGCCCGGGGATTGGGCTTGGGGATAGGGTGGGCGGGGCTGCCTGGAGGCCTGGGTGGGTTAGGCTGCCAGGAGGCCCGGGGATTGGGCTTGGGCATAGGGGGTAAGGGGTTGCCTGGAGGCTTGGGGGTGGGTTAGGCTGCCTGTGGAGGCCCGGGGATTGGGCTTGGGGATAGGGGTAAGGGTTGCCTGGCGCTTGGGGGTGGGTTAGGCCGCCAGGAGGCCCGGGGACTGGGCCTGGGCATGGTAAGGGTTGCCTGAGGCTTGGGGTGGTTAGGCTGCCTGAGGCCCGGATCACTTGGGCATAGGGTAAGGGTTGCCTGGGAGGCTTGGGGTGGGTTAGGCTGCTGGAGGCCCGGGGGATTGGCTTGTGTGATAGGGTAAGGGTTGCCTGGAGGCTTGGGGTGGGTTAGGCCGTGCCAGGAGGCTCCGGGGATTGGGCTTGGGCATAGGGTAAGGGTTGCCTGGAGGCTTGGGGGTGTAAGGTTAGGGCTGCCCGGGAGGCCCGGGGATTGGGCTTGTGATGGGGTATGGGGTTGCCCGAGGCTTGGGGCGGGGTTGCAAAGCTGCCAGATGCCCGGGATTGGGCTTGGGCATAGGGTAAGGGTTGCCTGGAGGCTTGGGGTGGTTAGCTGCCTGAGGCCTGGGGATTGGGCTTGGGGATAGGGTGGGCGGCTGCCTGAGGCTTGGGTGGGTTAGGCTGCCTGGAGGCCCGGGGATTGGGCTTGGGGATAGGGTAAGGTTGCCTGGAGGCTTGGGGGTGGGTTAGGGCTGCCAGGAGGCCCGGGGATTGGCTTGGGCATAGGTAAGGGTTGCCTGAGGCTTTGGGGGTGGGTTAGGCTGCCTGGAGGCTCCGGGGATTGGGCATTTGGGGATAGGGTAAGGGTTGCCTGAGGCTTGGGGGTGGGTTAGGGCTGCCAGGAGGCCCGGGGATTGGCTTGGCAATAGGTGAGGGTTGCCTGGAGGCTTGGGGTGGGTTAGTTGCCAGGAGGCCCGGGGATTGACTTGGGGATAGGGTAAGGGTTGCCTGAGGCTTGGGGGTGGGTTAGGGTTGCCAGGAGGCCCGGGGATTGGGCTGAGCATAGGGTAAGGGTTGCCTGAGGCTTGGGGGGTTAGGGTTAGGGTTGCCCGGGAGGCCTGGGGATTGAGCTAGGGATAGGTAAGGGTTGCCTGAGGCTTGGGTGTTAGGGCTAGGGTTTCCCGGAGGACTGGAATTGGTTAGGGTTGCCTGAGACTTGGGGGTGGGGTTAGGGCTGCCCTGGAGGCCTGGGGATTTGGGGCTTGGGGATAGGTTTAGGGGGTTACCCGGAGGCTTAGTTGTTGGGCTTGTGGAGCTTTGGCTTGCCTGGTTGCCTGAGGCTTGGGATGCCACCCCGCGCCAGCCAGGAAAGGTCAGCCAGACCAGCAGCAACCGCCCGGCTGCCGAAGGCAAAGCCGTCGCGGTCACCATCGGCACCGATTCGGCAAGGGAGAAGCAAGAAGCGGCCGCCAACCCCGACTATCCATGGTGGCCCCGAGCTCGTGGGTTCACAGCCGGGCGGGACGTGACCTCTTGGGTCTCTCTCTCCCGGAGGTCTGGGCAGCTGTCACTGGTTATCGCTGTGACCTGCTGTCGAGGCTCTCGGGGAAGGCGTGATTTAGTGGGACTTAGGTTTTGGAGCGCGTAACTACGTGCCACCGCCTTCCACCACCTGACCGATGGGAATTGTGCACGTGATGCCCCAGGGGCAGGTCTCTGCCCGGTCCTCCGACAAAGCGGGACCGCTGTGGTGGGGTGGGTTCCTCCCCGTTTTGGGTATTTCGAAAGGAGGGCGAGGCGGGCTCACCAGACCTCGACCCTTGCTCCTTTGCCAAAGACTCAGATGGCCCGTCGATCTCAGACTGCCCTGTACCGACGGAACAAGTGCGCCAGTCGGTGCTGTGCGTGTGTTTACCAGGGCAGATCCTCTCTCTGCCCCCTCCTTTTCCCCACCACCCGGGTGTAATCCCGGGCGCGTAGGCTGACTTGGGGGTGGGGAAGCTGTGTGGTAGGCGCGCGCGGCATCTGCGGCGCACTCGATCCCGTTGGGTGAAGCACGATCCTTCCTGGTGATGGCCCGACCCAGCGGCCAGCCCACGCGCCGGCAGACCCCCTCATTTCCATGTGTGCGGTGGGTGGTAGGCGGCGGTACCTGGGAGCCCAATCCCCTCCCACTTAGCCCAGCGCCCTCCTGCGAGCAGTCGTGACGCCTCGGCGGAGCGAGGCCCGCTTGTAGGTAGGCGGCGCGGGAGCCGTGGTGGGCGGAGGGCCCTCTTCCGCCGAGTCCTCCTTTCCCCTCTCTCCTCCGGCACTCTCCGGGCCTTACCACCACGCGATTCACCCGAGGCGAGGGCCACCTGGTTGATCCTGCCAGTAATATATGCTTGTCTCAAAGATTACTTCATGCAAGTCTAAGTGCACACGGCCGGTACAGTGAAACTGCGAATGGCTCATTAAATCTGCTGCGGTTCCTTTGATCGCTCCACCCGGTTACTTGGATAACTGTGGCAATTCCAGAGCCACATGCCAACGAGCGCCGACCTGCCGCCCTTCCTCCCTCGGGGCGGGGGTGGGTACCCGGGGACGCGCATTTGGTCAGATCCAAACCCATGCGGCGCGGGCGGTGGAGAGGGGGCCTCGCGCCTGGCCTGCCGCCGTCCGCCCCCGGCCTCGCTTTGCGACTCTAGATAACCCGGGCCGATCGCGCGCCCTCGCGGCGGCGACGGTTCATTCGAATGTCTGCCCTATCAACTTTCGATGGTAGGTCCGTCGCCTACCATGGTGACCACGGGTGACGGGGAATCAGGTTCGATTCCGAGAGGGAGCCCGAGAAACGGCTACCACATCCAAGGAAGGCAGCAGGCGCGCAAATTACCCATTTCCGACACGGAGAGGTAGTGACGAAAAATAATGCAGGTCTCTTTCGAGGCCCTGCAATTGGAATGAGTGCATCCCAAACCCATGGGCGAGGACCCATTGGAGGGCAAGTCTGGTGCCAGCAGCTGCGGTACTCCAGCTCCAATAGCGTATGCTAACGTTGTTGCAGTTAAAAAGCCTGTAGTTGATCCGGGGACCGGGCCGCGCGGTCCGGCCGCGAGGCGAGCCACCGCCGTCCTGGACCCCCAGGCCTCCCGGCGCCCTTCGGATGCCCTGACTGGGTGTCCTCGGTCAGGCCCGAGCGTTTACTTTTGAAAAATTAGAGTGTTCAAGGTAGGCCGGCACCGCGCCCCATTGAATACCCCAGCTAGGAATAATGGAATAGGACCCCGGTTCTATTTTCTGTGGGTTTCCGGAACCCGGGGCCATGATCGAGAGGACGGCCGGGGCATTCGTATTGCGCCGCTAGAGGTGAAATTCTTGGACCGGCGCAAGACGGACCGGAGCGAAAGCGTTTGCCAAGAACGTTTTCATTAATCAAGAACGAAAGTCGGAGGTTTGAAGACGATCAGATACCGTCGTAGTTCCGACCGTAAACGATGCCGACCCGCGAGATCCGGCGGCGTTTATTCCCATGACCCGCCGGGCAGCGTTGCGGGAAACCACGAGTCTCTGGGCTCCGGGTAGCATGGCTGCAAAGCTGAAACTTAAAGGAATTGACGGAAGGGCGACCACCAGGAGTGGAGCCTGCGGCTTAATTTGACTCAACACGGGGAACCTCACCCGGCTCCGACACGGAAAGGATTGACAGATTGAGGTTCTTCTCGATTCTGTGGTGGTGGTGCATGGCCGTTCGTAGTTGGTGAGCGATTTGTCTGGTTGATTCCGATAACGAACGAGACTCTGGCATGCTAACTGCCATTACGCGCCCGCGCGGTCGCAGGCGTCTGCAACTTAGAGGACAAGTGGCGTTCAGCCACGCGGAGACTGAGCAATAACAGGTCTGTGATGCCCTTAGATGTCCGGGCTGCACGCGCCACAATGGGCGGATCAACGTGCCTACCCTGCGCCGACAGCGGGTAACCCGTTGAACCCCGCCCGTGATGGGGACCGGGATTGAAACTATTTCCCGAGAACGAGGAATTCCCAGTAAGCGCAGGTCATCAGCTTGCGTTGATTAAGTCCCTGCCCTGTACACACCGCCCGTCGCTACTACCGATTGAGCGGCTCAGTGAGGTCCTCGGATCGGCCCCGCCCGGGGCTCCTACCGGGGCCCTGTGGGAGCGCCGAGAAGACGATCGAACTCGGTCGTTTAGAGGAAGTAAAAGTCGTATCAAAGGTCGCAGGTGAACCTGGGTGAAGGATCATTAATGGGCCGAGGGGATCTCTCCTCACGCCCCAGAGTGGCGAAGCCGCCAGGTAAGCTCCCGGCGCGGTGCGGGAAGCCCACGGGTCTATTCTCCCCACCATCGCGCGCGCGAGCGTGAATGGTGATCCAAAGGCGCTGGCGGCTGTGGCTCCCGGCGGGTACCTGGTCTCACCACCTCGACCTGTCCCCTCTGCGGGGGGGAGAGCGACGGAGGTGCGGGGGGCTGTGGGTTTGCAAGCACTCTTCGCGTTTCGCCACCCGGGGGAACAGGAGACGCCTGTCCCGGGGCCCTGCCCGCCATGTTTAATTTCCCCACCCCCTCCCGAAGCGTCCTCTGTCTCGGACCGTAACGATTGAAAGAACGAAACAAGAGTGTACAACTCTTAAAGCGGTGATCACTCGGCTCGTGCGTCGATGAAGAACGCAGCTATTGCGAGAACTAAATGTGAATTGCAGGACACACATTGATCATCGACCTTCGAACGCACATTGCGGCCCCGGGTCCATCCCGGGGCCACGCCTGTCTGAGGGTCGCTTGCTATCGATCGGACGGGGAAGAGTCGGCCTCGTGCCGCCCTGCCCCCGTCCGCGGCTGGAGCGATGACCCTGCCCTCCCGCGGTGGCCTACGTCCTCCAAGTGCAGACCGCCGAACTGTTCGTCCGCCCGCTTGGGGGCGGCTCCTACTTTCCCCGCGCGGCTGCTGGCGGTCTAACAGCTGCTCCGCGCGGTGGACGGGAGCTGGGCAGTAAACTCCCTCGCGTTCCGAGACGACGACAGCGCGATCGGTCGACGCGACATCGGCGTCCCGCTCCGTGTGGGACGACCGGCCGCCACCACCCGCCCGTTGGCCCACGACCAGCTCAGACGAGAAGACCTGCTGAATTTAAGCATATTACTAAGCGGAGGAAAAGAAACCAACCGGGATTCCCCCAGAAGCGGCGAGCGAAGAGGAAGTCCATGCCGAATCCCTGCCTTTCTGCCGAGGGCGAGGGACCTGTGGCGTACGGAGGGCCGCCCTCTCGGCGCGGGCCGGGGGCCAAAGTCCTGATGGAGGTGCCAAGCTTAGCCCGCGGACGGTGTGAGGCCGGTGTCGGCCCCGCCCTGCCGGGGTGCGGTTCCCGGAGTCGGGGTTGTTTGGGAATGCAGCCCAGCGGGTGGTAAACTCCATCTATAGCTAAATACCGGCACGAGACCGATAGCGGACAAGTACCGTGAGGGAAAGTTGGAAAGAACTTTGAAGAGAGAGTTCAACAGGGCGTGAAACCGTTAAGAGGTAAACGGTGGGGACCGCACCGTCCGCCCGGTGGGATTCAGCCCGGCGGGGCGGGGTCGGCCCGTCCGGCGCGCTCCCTTCGCCTCATTCCTGGGGGTGGCGCGGGGGCTGACGCCCGCGATGGCCTGGCCGCCGCCGGGTGCATTTCCGCCGCGGCGGAGCGCCGCGACCAGCCCTGGTTCGGCTTAGCGGGTCAGGGGCGAAGGTGGCCCGTCGGTTCAGCCGTCGGGCTTTACAGCGCCCTCTCCCGCCCCGACTTCTTCGCTTGCCTCCGGGGCCGCGGGTGCGTCCTCCGCGCTCCTCTCTGCCCTCCTCCTCCCTGTGGGGAGGCGGACGGGGTCCCCGCCCCCGGCGTGCGCGACAGGGGTGGACTGTCCTCGTCCGCCCACGGCCGCGCCGCGCCGCCCAGGGCGGATCCGACCCACGTTCGGGCGCGCCCGAGGCCCGCGGCGACGCCGGCCTCCCACCTGACCCGTCTTGAAACACGGACCAGAGTCCAACGCGCGCGCGAGTCAGGAGTGGGTGGCTCGCGAGTTTTCCTGCGGCAATGAAGGTGAGAACGGGGTCTCCCCCGCGGATCCCCCCGCCCCCGGTGCGGGGCGCACCACCGGCCCGCTCCCGCGACCAAAGGTGGAGTTGAGCGCGCGATGGCACCCGGAAAAGATGGGTGAACTATGCCCGGGCAGGGGCGAAGCCAGGGGAATCCTGGTGAGGCCCGCCAGTGGTCCTGACGTGCAGAACTGGATGTCCGACCTGGGCATAGGGGCGAAAGACCAATCGAACCATCTAGTAGCTGGTTCCTCCAGTTTCCTCAGATGCTGGCGCTCTTCGCCGATCAAGCAGTTTTATCCGTAAAGCTAATGACTAGAGGCCTTGGGGCCGAAACGGCCTCACAACTTATTCTCAAACCCTTAAATGGTAAGAGTCCGCTCGCTGGCGCTGGAGCCGGGCGTGGAATGCGACGCGCCCAGTGGGCCATTTTTGGTAAGCAGAACTGGTGCTGCGGGATGAACCGAACGCCGGGTTAAGGCGCCCGATGCCGACGCTCATCAGGACCCCATAAAGGTGTTGGTTGATATAGACAGCAGGACGGTGGCCATGTGGAAGTCGGCACCCGCCATGATGTAACAACTCACCTGCCGAATCAACTAGCCCTGAAAATGGATGGCGCTGGAGCGTCGTAGCCCATACCCGGCCGTCGACGGCACAAGGGACACGCGCGAGCGGTCGCGCGCGCTGCAAAGCCTCGACGAGTAGGAGGGCCGCCGCGGCGGCGCTGGAAGCCTGCGGCGCGGGCCCGGTGGAGCGGCCGCGGGCGCAGATCTTGGTGGTAGTAGCAAATATTCAAACGAGAGCTTTGAAGGCCGAAGTGGAGAAGGGTTCCATCGTGAACAGCAGTTGAACATGGGCGAGCCGGTCCTATGAACGGGCCTACGCCGTTCGGAGGGAGGGGCGATGGCTCCCGTCGCCCCCGCTCGACCGAAAGGAGTCGGGTCCAGATCCCCGAGCCCGAGCGGCGGAGACGGGCGCCGCGAGGCGCCCAGTGCGGTGACGCAAACGAACCCGAGATGCCGGCGGGTGCCCCGGGAAAGAGTTCTCTTTCTTGTGAAGGGCAGGGCGCCCTGGAACGGGTTCGCCCGAGAGAGGGGCCCGCGCCCTGGAAAGCGCCGCGCTTCTGGCGGCGTCCGGGTGAGCTCTCGTCGGCCCTTGAAACTGGGAGAAGGTGAATCTTCGCGCCGGGTCGTACCCATATCCGCAGCAGGTCTCCAAGGTGAACAGTTCTGGCGTGTTGGAACAAGGCAGAGTAAGGGAAGTCGGCAAGTCAGATCCGTAACTTCGATAAGGATTGGCTCTAAGGGCTGAGCCGGTCGGGCTGAGGTGCGAAGCGGGCCTGGCCCGAGCTGCGACTGGGGAGCGGCCGCCCCGAGGTGCCCTGACCCCCGCTCCCGAGCCGCGGGGTGCGCGGTGGCGCACCTCTCTCCCCCGCCGCGTCCGCCTGGCCTTTGCGGTCTCGACTCCCGACTTTTCCCCCTCCGCCCTTCCCGTCCGTGGGGTCGGGCGGGGGGTCTCGGAGGAGGGTGGAGAGGCCGTGGAGGAAAGGGGTGGTCTGTGGCGTCGGGGGCAACGGGAGGGTCCTCGTTCGCCGCGCGATGCCTCCGCTGGCCGGGCCCGCGGGGGCGGGATGCGCAGCGGTTGGCGGCGGCGACCCTGGCGCGCCGCGCCCTTCCCGCGGATCTCCGCGTTGGGCCGGCCCCGCGCCGGGCCCGCGTCCGCGCGCCCCTCCGGGGGAGGGCGTTAGCCTCGGCTCCCCCGGTGCGGGCGCTCGGCCGGCGGCTAGCAGCCAGTTTAGAACTGTCGCGGACCAGGGAATCCGACTGTTTAATTAGTGCAACAGCATCGCGAAGGCCCTCGGCGGTGTTGACGCGATGCATTTCTGCCCAGTGCTCTGGAATGTCAAAGGAAGAAATTCAACGAAGCGCGGGGTAAACGGCGGGAGTAACTATGACTCTCTTAAGGTAGCGCCAAATGCCCTCGTCATCTAATTAGTGACGCGCATGAATGGATGAACGATTCCCACTGTCCCGACTCGCTATCTAGCGAAACCACAAAGCCAAGGGAACGGGCTTGGCAGAATCAGCGGGGAAAGAAGACCCTGTTGAGCTTGACTCTAGTCTGCCCTGTGAAGAGACATGAGGGGTGTAGAATAAGTGGGAGGCCCCCGGGCTTCCCGGGCCGGCCGTCGGTGAAATACCACTATTTCCATCGTTTCCTCACTTACCCGGTGAGGCGGAGCTGCCGCTCCCGGGGGCCGCCCCTGCTTCGGCGTCAAGCGCCCCGGGGCCGTGCGGGGGTGGGGGCAACCCCTTCCCTCCCGGCCGCCGACCGGGCGCGACCCGCCCCGGGACAGCGTCAGGTGGGAGTTTGATCAGGCGGTACACCTGTCAAACGGTAACGCAGGTGTCCTAAGGCGAGCTCAGGGGACAGAAACCTCCCGTGCAGAGCAGAAGGGCAAAAGCTCGCTTGATCTTGATTTTCAGTATGAGTACGGACCGCGAGCGGGGCCTCACGACTTTTCCGGCTTTTGGGGTTTTAAGCGGGAGGTGTCGAAAGTTACCATATACAACTGGCTTGTGGCGGCCAAGCGTTCATAAGCGACGTCGCTTGATCCTTCGATGTCGGCTCTTCCTATCATTGTGAAGCAGAATTCACCAAGTGTTGATTGTTCACCCACTAACAGGGAACGTGAGCTGGCTTAGACCGTCGTGAGACAGGTTAGTTTTACTCCTACTGATGTGAGCCGTTGTTGCAATAGTAATCCTGCTCAGCACGAGAGAGGAACCGCGGGTTCAGACATTTGTCTGTGCGCTTGGTTGGAGGAGCCACTGGCGCGAAGCCACCATCTGCGGGATTATGACTGAACGCCTCTAAGTCAGAATCCCGCCTAAAAGCAACGATACAGCAGAAGCGCCGTCGGATCTCCGGATAGCCCTGCAACCGGGAAGCCCTCTCCGGGGGCCCCGGCGCGGAGAGCTATCGTGACAGGAACCGGGGGCGCGTGGTTGTGAACGAGCGCCGCCCTTCTTCCAGTGACGCACCGCATGTTTGTGGGGAACCCGGTGCTTAAATGACTCGTAGACGACCTGATTCTGGGGTCGGGTGTCGCGTGGCAGAGCAGCTCTGTTGCTGCGATCCATTGAAAGTCAGCCCTCGATCCAAGTTTTGTCGGGCCGGACGCAGGCGACAGGGGCCACTGGTCCGACAGACCCGCGGCGTGCGAAGGGGCACCAGCTGTGTCGGTGGTTGAATGTGCCATCCCCCTCGCCACTTGCGCGCAGGCCGAGGCACCAGCGGCGGAGCGTGGCGCGATCCCACACCTGGGGCACCAGCGGCAGAGGGAGGCCTAGCCTCGGCCCGCAACCTCACAGGCAAAGCCCCTACTGGGGCACCAGCAGCACGCAGTACCTATGTAAGTAATCCAGCGGCAGAAGGTCTCTGTCCCGACCCTAACCCCACAGCAGAGCCTTTATTGGGGCACCAGCAGCACGCAGTACCCACGTGGGTCGCCAGCGGCAGAAGGGGGTCCAGCCTCGCCCCGCACCCCACAGCAAAGCCTTTGCGGGGCACCAGCAGCACGCAGTACCTACATGGGGTTACTAGCGGCAGAAGAGGTCCGGACCCGACCCACACCCCACAACAAAGCCTTCGATGGGCAACAGCAGCACGCAGTACATAAATGGGTCACCAGCGCAAGGAGTCTGGCGCGCGCCTACCGGGAAGTCAGCGTCCCCGGCTTAAGAGGTGCCAAGTACGGTTGGTGAACGGGGTTCCATGGTGGTCAGAGGCCGGCCGGTGTCCCCGGCCTAGCGCCGCCAGTCAGTCAGCCCGGCCCAGCCCAGCCCAGCCCAGCCAGCCAGCCAGCCAGCCAGCCAGCTCCAGCCCAGCGCTGTCCAGCCAGCCCTGTCCAGCCCAGCCCAGGCAAGTTCAGGCCGGCCGGCCAGCCAGCCAGCCAGCGCCCAGGCCAGGCCAGCCCAGCCAGCCCAGCCAGCCTCCTGCCCTGCCCTGCCCTGCCCTGCCCAGCCCAGCCCAGGCCGGCCAGCCAGCCAGCCAGCCAGCCAGCCAGCCAGCCCAGGCCAGGCCAGGCCAGGCCAGGCCAGGCCAGGCCAGGCCAGGCTAGGCCCAGCCTGGCCAGGCCAGGCTCAGCCCAGCCAGCCCAGTCAGCCAGCCAGCCCAGCCCAGCCAGCCAGCCAGCCAGCCAGCTGAGCTCCAGCCCAGCTCCAGCCCAGGCAAGCCCAGGCCGGCCAGCCAGCCAGCCAGCCAGCCAAGGCCAGGCCAGCCCAGCCCAGCCAGCCAGCCAGCCAGCCCAGGCAAGGCCAGGCCAGGCCAGGCCAGGCCAGGCCAGGCCAGGCCAGGCCAGGCCAGGCTCCAGCCCAGCCAGGCCAGCCCAGCCGAGCCCAGCCAGCCAGCCAGCCAGCCAGCCCAGCCCAGCCCAGCCCAGCCCTGCCCTGCCCTGCCCTGCCCTGCCCTGCACCAGCCTGTGCCCAGCCCAGCCCAGCCAAGCCCAGGCCGGCCAGCCAGCCAGCCAGCCAGCCAGCCCAGGCCAGGCTAGGCCAGGCCAGGCCAGGCCAGGCCAGGCCAGGCCCAGCCCAGCCAGCCAGCCAGCCAGCCAGCCGCCCAGGCCAGGCCAGGCCAGGCTGCCCAGCCCAGCCAGCTGTGCCAGCCCAGGCCAAGCAAGCCCTGGCTGGGCAGCAAGCCCAGCTCAGCCAGCCAGCCCAGGCCAGGTGGCAAGCCCTGGCCGGCCAGGCCAGCCAAGCCCAGCCAGCCAATCAGTCAGTCAAGCCAAGGCCAGTCAGTCAGGTCAATCAGCCCAGCCCAGCCCAGCCTAGCCTAGCCTAGCCTAGCCCAGCCCAGCCCAGCCCAGCCTTGGCCTCTGCCCCCTGTCTCTGCCTGGACCAGGCCTGCTCAGCCAGCCAGCCAGCCATCCTGGTGTTGGAGAGGCTTTCTTTCTGGGCTTTAGAGTGGTTGGTCGATAGTCCATGGCTGGAGGGAGAGTGAGCACCATCTCCCTCCCACTTCCCTCCTCTTCCTTTACCACTCACCACTCACACCACCACCACCACCACCACCACCACCACCACCCCGGAGGCCCGGGATTGGGCTTGGGATAGGGTAAGGTTGCCTGGAGGCTTGGGGGTGGGTTAGGGCTGCCCGGGAGGCCTGGGGATTGGGCTTGGGATAGGGGTGCGGCTACCTGGAGGCTTGGGGGTGGTTAGGGCTGCCCGGGAGGCCCGGGGATTGGGCTGGGGATAGGGTAAGGGCTGCCTGGAGGCTTGGGTGGTTAGGGCTGCCAGAGGCCCGGGGATTGGGCCGGGGCATAGGGTGGGCGGCTGCCTGGAGGCTTGGGGTGGTTAGGCTGCCTGAGAGGCCCGGGGATTGGGCTTGTGGGATAGGGGTAAGGGTTGCCTGGAGGCTTGGGGGTGGTTAGGGGTTGCCAGGAGGCCCGGGGATTGGGCTTGGGCATAGGGTAAGGGTTGCCTGAGGCTTGGGGGTGGGTTAGGGCTGCCTGGAGGCCCGGGGGATTGGGCTTGGGCATAGGGTAAGGGTTGCCTGAGGCTTGGGGGGTGGGTTAGGCTGCCTGGAGGCCTGGGATTGGGCTTGGGGATAGGGTAAGGTTGCCTGGGAGGCTTGGGGGGTGGGTTAGGGGCTGCCAGGAGTGCCCGGGGATTGGGCTTGGGTATAGGGGTAAGGGTTGCCTGGAGGCTTGGGGGTGGGTTAGGCTGCCTGGAGGCCCGGGGATTGGGCTTGGGGATAGGGTATGGGTTGCCTGGAGGCTTGGGGTAGGGGGTTAGGGCTGCCAGGAGGCTCCGGGGATTGGGCTTGGGCATAGGGTAAGGGTTGCCTGGAGGCTTGGGGTGGGTTAGCTGCCTGGGAGGCCCGGGGATTGGGCTTGGGATAGGTAAGGGTTGCCTGGAGGCTTGGGGGTGGGGTTAGCTGCCTGGAGGCCCGGGATTGGGCTTAGATAGGGTATGGGGCTGCCTGAGGCTTGGGGTGGGTTAGGGGCCAGGAGGCCCGGGGATTGGGCTTGGCATAGGGGTAAGGTTGCCTGGAGGCTTGGGGTGGGGTTAGGCTGCCTGGAGGCCTGGGATTGGGCTTGTGGGATAGGTAAGGTTGCCTGGAGGCTTGGGGTGGGTTAGGCTGCCAGAGGCCCGGGGATTGGGCTTGGGCATAGGGTATGGGTTGCCTGAGGCTTGGGGTGGGTTAGGGTTGCCAGGAGGCCCGGGGATTGGGTTTGGGGATAGGGTAAGGGTTGCCTGCGTTTGGGTGGGTTAGCTGCCAGGGAGGCCCGGGGATTGGGCTTGAGCATAGGTAAGGGTTGCCTGGAGGCTTGGGGGGGTTAGGGTTAGGGTTGCCCGGAGGCCTGGGATTGAGCTTGGGGATAGGGGTAAGGGTTGCCTGAGGTTTGGGTGTTAGGGTTAGGGTTTCCTGGAGGACTGGGAATTGGGTTAGGGGTTGCCTGGAGATCAGTGGGGTTAGGGCCTGGAGGCCTGGGATTGGGCTTGGGGATAGGGTTAGTTGCCCGAGGTTTAGTTGTTGGGCTTCAGGAGCTTTGGCTTGCCTGGTTGCCTGAGGCTTGGGAGAAGTGCCACCCGCGTGCCAGCCAGAAAGGTCAGCTGTGGGTGACCAGCACCGCTCCGGCTGCCGAAGGCAAAGTCAGTCGCGGGTCACCATCGGCACCCGATTCGGCAAGGGAGAAGCAAGAAGCGGCCGCTAACCCTGACTATCCATGGCCCCGAGCTCAGGTTCACAGCTATGGACGTGACCTTCTTGGGTCTCTCTCTCCCGGAGGTCTGGCAGCCGTCACTGGTTATCGCTGACCTGCCGTCGAGGTTTCGGGGAAGGCGTGATTTAGTGGACTTAGGTTTTGAGCGCGTAACTACGTGCCACCGCCTTCCACCACCTGACCGATGGGAATTGTGCGCGATGTCCCAGGGCAGGCCCCTTGCCCGGTCCTCCGACAAAGGGGACCGCTGTGGTGGGGTGGGTTCCTCCCCGTTTTGGGCATTCGAGTGAGGCGAGGCGGGCTCACCCAGACTCCTCGACCTTGCTCCTTTGGGTTAAAGACTCAGATGGCCCGTCGATCTCAGACCGCCCTGTACCGACGGCGCGCCAGTCGGTGCTGTGCAGTGCGCTTCACCAAGCAGCATCCTCTGCCCCCCTCCTTCCTCACCACCCGGGTGTAATCCCGGGCGCGTAGGCTGACTGGGGTGGGGAAGCTGTGTGGTAGGCGCGCGGGCATCTGCGGCGTACCGATCCTGCTGCGAAGCACGATCCTTCCTGGTGATGGCCTGACCCAGCGGCCAGCCCAACGCCGGCAGATCCCCCTTACTTATGTGTGCGCGGGGGTGGCAGGCGGCGGTACCTGGAGCCCAATCCCCCTCCCACTTAGAAGTCTGCGTGCCCTCCCAGAGCAGTCGACGCCTCGGCGGAGCGAGGCCCGCCGGTAGGTGCAGGCGCGGAGCTGTGGTGCGGGCGGAGGGTTCTCTTCCGCCGAGTCCTTTCCCTCTCTCCTCCGGCAATCTCCGGCCTACCACCACGCGATTCACTCCTGGAGGCGAGGGCTACCTGGTTGATCCTGCCAGTAATATATGCTTGTCTCAAAGATTAAGCCATGCAAGTCTAAGTGCACACGGCCGGTACAGTGAAACTGCGAATGGCTCATTAAATCAGTTATGGTTCCTTTGGATCGCTCCACCCGGTTACTTGATAAACTGTGGCAATTCCAGAGCTAATACATGCCAACGAGCGCCGACCTGCCGCCCTTCTCCCCTCGGGGCGGGGGGTGGGTAAATTCGGGGACGTGCGTGTATTTATCAGATCCAAACCATGCGGGGTGCGCGGCGGTGGAGAGGGGGCTCGCGCCTACCCGCTCGCCGTCCGCCCCCGGCCTCGCTTTGGTGACTCTAGATAACCCGGGCCGATCGCGCGCCCTCGCGGCGGCGACGGTTCATTCGAATTGCCCTATCAACTTTCGATGGTAGGTCCGTCGCCTACCATGGTGACCACGTGACGGGGGAATCAGGGTTCGATTCCGGAGAGGGAGCCTGAGAAACGGCTACCACATCCAAGAGGAAGGCAGCAGGCGCGCAAATTACCCATTTCCGACACGGAGAGGTGGTGACGAAATAACAATGCAGGTCTCTTTCGAGGCCCTGCAATTGAAATGAGTGCATCCCAAACCCATGGGCGAGGACCCATTGAGGGCAAGTCTGGTGCCAGCAGTCGCGGTAATTCCAGCTCCAATAGCGTATGCTAACGTTGCCGCAGTTAAAAAGCTCGTAGTTGATCTCGGGACCGGGCCGCGCGGTCCGCCGCGAGGCGAGCCACCGCCGGTCCTGACCCCAGGCCTTCCCGGCGCCCCCGGATGCCCTTCTTGACTGGGTGTCTCGGTTTGGGCCCGGAGCGTTTACTTTGAAAAATTAGAGAAACTAAGGCAGGGCCGGGCACCGCGCCCCATTGAATAATTCTGCTAGGAATAATGGAATAGGGACCCTGGTTCTCTATTTCTGTGGGTTTCCGGAACCCGGGGCCATGATCGAGAGGGACGGCCGGGGGCATTCGTATTGCGCCGCTAGAGGCGAGTTCTTGGACCGGCGCAAGACGGACCGAGCGAAAGCGTTTAAGCACAAGAACGTTTTCATTAATCAAGAACGAAAGTCGGAGGTCTGAAGACGATCAGATACCGTCGTAGCTCCGACCGTAAACGATGCCGACCCAGCGATCCGGCGGCGTTTATTCCCATGACCGCCGGGCAGCGTTGCGGGAAACCACGAGTTCCTGGGCTCCGGGGCATGGTTGCAAAGCTGAAACTTAAAGGAATTGACGGAAGGGCACCACCAGAGTGGAGCCTGCGGCTTAATTTGACTCAACACGGGGAACCTCAACACCGGCCCGGACACGGAAAGGATTGACAGATTGACGGCTCTTTCTCGATTCTGTGGGTGGTGGTGTATGGCCGTTCGTAGTTACAGGAGCGATTTGTCTGGTTGATTCCGGATAACGAACGAGACTCTGGGCATGCTAACTAGTTACGTGTCTCCGCGCGGTCGGCGTCTGCAACTTAGAGGGACAAGTGGCGTTCTGCCACGCGGAGACTGGAGCAATAACAGGTCTGTGATGTCTCTTAGATGTCCGGGGCTGCACGCGCGCCACAATGGGCGGATCAACGTGCCTACCCTGCGCCGACAGGCGCGGGTAACCTGTTGAACCCCGCCCGTGATGGGGACCGGGGATTGGAAACTATTTCCCGAGAACGAGGAATTCCCAGTAAGCGCAGGTCATCAGCTTGCGTTGATTAAGTCCCTGCCCTTGTACACACCGCCCGTCGCTACTACCGATTGAGCGGCTCAGTGTGTCTCGGATCGGCTCCTGCCTGGGCTCCTTACCGGGGCCCCAGAGCGCCGAGAGAAGACGATCGAACTCGGTCGTTTAGAGGAAGTAAAAAGTCGTAACAAGGTTTCCGTAGGTGAACCTGCGGAAGGATCATTAACGGGGCCGAGGATTCTCCTCCTCACGCCCAGAGGGCGAAGTTCACGGTGCTTTCCGCGCGGTGCGGAAGTCCTCACGGTCTGCCTCCCCACCACGCGCGCGCGAGCGTGATCGGTGATCCAGCGTTGGCGTCGCGGGGCTCCCGGCGGCACCCGGCCGGTCTCGACAATTCTCGACCTGTCCCCTCTGGGTGGGGAGCGACGGAGGTGCGTGGGGGCCGTGGGTTTAAAAGCACTCTTCGCGTTTCACCCGGGGAAGAGAGAGGAGACGCCTGTCCCGGGCCCTGCCGGCCGATGTTTAATTTCCCCACCCCCTCCCGAAGCGTCCTCTGTCTCGGACCGTAACGATTGAAAGAACGAAAACAAGAGTGTACAACTTTAGCGGTGGATCACTCGGCTCGTGCGTCGATGAAGAACGCAATAGCTGCGAGAACTAATGTGAATTGCAGGACACACATTGATCATCGACCTTTCGGAACGTACACAGGCCCGGGTCCATCCCGGGGCCACGCCTGTCTGAGGGTCGCCTGTTATCGATCGACGGGGAAGAGTCGGTCTTCGTGCCGCCCTGCCCTGTCCGCGGCTGGAGCGTCGCAGACCCTGCCCTCCCGCGGTGGCCTACGTCCTCTGGCGCAGACCGCCGAACCGTTCGTCCGCCCGCTTGGGGGCGGCTCCTCGCCGCGCGGTTGCCGGCGGTCTAACAGCTGCCCGCGCGCGGTGGACGGGAGCTGCAGCAACTCCCCTCGCGTTCCGAGACGACGACAGCGCGATCGGTCGACGCGAGACCGGCGTCCCGCCCGTGGGACGACCGGCCACCACCCGCCTGTTGGCCCACGACCTCAGCTCAGGACGAGAAGACCCGCTGAATTTAAGCATATTACTAAGCGGAGGAAAAGAAACCAATCGGGATTCCCCCAGTAGCGGCGAGCGAAGAGTGGAAAAGTCCAGCGCCGAATCCCCGCCCTCTGCCGAGGGCGAGGGACCTGTAGCGGCCGGAGGGCCGCTCTCTCTCGGCGCGGGCCGGGGGCCAAAGTCCTTCTGATGGAGGCTTAGCCCGCGGACGGTGTGAGGCCGGTGTCGGGCCCCGCCCCGCCGGGTGCGGTTCCTCCCGAGTCGGGTTGTTTGGGAACGCAGCCCAAAGCGGCGGTGCAAACTCCATCTAAGGCAATTACCGGTACGAGACCGATAGCGGACAAGTGTACCGTGAGGGAAAGTTGAAAAGAACTTTGAAGAGAGAGTTCAACAGGGCGTGAAACCGTTAAGAGGTAAACGGTGGGGACCGCACCGTCCGCCCGGTGGATTCAGGCCCGGCGGGGCGGGGTCGGCCCGTCCGGTGCGCGCTCCTCTGCTCCCTCATTCCTGGGGTGGCGCGGGGGGTTGACGCCTGGGCGAAGGCTCGGCCGCCGCCGGGTGCATTTCCGCCGCGGTGGAGCGCCGCGACCGGCTCCGTCTGGCTTGAAGGGTCAGGGGGCGAAGGTGGCCCGTCGGTTCAGGCCGTCGGGCTTACAGCGCCCTCCGCCCCGACTTCGCCGCTTGCTCTCCGGGGGCCGCGGTGAGTGTCCTCCGCGCTCCCTCTGCCCCCTCCCTCCCTGGGGAGGTGCGGGGACGGGGTCCCCCGCCTCGGCGTGGCGCGACAGGGGTGGGACTGTCCTCAGTCTGCCCACGGCTGGGTGGCCGCGCCGCCCAGGGCGGGGATCCGACCCACGTTCGCGCCTGAGGTCCGCGGCGACGCCGGCCTCCCACCCGACCCGTCTTGAAACACGGGACCAAGGAGTCCAACGCGCGCGCGAGTCAGAGGGTGGTTCGCGAGCCCTGCGGCGCAATGAAGGTGAGAACGGGGGTCTCCCTGGGCGGGGATCCCCGCCCCGGCGGGGCACCACCGGCCCGCCCCGCGACCCGGGGGCAGGTGGAGCTGGAGCGCGTGCGATGGCACCCGAAAGATGGTGAACTATGCCTGGGCAGGGCGAAGCCAGGGGAAACTTATGGAGGCCCGCCGCGGTCCTGACGCAAATCGGTCGTCCGACCTGGGCATAGGGCGAAAGACCACCGAACCATCTAGTAGCTGGTTCCTCCGAAGTTTCCTCAGGATAGCTGGCGCTCGCCGATCAAGCAGTTTTATCCGGTAAAGCCAATGACTAGAGGCAGGCCTTGGGGCCGAAAACGGCCTCAACTCATTCTCAAACTTTAAATGGTAAGAGGCCCGGCTCGCTGGCGCTGGAGCCGGGCGTGAAATGCGACGCGCCTATGGGCCATTTTGGTAAGCAGAACTGGTGCTGCGGGATGAACCGAACGCTGGTTAAGGCGCCCGATGCCGACGCTCATCAGACCCCATAAAAGGTGTTGGTTGATATAGACAAGGTGATCGGTGGCCATGGAAGCCGGTGGCACCCGCCAAGGAGTGTGTAACAACTCACCTGCCGAATCAATTCAGCCCTGAAATGGATGGCGCTGAGCGTCGGGCCCATGCCCACGTCGACGGCACAAGGACACGCGAGCGGTCGCGCGCGCTGCAAGCCTCGACGCAGGAGGGCCGCCGTGGTGGCGCCGAAGCCCAGGGCGCGGGCCCGCGGAGCGTGCCGCGGCGCAGATCTTGGTGGTAGTAGCAAATATTCAAACGAGGAGCTTTGAAGGCCGAAGTGGAGAAGGGTTCCATGTGAACAGCAGTTGAACATGGGTGGAGCCGGTCCTAAGGGACGGGCCTACGCCGTTCGGCGGGAGGCGATGGCCTCGTCGCCCCCGCTCGACCGAAAGGAGTCGGGTCCAGATCCCCGAGCCCGAGCGGTGGAGTACGGGCGCCGCGAGGCGCCCAGTGCGGTGACGCAAACGAACCCGGAGATGCCGGCGGGCGCCCCGAAAGAGTTCTTTTTCGTGAAGGGCAGGGCGCCCTGGAACGGTTCGCCTCGAGAGAGGGGCCCGCGCCTGAGCGCCGCGCTTCTGGCGGCGTCCGGTGAGCTCTGTCGGCTCTTGAAAATCCGGGAGAAGGTAATCTCCGCGCCGGGCCGTACCTATATCCGCAGCAGGTCTCCATAGAACAGTTTCGGCGTGTTGGAACAAGGCAGAGTAAGGAAGTCGGCAGCTCAGATCCGTAACTTCGGGATAAGGATTGGCTCTAAGGGCTGAGCCGGTCGGGCTGAGGTGCGAAGCGGCCTGGCCCGAGCCGCGACTGGGGAGCGGCCGTCCCGAGTGGCGCTCCTGACCCCGTTCCGGTGCCAGTGGTGCGCGCGCGGTGGCGCGCGGTCCCTCTCTCCCCCGCTGCGTCCGCCCGGGCCTTTAGGAAATTTCGACTCCCCCGACTCTTCCCTCCGCTCTTTCCCCGTCCGCGGGGTCATGGGGGTCTCGGAGGAGGGGGGTGGGAGAGGCCGTGGGGGAGGAAAGGGGTGGTTCGCGTCGGGGCAACGGAGGTCCTCGCCTGCCGCGGGGCGATGCCTCCATGCTGGCCGGGGCCCGCGGGGGCGGATGCGCAGCGGCTGGCGGCGGCGACCCTGGGCGCGCGCCGCGCCCTTCCCGCGGATCTTCCAGCTACAGGCCTCTGCGCCGGGGCCCGCGTTCGCGCGTGCGCCCGGGGAGGCGGCCTCGGCTCCCCGGTGTGGGCGCCTCGGCCGGCGGCTAGCAGCCAGCTTAGAACTGGTCGCGGACCAGGGGAATCCGGACTGTTTAATTAAAAACAGCATCGCGAAGGCCCTCGGCGGGTGTTGACGCGATGTGATTTCTGCCCAGTGCTCTGAATGTCAAAGTGAAGAAATTCAACGAAGCGCGGGTAAATGGCGGAGTAACTATGACTCTTAAGGTAGCCAAATGCCCTTTTCGTCATCTAATTAGTGACGCGCATGAACGGATGAACGAGATTCCCACTGTCCCTACCTGCTATCTAGCGAAACCACAGCCAAGGGAACGGGCTTGGCAGAATCAGCGGAAAGAAGACCCTGTTGAGCTTGACTCTAGTCTGGCCCTGTGAAGAGACATGAGTGGGGTGTAGAATAAGTGGGAGGCCCTGGCTTCCCGGGCCGGCCGCCGTAGAAATACCACTACTTCCATCGTTTCCTCACTTACCTGGTGAGGCGGAAGGCGTTCCAGCCGGCGGGGCCGCCCTGCTTCTGGCGTCAAGCGCCCCGGGGCCGTGCGGGGGTGGGGCAACCCCCCTTCCCCTCCCGGCCGCCGACCGGGGCGCGACCCGCTCCCGGGGACAGCGTCAGGTGGGGAGTTTGACTGGGGCGGTACACCTGTCAAAACGGTAACGCAGGTGTCCTAAGGCGAGCTCAGGGCCAGAACCTCCCGTAGAGCAGAAGGGCAAAGCTCGCTTGATTCTGATTTTCAGTATGAGTACGACCGCGAGTACCTCACGATCCTTCCGGCTTTTGGGGTTTTAAGCGGAGGTGTCAGAAAAGTTACCACAGGGATAACTGGCTTGTGGCGGCCAAGCGTTCATATGACGTCGCTTTTTGATCCTTCGATGTCGGCTCTTCCTATCATTGTGAAGCAGAATTCACCAAGCGTTGGATTGTTCACCCACTAACAGGGAACGAGCTGGGTTTAGACCGTCGAGACAGGTTAGCCTTACCTCACTGATGTGAGCCGTTGTTGCAATAGTAATCCCGCTCAGTACGAGAGGAACTGCGGGGTTCAGACATTTGCCCGTGTGCTTGGCTGGAGGAGCCACTGGCGCGAAGGTTTCACCATCTGCGGGATTATGACTGAACGCCTAAGTCAGAATCCCGCCTGCGCCAAACGATACAGCAGCGCCGTCGCATTCCGATAGGCCCTGGGTAACCGGGAAGATTCTCTCCGGGGGGGCCCCCGGCGCGGAGAGCTATTCGTGACAGGAACCGGGGCGCGGCTAGAACGAGCGCTGCCCCCTCTTTCCAGTGACGTAACCGCATGTTTGTGGGGGAACCGGTGCTTAAATGACTCGTAGACGACCTGATTCTGTCGCGTCGGTGCAGAGCAGCTCTGCTGCTGCGATCCATTGAAAGTCAGCCCTCGATCCAAGTTTGTCACCGGACGCAGGTGGCGACAGGGCCACCGGTCCGACAGACCCGCGGCGTGCGAAGGGGCACCAGCTGCAAGTCGGTGAAGGCTTTTCAGCCATCCCCTCGCCACTTGCGCGCAGGCCGAGGCACCAGCGGCGGAGCGTGGGCGCGATCCCAATACACCTGGGGCACCAGCGGCAGAGGGAGGCCCAGCCTCGGCCCGCACCTTCACAGCAAGCCCCTACTGGGGCACCAGCAGCACGCAGTACCTATGTGTGCACCAGCAGAAGGGGTTCTGTCCAGACCTCAACCCCAGAGCAGACTTTCTTTATTAAAACAGCAGCAAATGCAGCACCCACGTGGGTCGCCACAGCATAGCAACCAGCCTGCCCCGCACCCCACAGCAAAGCTTAGGGCACCAGCAGCACGCGTACCTACATGTTTACCAGCGCAGAAGAGGTCCGGACCCGACCCACACCCCACAACAGCCTTCGATGGGGTACCAGCAGCACGCAGTACATAAATGGGTCACCAGCGGCGAGTCTCGCAGCGCGCGCCTTACCGAAGTCAGTGTCCCTGGCTTATATGGGTTCCGGGCACGGCTGGTAACCGGTTCACGGTGGGGTCAGGAGGCTTCGCCGGTGCGTCCCGGCCCAGCCTGGGGCCGCGCCAGTCAGTCAGCCCGGCCCGTTCAGCCCAGCTCTGTAGCCAGCCAGCCAGCCAGCCAGCCAGCCCAGCCCAGCGCTGTCCAGCCAGCCCCGTCTGTGCCCAGCTCAGGCGCCCCAGGCCGGCCGGCCAGCCAGCCAGCCGCAAGCTCCAGGCCAGCCAGGCCCAGCCAGCCCAGCCAGCCCTGGGCCCTGCCCCAAGCCCTGCCCCAGCCCAGCCCAGCCCAGGCCGGGCCAGCCAGCCAGCCAGCCAGCCAGCCAGCCAGCCCAGGCCAGGCCAGCCAGGTTGTGTTGAGCCAGGGCCAGGCCAGGCCAGCCCAGCCTGGCCAGGCCAGGCCCAGCCTAGCTGCGCCTCCAGTCGGGCCAGCCAGCCCAGCCCAGTTGTTAAAGCCAGCCAGCCAGCCAGCCCAGCCCAGCCCAGCTCAGGCAAGCCCAGGCCGGCCAGCCAGCCAGCCAGCCAAAGTCAGGCCAGGCCCAGCCCGCCTGGGCCAGCCAGCCGAAGCCCAGGGTGGCTGTTGCAGGCCAGCCAGCCAGGCCAGGTCAGGCCAGCCAGGCCAGCCAGGCCCAGCCCAGCCGGGAGGCCAGCCCAGCCGAGCCCAGCCAGCCAGCCAGCCAGCTGTAGCCCAGCCTGTGTTCAGCCCAGCTCCCGCCCTGCCCTGCCCTGCCCTGCCCTGCCCAGCCCAGCCCAGCCCAGCCCAGCCAAGCCCAGGCCGGCCAGCCAGCCAGCCAGCCAGCCAGCTCACAGGCTGGGCCAGGCTGCAGGCCAGGCCAGGCTAGAAGCAGGCCCAGCCCAGCCAGCCAGCCAGCCACTGAAGCCAGCCCAGGCCAGGCCAGGCCAGGCCATGCCCAGCCCAGCCAGCCAGCTCGAGCCCAGGAGTCATGCTGCTCCTGGCTGGGCAGCAAGCCCAGCTCAGCCAGGGCCAGCTTCCAGGCCAGGCAAGCCCTGCCGCCAGCCAGCGAAGCCCAGCCAGCCAATCAGTCAGTCAGCCGCGCCAGTCAGTCAGGTCAATCAGCCCAGCCCAGCCCAGCCCAGCCTGCCTTGGGGCCCAGCCCAGCCCAGCCCAGCCCAGCCCAGCCCTGCCCTGCCCAACCTGACAGCCAGGCCTGCCTAGCCAGCCAGCCAGCCATCCTGGTGTTGGAGAGGCTTGGGGCTTTAGAGTGGCTGGTCGACAGTCCAGTTGGCCGGAGGGAGAGGAGCACTATCTCCTCCCACTTCTCCTCTTCCTTACCACCACCACCACCACCACCACCACCACCACCACCACCACCACCACCCCGGAGGCCCGGGGATTGGGCTTGGGGATAGGGTAAGGGTTGCCTGGAGGCTTGGGGGTGGGTTAGGCTGCCCGGAGGCCCGTGGGATTGGGCTTGGGGATAGGGGCAAGGGGCTGCCTGGAGGCTTGGGGGTGGTTAGGCTGCTCCGAGGCCCGGGGATTGGGCTTGGGGATTAGGGTGGGCGGTTGCCTGGAGGCTTGGGGGTGGGGTTAGGGTTGCCACAGGAGGCCGGGGATTGGGCTTGGGCACAGGGTAAGGGCTGCCTGGAGGCTTGGGGGTGGTTAGGGCTGCCTGGAGGCCCGGGATTGGGGCTTGGGGATAGGGCAAGGGCTGCCTGGAGGCTTGGGGTGGGTTAGGGCTGCCAGGAGGCCCGGGGATTGGGCTTGGGCATAGGGTAAGGGGTTGCCTGGAGGCTTGGGGTGGGTTAGCTGCCTGGAGGCCCGGGGATTGGGCTTGGGCATAGGGTGGCGGTTGCCTGGAGGCTTGGGGGTGGGGTTAGGCTGCTCTGGAGGCCCGGGATTGGGCTTGGGGATAGGGTGCTGGGTTGCCTGGAGGCTTGGGGTGGGTTAGTCACAGGGAGGCCCGGGGATTGGGCTTGGGCATAGGTAAGGGTTGCCTGAGGCTTGGGGGTGGGTTAGGGCTACCTGGAGGTCTGGGGATTGGGCTTGGGGATAGGGCAAGGGTTGCCTGGAGGCTTGGGGGTGGGTTAGGGCTGCCAGGAGGCCTGGGGATTGGGCTTGGGCATAGGGTAAGGGTTGCCTGGGAGGCTTGGGGTGGGGTTAGGCTGCCTGGAGGCCCGGGGATTGGGCTTGGGGATAGGGGTAAGGGTTGCCTGGAGGCTTGGGGTGGTTAGGCTGCCTGAGGCCTGGGATTGGGCTTGGGGATAGGGTAAGGGTTGCCTGAGGCTTGGGGTGGGTTAGGGCTGCCAGGGGAGGCCCGGGGATTGGCTTGGGCATAGGGTAAGGGTTGCCTGGAGGCTTGGGGGTGGGTTAGGCTGCCTGGAGGCCCGGGATTGGGCTTGGGGATAGGGTAAGGGTTGCCTGGAGGCTTGGGGGTGGGTTAGGGCTGCCAGGAGGCCCGGGGATTGGGCTTGGGCATAGGGTAAGGGCTGCCTGGAGGCTTGGGGTGGGTTAGGGCTGCCAGGGAGGCCCGGTGGATTGGCTTGGGGATAGGGTAAGGGCTGCCTGAGGCTTGGGGTGGGTTAGGGCTGCCAGGAGGCCCGGGATTGGGTTTGAGATAGGGTAAGGCTGCCTGGCACTTGGGGGTTAGGGTTAGCTGCCCGGAGGCCTGGGGATTGAGCCTGGGGATAGGGTGGCTGCCTGGAGGCTTGGGTGTTAGGGCTAGGGTTTCCGGAGGACTGGGAATTGGGTTAGGTTGCCTGGAGACTTGGGGTGGGTTAGTTGCCCGGAGGCCTGGGGATTGGGCTTGGGGATAGGGGTTAGGGCTGCCCGGAGGCTTAGTTGTTGGGGCTTGTGGAGCTTTGCTTGCCTGGTTGCCTGAGGCTTGGGAGAAGTGCCACCCCGCGTGCCAGCCAGAAAGGTCAGCCAGTGACCAGCACTGCCCGGCTGCCGAAGGTAAAGTCAGTCGCGGGTCACCATCGGCACCCGATTCGGCAAGGAGAAGCAAGAAGGGCCAAGCAACCCTGACTATCCATGGCCCCCCGCGCTCAGGTTCACAGCCGGGCGGGACGTGACCTCTGGGTCTCTCTCCCGGAGGTCTGGGCAGCCGTCACTGGTTATCGCTGACTCGTTGCCGAGGTTTCGGGAAGGCGTGACTTAGTGGACTTAGGTTTTGAGCGCAACTACGTGCCACCGCCTTCCACCACCTGACCGATGGAATTGTGGCGCGATGCCCCAGGGGCAGGCTCCCTTGCCCGGTCCTCCGACAAAAGGGTAATCGCTGGGGTGGGGTGGGTTCCTCCCCGTTTTGGGCATTCGAAAGGAGGGCGAGGCGGGCCTTCACCAGACCTCGACCCTTGCTCCTTTGGGTTAAAGACTCAGATGGCCCGTCGATCTCAGACCGTTCTGTACCGACGGAACAAGTGCGCTGTCGGTGCTGTGCGTGCGTTTACCGGGCAGCATCCTTCGCCCCCTCCTTCTCCTCACCACCCGGGTGTAATCCCGGGGCGCGTAGGCTGACTTGGGGGTGGGGAAGCGTGGTGGCGCGCGCGGTATCTGCGGCGCACTCGATCCGTTGCGAAGCACGATTCTTCCTGGTGATGGCCCGACCCAGCGGCCAGCCCAACGCCGGCAGACCCTCATTTCCATGTGTGCGGTGGGGGTGGGTAGGCGGCGGCACTCTGGAGCCCAATCCCTCTCTCCCACTTAGCCCGCGTGCCCCTCCTGCGAGCAGTCGCACGCCTCGGCGGAGCGAGCCCGCTTGTGGTAGGCGGCGCGGAGCTGTGGTGCGGCGGGAGTGGTTCTTCCGCCGAGTCCTTCCCCCTCTCTCCTCCGGCACTCGGGCCTTTACCACCACGCGATTCACCCGAGGCGAGGCTACCTGGTTGGGATCCTGCCAGTAATATATGCTGTCTCAAAGATTAAGCCACAAGTCTAAGTGCACACGGCCGGTACAGTGAAACTGCGAATGGCTCATTAAATCAGTTATGGTTCCTTTGGATCGCTCCACCCGGTTACTTGGATAACTGTGGCACTCCAGAGCTAATATGCCAACGAGCGCCGACCTGCTGCCCTTCCCTCGGGGCGGGGGGGGTGGCACCCGGGGACGCGTGCATTTATCAGGATCCAAAACCCATGCGGGTGCGCGGGCGGTGTGAGAGGGGGCCCGCGCCTACCCGCCGCCGTCCGCCCCCGGCCTCGCTTTGGTGATTCCCAGATAACCTCGCCGATCGCGTTCTCGCGGCGGCGACGGTCTATTCGAATGCGTCTGCCCTATCAACTTTCGGGATGGTAGGTCCGTCGCCTACCATGGTGACCACGGGTGACGGGGAATCAGGGAAGTTCGATTCCGAGAGGGAGCCTGAGAAACGGCTACCACATCCAGTTAAAGCAGCAGGCGCGCAAATTCACCCATTTCCGACACGGAGAGGTAGTGACGAGAATAACAATGCAGGTCTCTTTCGAGGCCCTGCAATTGGAATGAGTGCATCCCAAACCTATGGGCGAGGACCCATTGAGGGCAAGTCTGGTGCCAGCAGCCGCGGTAATTCCAGCTCCAATAGCGCATGCTAACGTTGCTGCAGTTAAAAAGTCAGGCTGGGATCTCGGGGACCGGGGTCATGCGCGGTCCGCCGCGAGGCGAGCCACCGCCGGTCCTGACCCCTTCAGGCCTCCCGGCGCCCCCCGGATGCCCTTGACTGCGTCCTCGGCTTCTAGCTCCGGAGCGTTTACTTTGAAAAATTAGAGTGTTCAAGGTGGCCGGCACCGCGCCCCATTGAATACCCCAGCTAGAATAATGGAATAGGACCCCGGTTCTATTTTTCAGGGTTTCCGGAACCCGGGGGCCATGATCGAGAGGACGGCCGGGGGCATTCGTATTGCGCCGTTAGAGGTGAAATTCTGACCGGCGCAAGACGGACTGGAGCGAAAGCGTTTGCCAAGAACGTTTTCATTAATCAAGAACGAAAGTCGGAGGTCTGGAAGACGATCAGATACCGTCGTAGTTCCGACCGTAACGATGCCGACCATGATCCGGCGGCGTTTATTCCCATGACCCGCCGGCAGCGTTGCGGGAAACCACGAGTCTCTGGGCTCCGGGGAGTATGGTTGCAAAGCTGAAACTTAAAGGAATTGACGGGAAGGGCAAATTCACCAGGAGTGGAGCTCGCGGCTTAATTTGACTCAACACGGGGAACCTCACCCGGCCCGACACGGAAAGCGATTGACAGATTGACGGCTCTTCGATTCTAGTGTGCATGGCCGTTCGTAGTTGGTGGAGCGATTTGTCTGGTTGATTCCGATAACGAACGACTCTGGCATGCTAACTAGTTACGCGCCCCGCGCGGTCGGCGCCGCGCAACTTTAGACAAGTGGTTCAGCCACGCGAGACTGAGCAATAACAGGTCTGTGATGCCCTTAGATGTCCGGGGCTGCACGCGGTGCCACAATGGGCGGATCAATGTGCCTACCCTGCGCCGACAGGCGCGGTAACCCGTTGAACCCCGCCCGTGATGGGGACCGGGATTGAAACTATTTCCCGAGAACGAGGAATTCCCAGTAAGCGCAGGTCAGCTTGCGTTGATAAGTCCCTGCCCTTTGTACACCGCCCGTCGCTACTACCGATTGGAGCGGCTCAGTGAGGTCCCCGGATCGGCTCCCGCCCGGGGCCCTTTACCGGGGCCCTGGTGGAGCGCCGAGAAGACGATCGAACTCGGTCGTTTAGGAGGAAGTAAAAAGTCGTAACAAGGTTTCCGTAGGTGAACCTGGGTGGGATGATCATTAACGGGGTCGAGGGGATCTTCCTCACGCCCAGAGGGCGAAGCCACGGTGTTTCCGCGCGGTGTGGAAGTCCCTACGGGTCTACCTCCCCACCATCGCGCGCGCGAGTGCGATCGGTGATCCAAAGGTTGGCGTCGCGGGCTCCCGGCGGGTACCTGGTTATCTCGACCACCTCGACCTGTCCCCTCTGCGGGGGAGAGCGACGGAGGTGCGTGGGGGCCGTGGGTTTAAAGCACTCTTCGCGTTTCCTCACCCGGGGAGCAGAGGAGACGCCCGTCCCGGGGCCCTGCCGGCCGATGTTTAATTTCCCCACCCCTCCCCCGAAGCGTCCTCTGTCTCGGACCGTAACGATTGAAAGAACGAAAACAAGAGTGTACAACTTTAGCGGTGGATCACTCGGCTCGTGCGTCGATGAAGAACGCAAAGGCTGCGAGAACTAATGTGAATTGCAGGACACACATTGATCATCGACCTTCGAACGCACACAGGCCCCGGGTCCATCCCGGGGCCACGCCTGTCTGAGGGTCGCCTTGCTATCGATCGGACGGGGAAGAGTCGGTCTTCGTGCCGCCCTGCCCCGTCCGCGGCTGGAGCGTCGCAGACCCTGCCCTCCCGCGGTGGCCTACGTCCTCCTGCGCAGACCGCCGAACCGTTCGTCCGCCCAAGGCTTAAAGTGGCCCTACTTCCCTCTCCCTCGCGCGGCTGCGCCGGCGGTCTAACAGCGCTGCCCGCGCGCGGTGGACGGGAGCTGCAGCAAACTCCCCTCGCACTCTGAGACGACGACAGCGCGATCGGTCGACGCGAGACCGGCGTCCCGCCCGCGTGGGACGACCGGCCGCCACCACCCGCCTGTTGGCCCACGACCTCAGCTCAGACGAGAAGACCCGCTGAATTTAAGCATATTACTAAGCGGAGGAAAAGAAACCACCGACTCCCCCAGCAGCGGCGAGCGAAGAGGGAAAAGTCCAGCGCCGAATCCCCGCCCTCCGGGGCCGAGCGAGGGACCTGTGGCGTACGGAGGGCCGCTCTTCCGGCGCGGGCCGGGGGGCCAAAGTCCTTTCCTGATGGAGGCTTAGCCTGCGGACGGTGTGAGGCCGGTGTCGGCCCCCCGCCCCGCCGGGGTGCGGTTCCTCCCGGAGTCGGGTTGTTTGGGGAATGCAGCCCAAAGCGGGTGGTAAACTCCATCTAGCTAGCTAAATACCGCACGAGACCGATAGCGGACAAGTACCGTGAGGAAAGTTGAAAAGAACCTTGAAGAGAGAGTTCAACAGGCGTGAAACCGTTAAGAGGTAAACGGTGGGGACCGCACCGTCCGCCTGGTGGATTCAGCCCGGCGGGCGGGGTCGGCCCGTCTGGTGCGCGCTCCCTTCGCTCCCTCATTCCTGGGGGTGGCGCGGGGGTTGACGCCCGCGAAGCGCTCGGCCGCCGCCGGGTGCATTTCCGCCGCGGAGCGCCGCGACCGGCTCCGGTTCGGTTTGAAGCGGGTCAGGGGGCGAAGGTGGCCTGTCGGTTCAGGCCGTCGGGCTTACAGCGCCCTCCCGCCCCGACTTTCGCCGCTTGCTCCCTCCGGGGCCGCGGGTGAGTGTCCTCCGCGCCCTCTCTGCCCTCCCTCCCTGTGGGGGAGGTGCGGGGACGGGGTCCCCCGCCCCCGGCGTGGCGCGACAGGGGTGGACTGTCCTCAGTCCGCCCACGGCTGCGCCGCGCCGCCCAGGGCGGGGATCCGACCCACGTTCGGGCGCGCCCGAGGTCCGCGGCGACGCCGGCCTCCCACCCGACCCGTCTTGAAACACGGACCAAGGAGTTCCAACGCGCGCGCGAGTCAGAGGGTGGCTCGCGAGCCCTGCGGCGCAATGAAGGTGAGAGAACGGGGGTCTCCCTGGTGGATCCCCCGCCCCGGCGTGGGGCGCACCACCGGCCCGCCCCCGCGACCCCCGGGGGCAGGTGGAGTTGGAGCGCGCGCGATGGCACCCGAAAGATGGTGAACTATGCCTGGGCAGGGCGAAGCCAGGGAACCCGGTGGAGGCCCGCCGCGGTCCTGACGTGCAAATCGGTCGTCCGACCTGGGCATAGGGAATAGTGATCATTAATCGAACCATCTAGTAGCTGGTTCCTCCGAAGTTTCTCAGATGGGCTGCTGGCGCTCGCCGATCAAGCAGTGTTTATCCGGTAAAGCCAATGACTAGAGGCCTTGGGCCGGGAAACGGCCTCAACCTATTCTCAAACTTTAAATGGTAAGAGGCCCGGCTCGCTGGCGCTGGAGCCGGGCGTGGAATGCGACGCGCCCAGTGGGCCATTTTGCAAGCAGAACTGGTGCTGCGGGATGAACCGAACGCCGGGTTAAGGCGTTCGGATGCCGACGCTCATCAGACCCCATAAAAGGTGTTGGTTGATATAGACAGCAGGACGGTGGCCATGGAAGTCGGCACTGCCAAGGAGTGTAACAATCACCTGCCGAATCAACTAGCCCTGAAAATGGATGGCGCTGGAGCGTCGGGCCCATACCGGCCGTCGACGGCACAAGGGACACGCGAGCGGTCGCGCGCGCTGCAAGCCCGACGAGAAGGGAGGCTGCCGCGGTGGCGCCGGCCCAGGGGCGGGCCCGGTGAGCGCTGCGGGCGCAGATCTTGGTAGTAGCAAATATTCAAACGAGAGCTTTGAAGGCCGAAGTGGAGAAGGGTTCCAGCTGTGAACAGCAGTTGAACATGGGTGAGCCGGTCCTAAGGGACGGGCCTACGCCGTTCGGAGGGAGTGGGGCGATGGCCCTGTCGCCCCGCTCGACCGAAAGTGAGTCGGGTCCAGATCCCCGAGCCCGGAAAGTGAAGGGAGACGGGCCGCGAGGCCCAGTGTGGTGACGCAAACGAACCCGAGATGCCATGGTGGGCGCCCTGAAAGAGTTCTCTTTTTCTTTTGTGTGAAGGGCAGGGCGCCCGAACGGGTTCGCCCCGAGAGAGGGGCCCGCGCCTCTGGAAAGCGCCGCGCTTCTGGCGGCGTCCGGTGAGCTCGTCGGCCCTCTGAAAACTGGGGAGAAGGTAAATTCGCGCCGGGCCGTACCCATATCCGCAGAAGTCTCCAAGGTGAACAGCTTCTGGCGTGTTGAACAAGGCAGAGTATGGAAGTCGGCAAGTCAGATCCGTAACTTCGGGATAAGGATTGGCTCGCGTGGCTGAGCCGGTCGGCCAGGTGCGGCGGGCCTGGGCCCGGAGCTGCGACTGGGGAGTGGCCGCCCTGAGGTGCCCTGACCCCGTTCCCGAGCCGTGGGGGCGCGCGGTGGCGCGCGGTCCCTCTCTCCCCGCCGTGTCCGCCCGGGCCTTTTTGCGTCTCGACTCCCCCGACTCTTCCCCTCTGCCCTTTCCCCGTCCGCGGGGGTCGGGCGGGGGTCTCGGAGGAGGGGGTGGAGAGGCCGTGAGGAAAGGGGTGGTTCGTGGCGTCGGGGGCAACGGAGGGTCCTCGCCTGCTGCGCGATGCCTCCGCTGGCCGGGGCCCGCGGGGGCGGGATGCGCAGCGGTTGGCGGCGGCGACCCTGGGCGCGCGCGCCGCGCCCTTCCCCGCGGATCTCCGCAGCTACGGCCCCGCGCCGCCTGCGGTCTCTGGCGGCGGCGCTCTTCTCCGGGGAGTGGCGCGGCCTCGCTCCCCGGTGCGGGCGCCCGGCCGGCGGCTAGCAGCCAGCGAACGTCGCAGGACCAGGGAATCCGACTGTTTAATTAAAACAAAGCATCGCGAAGCGCCCTCGGCGGGTGTGCTGACGCGATGTGATTTCTGCCCAGTGCTCTGAATGTCAAAGCGAAGAAATTCAACGAAGCGCGGTAAACGGCGGGAGTAACTATGACTCTCTTACAGTAGCCAAATGCCTCGTCATCTAATTAGTGACGCATGAATGGATGAACGAGATTCCCACTGTCCCTACCTGCTATCTAGCGAAACCACAGCCATGGAACGGGCTTGGCAGAATCAGCGGGAAAGAAGACCCTGTTGAGCTTGACCTAGTCTGTCTTTATAGAAGAGACATGAGGGGTGTAGAATAAGTGGAGGCCCCCTGGCCTCCCGCCGGCCGCCGGCAATACCACTAGCCTTTATCGTTTCACTTACCTGGTGAGGCGGGGAAGCCGGGCTGCCCGCCCGGCGGGGGCCGCCCTGCTTCTGGGGCGTCAAGCCCGGGCCGTGCGTGGGCAACCCCTCCCCTCCCGGCCGCCGACTGGGGCGCGACCCGCCCCGGGACAGCGTCAGGTGGGGAGTTTGACTGGGGCGGTACACCTGTCAAACGGTAACGCAGGTGTCCTAAGGCGAGCTCAGGGGGACAGAAACCTCCCGTAGAGCAGAAGGGCAAAAGCTCGCTTGATCTTGATTTTCAGTATGAGTACGGACCGCAGCGGGGGCCTCACGATCCTTCCGGCTTTTTGGGGTTTTGGCGGAGGTGTCAGAAAAGTTACCACAGGGATAACTGGCTTGTGGCGGCCAAGGTTCATAAAGGCGACGTCGCTTTGATCCTTCCCGATGTCGGCTCTTCCTATCATTGTGAAGCAGAATTCACCAAGCGTTGATTGTTCACCCACCAACAGGAACGTGAGCTGGGTTTAGACCGTCGTGAGACAGGTTAGCTTCACTCCTACTGATGTGAGCCGTTGTTGCAATAGTAATCCCGCTCAGTACGAGAGGAACCGCGGGTTCAGACATTTGGTTCGTGCGCTTGGCTGAGGAGCCACTGGCGCGAAGCCACCATCTGCGGGATTATGACTGAACGCCTCTAAGTCAGAATCCCGCCTAAAAGCAACGATACAGCAGCGCCGTCGGGATCTCCGATAGGCCCTGGTAACCGGAAGCCCTCTCCGGGGGCCCCGGCGCGGAGAGCCATTCGTGACAGGAACCGGGGCGCGGCTAGAACGAGCGCCGCCCTCTTCCAGTGACGCACCGTACGTTTCAGGGGGAACCCGGTGCTTAAATGACTCGTAGACGACCTGATTCGGGTCGGGGTGGCTGTGCGTGGCAGAGCAGCCCTGTTGCTGCGATCCATTGAAAGTCAGCCCTCGATCCAAGTTTTGTCGGGCCGGACGCAGGCGACAGGGGCTACCGGTCCGACAGACCCGCGGCGTGCGAAGGGGCACCAGCTGCGTCGGTGAAGGCTTCTGCCATCCCCCTCGCCACTTGCGCAGGCCGAGGCACCAGCGGCGGGAGCGCGGCGCGATCCCAATACACCTGGGGCACCAGCGGCAGAGGGAGGCCCAGCCTCGGCCCGCACCTCACAGCAAAGCCCCTACTGGGGCACCAGCAGCACGCGTACCTATGTGGGTAACCAGCGGCAGAAGGGGTTCTGTCCCGACCCTAACCCCACAGCAGAGCCTTTATTGGGGCACCAGCAGCACGCAGTACCCACGGGTCGCCAGCGGCAGAAGGGGTCCAGCCTGCCCGCACCCCACAGCAAAGCCTTTAGTAAGCACCAGCAGCACGCAGCACTACATGGGGTTACCAGCGGCAGAAGAGGTCCGACCCGACCCACACCCCACAACAAAGCCTTCGATGGGGTACCAGCAGCACGCAGTACATAAATGGGTCACCAGCTATGCGTGAGCTGGCGCGCGCCTACCGAAGTCAGTGTCCCCGGGCTTATGAGTGGGTGCCCGGGCACGGCTGGTGAACGGGTTCCATGGTGGGGTCAGAGGCCGGCTGGTGTCCCCGGCCTAGCCAGCCAGCCAGTCAGTCAGCCCGGCCCAGCCCAGCCCAGCCCAGCCAGCCAGCCAGCCAGCCAGCCAGCCCAGCCCAGCTGTCCAGCCAGCCCTGTCCAGCCCAGTCCAGGCAAGCCCAGGGTCGTGGTCGGCCACAGCCAGCCAGCCAGCAAGCCCAGGCCAGGCTCAGGCCCAGCCAGCCCAGCCAGCCCTGCCCTGCCCTGCCCCTGCCCTGCCCAGCCCAGCCCAGGCCGGCCAGCCAGCCAGCCAGCCAGCCAGCCAGCCAGCCCAGGCCAGGCCAGGGCCAGGCCAGGCTGTAGCCACAGGCCAGGCCAGGCCAGGCCCAGCCCGGCTGTGGCCAGGCCCAGCCCAGCCAGCCCAGTCAGCCAGCCAGCTCCAGCCCAGCCAGCCAGCCAGCCAGCCAGCCAGCCCAGCCCAGCCCAGCCCAGGCAAGTTCCCAGGCCGGCCAGCCAGCCAGCCAGCCAAGGCCAGGCCAGGGCTCAGCCCAGCCAGCCAGCCAGCTAGCCCAGGTACAGGCCAGGCCAGGCCAGGCCAGGCCAGGCCAGGCCAGGCCAGGTTAGCCCAGCTCCAGCCAGGCCAGCCTGCCGAGCCCAGCCAGCCAGCCAGCCAGCCAGCCCAACAGCCTGAGCCCAGCCCTGCCCTGCCTTGCCCTGCCCTGCCCTGCCCAGCCCAGCCCAGCTCAGCCCAGCCAAGCCCAAGGCCGGGCCAGCCAGCCAGCCAGCCAGCCAGCCCAGGCCAGGCCAGGCCAGGCCAGGCCAGGCCAGGCCAGGCCAGGCCCAGCCCAGCCAGCCAGCCAGCCAGCCAGCCAGCCCAGGCCAGGCCAGGCCAGGCCATGCCCAGCCTCCAGCCAGCCAGCCAGCCCAGGCCAAGGCAAGCCCTGGCTGGGGCAGCAAGCCCCAGCTCAGCCAGCCAGCCCAGGCCAGGCAAGCCCTGGCCGGGCCAGTGCCAGCCGCCCAGCCAGCCAACAGTCAGCCGTCAAGGCCAGTCAGTCAGGTCAATCAGCCTAGCCCAGCCCAGCCCAGCCTAGCCCAGCCCAGCCCAGCCCAGCCCAGTCCAGCCTAGCCTCCTGCCTCGCCCACCTGACAGCCAGGCCTGCTCAGCCAGCCAGCCAGCCATCCTGGTGTTGGAGAGGCTTGGGCTTTAGGAGTGGCTGGTCGATAGTCCATGGCCGGAGGGAGAGTGAGCACTATCTCCCCTCCCCCACTTCTCCTCTTCCTTACCACCACCACCACTCACCACCACCACCACCACCACCACCACCACCCCGGAGGCCCGGGGATTGGGCTTGGGGATAGGGTAAGGGCTGCCTGGAGGCTTGGGGTGGGAGTTAGGGCTGCCCGAGGCCCGGGGATTGGGCGGGATAGGGCAAGGGTTGCCTGGAGGCTTGGGGGTGGGTTAGCGTCTGGAGGCCCGGGGATTGGGCTTGGGGATAGGGCAAGGGTTGCCTGGAGGCTTGGGGGCGGGTTAGGGCTGCCAGGAGGCCCGGGATTGGGCTTGGGCATAGGTGGGCGGTTGCCTGGAGGCCTGGGGGTGGGTTAGGGCTGCCTGGAGGCCCGGGGATTGGGCTTGGGGATAGGTAAGGGTTGCCTGGAGGCTTGGGTGGTTAGGCTGCCAGGAGCTCCGGGGATTGGGCTTGGGCATAGGGTATGGGCCTGGAGGCTTGGGGGTGGGGTTAGGCTGCCTGGAGGCCCGGGGGATTGGGCTTCTGGGGCATAGGGCAAGGGCTGCCTGGAGGCTTGGGGTGGTTAGGGTTGCCTGGAGGCCCGGGGATTGGGGCTTGGGGATAGGGTGTGGTTGCCTGGAGGCTTGGGGGTGGGTTAGGCTGCCAGGAGGCCCGGGGATTGGGCTTGGGCATAGGGTAAGGGTTGCCTGGAGGCTTGGGTGGGGTTAGGCTGCCTGGAGGCCCGGGGATTGGGCTTGGGGATAGGCAAGGGTTGCCTGGAGGCTTGGGGGTGGGGTTAGGGGCCGCCAGGAGTCTGGGATTGGGCTTGGGCATAGGGGTAAGGTTGCCTGGAGGCTTGGGGTGGGTTAGGCTGCCTGGAGGCCCGGGGGATTGGGCTTGGGATAGGGTAAGGGTTGCCTGGAGGCTTGGGGTGGGTTAGGGGCTGCCTGAGGCCCGGGGATTGGGCTTGTGATGGTAGCAGCTGCCTGGAGGTTGGGGTGGGTTAGGCTGCCAGGAGGCCCGGGGGATTGGGCTTGGGCATAGGGTAAGGGTTGCCTGGAGGCTTGGGGTGGGGTTAGCTGCCTGGGAGGCCCGGGGATTGGGCTTGGGGATAGGGTAAGGGTTGCCTGGAGGCTTGGGGGTGGGTTAGGGCTGCCAGGAGGCCCGGGGATTGGGCTTGGGCATAGGGCAAGGGTTGCCTGGAGGCTTGGGGGTGGGTTAGGTTGCCAGGGAGGCCCGGGGGGATTGGGCTTGGGGGATAGGGTAAGGTTGCCTGAGGCTTGGGGGTGGGTTAGGGCTGCCAGAGGCCCGGGATTGCTTTTGGAGCATAGGGTAAGGGTTGCCTGAGGCTTGGGGGGTTAGGTTAGGGTTGCCCGGAGGCCTGGGATTGAGCTTGGGGATAGGGTAAGGTTGCCTGAGGCTTGGGTGTTAGGTTAGGTTTCCCGGGAGGACTGGGAATTGGGTTAGGTTGCCTGGGAGACTTGGGGGGTTAGGCTGCCCGAGGCCTGGGGATTGGGCTTGGGGATAGGGTTAGGTTGCCCGGAGGCTTGCTGTTGGGCTTGTGGAGCTTTGGCTTGCCCAGTTGCCTGAGGCTTGGGAGAAGTGCCACCCGCGTGCCAGCCAGTGGTCAGCTGGTACCAGCAGCAATTGCCTGGCTGCCGAAGGTAAAGTCAGCTCGCGGTCACCATCGGGCACCCGATTCGGCAAGGAGAAGCAAGAAGCGGCCGCTAACCCTGACTATCCATGGCCCCCGAGCTCAGGTTCACAGCCGGGCGGGACGTGACCTCTGGGTCTCTCTCTCCCGAGGTCTGGGCAGCCGTCACTGGTTATCGCTGACCTGCCGTCGAGGTTTCGGGGAAGGCGTGATTTAGTGGACTTAGGCTTTGAGCGCGTAACTACGTGCCACCGCCTCCACCACCTGACCGATGGGAATTGTGCGCGATGCCCCAGGGCAGGCCCCTGCCCGGTCCTCCGACAAAGGGGACCGCTGTGGGTGGGGTGGTTCCTCCCGTTTTGGGCATTCGAAAGGAGGGCGAGGCGGGCCTTCACCTGCGACCTCGACCCTTGCTCCTTTGCCAAAGACTCAGATGGCCCGTCGATCTCAGACCGCCCTGTGGCCGACGGAACAAGTGCGCTTAGTCGGTGCCAGTGCCTACCGGGCAGCATCCTCCTGCTCCCCCTCCTTCTCTCCTCACCACCCGGTAATCCCGCGCGTAGGCTGACTTGGGGGTAGGTTGTGTGGTAGTGGCGCGCGCGGTATCTGCGGCGCACTGATCCCGTTGCGAAGCACGATCCTTTCCTGGTGATGGCCCGACCCAGCGGCCAGCCCACGCCGGCAGACCCCTCATTTCCATGTGTGCGGTGGGGGTGGTAGGCGGCGGTACCTGGAGCTCAATCCCCTCCCACTTAGCCTGCGTGCCCTCCTGCGAGCAGTCGACGCCTCGGCGGAGCGAGGTCCGGGCTTGTAGGCAGGCGGCGCGGAGCTGTGGTGCGGCGGGAGGTTCTCCGCCGAGTCCTTCCCTCTCTCCTCCGGCACTCGGGCCTTTTACCACCACGCGATTCACCCGAGGCGAGGGCTACCTGGTTGATCCTGCCAGTAATATATGCTTGTCTCAAAGATTAAGCCATGCAAGTCTAAGTGCACACGGCCGGTACAGTGAAACTGCGAATGGCTCATTAAATCAGTTATGGTTCCTTTGATCGCCCACCCGGTTACTTGGATAACTGTGGCAATTCCAGAGCTAATACATGCCAACGAGCGCCGACCTGGGCCGCCCTCTCCTCGTGGGGCGGGGGTGGTACCCGGGGACGCGTGCATTTATCAGATCCAAACCCATGCGGTGCGCGGCGGTGAGGGGGCTCAGCTCACCTGCCGCCGTCCGCCCCGGCCTCGCTTTGCGACTCTAGATAACCCGGGCCGATCAGTACGCTCCTCGCGGCGACGGTTCATTCGAATGTCCGCCCTATCAACTTTCGGGATGGTGGTCCGTCGCCTACCATGGTGACCTCACGGGTGACGGGGGAATCAGGGTTCGATTCCGGAGAGGAGCCTGAGAAACGGCTACCACATCCAAGGAAGGGGCAGCAGGCGCGCAAATTACCCATTTCCGACACGGAGAGGTAGTGACGAAAAATAACAATGCAGGTCTCTTTTCGAGGCCCTGCAATTGGAATGAGTGCATCCCAAACCCATGGGCGAGGACCCATTGGAGGGCAAGTCTGGTGCCAGCAGCCGTGGTAATTCCAGCCCAATAGCGTAAGCAACGTTGCTGCAGTTAAAAAGCTCGTAGTTGATCCGGATCACGCGCGGTCCGCCGCGAGGCGAGCCACCGCCGGTCCTGGACCCCAGGCCTCCCGGCGCCCCCGGATGCCCTTGACTGCGTCCTCGGCTTGGGGCCCGAGCGTTTACTTTGAAAAATTAGAGTGTTCAAGGCAGGGCCGGCACCGCGCCCCATTGAATACCCCAGCTAGAATAATGGAATAGGACCCCGGTTCTATTTTCTGTGGGGTTTCCGGAACCCGGGCCATGATCGAGAGGACGGCCGGGGGCATTCGTATTGCGCCGCTAGAGGTGAAATTTTGGACCGGCGCAAGACGGACCGAGCGAGTTGTTTGCCAAGAACGTTTCTCATTAATCAAGAACGAAAGCCGGAGGTTCGAAGACGATCAGATACCGTCGTAGTTCCGACCGTAAACGATGCCGACCCGCGATCCGGCGGCGTTTATTCCCATGACCCGCCGGGCAGCGTTGCGGGAAACCACGAGTCTCGTCCCAGGAGTAATGTTGCAAAGGTTGAAACCAGCAGCTAATTGACGGAAGGCACCACCAGAGGAGCCTGCGGCTTAATTTGACTCAACACGGGGAACCTCACCCGGCCCGACACGGAAAAAGATTGACAGATTGACGGCTCTTCTCGATTCTGTGGGTGGTGCATGGCCGTTCGTAGTTGTAGAGCGATTTGTCTGGTTGATTCCGATAACGAACGAGACTCTGGCATGCCAATTCAGTTACGCGGCCCCGCGCGTCGGCGTCTGCAAACTTCTTAGAGGGACGCGGCGTTCAGCCACGCGAGACTGAGCAATAACAGGTCTGTGATGCCCTTAGATGTCCGGGGCTGTATCGCGCCACAATGGCGGATCAATGTGCCTACCCTGCGCCGACAGGCGCGGGTAATTCATTGAACCCCGCCCGCATGGGGACCGATTGAAACTATTTCCCGAGAACGAGGAATTCCCAGTAAGCGCAGGTCATCAGCTTGCGTTGATTAAAGTCCTGCCTCTTGTACACACTGCCCGTCGCCACTACCGATTGAGCGGCTCAGTGAGGTCTCTCGGATCGGCTCCGCCCGCCCTTACCAGCCCTGGTGGAGCGCCGAGAAAGACGATCGAACTCTCGGTCGTTTAGAGGAAGTAAAAGTCGTAATGAAGGTTTCCGTAGGTGAACCCGGGCGGAAGGATCATTAATCGTAGTCGAGGGATCTCTCTCCTACGCCCAGAGGGCGAAGCCACGGTAAGCTTCGCGCGGTGCGGAAGTCCCTACGGGTCTACCTTCCCACCATCGCGCGCGAGCGTGATCGGTGATCCAAAGGTTGGCGTCGCGGCTCCCGGCAAATTGGTCTGACTTCACCTCGACCTGTCCCCTCTGCGGGGGGAGAGCGACGGAGGTGCGGGGCCGGGTTTGCAAGCACCTTCGCGCTTCCCACCCGGGGGAAGCAGAGAGGAGACGCCTGTCCCGGGGCCCCGCCGGCCGATGCTTAATTTCCCACCCTCCCGAAGCGTCCTCTGTCTCGACCGTAACGATTGAAAGAACGAAAACAAGAGTGTACAACTCTTAGCGGTGATCACTCGGCTCGTGCGTCGATGAAGAACGCAGCTAGCTGCGAGAACTACGTGAATTGCAGGACACACATTGATCATCGACCTTTCTGAACGCACATTGCGCCCCGTCCATCCCGGGGCCACGCCTGTCTGAGGGTCGCCTTGCTATCGATCGGACGGGGAAGAGTCGGTCTTCGTGCCGCCCTGCCCCCTCTGTCCGCGGCTGGAGCGTCGCAGACCCTGCCCTCCCGCGGTGGCCTACGTCCTCCCAAGTGCAGACCGCCGAACCGTTCGTCCATTGCCCGTTTGGGGGCGCGGCTCCCATTCTCTCCCCGGCGGCTGCCGGCGGTCTAACAGCTGCCCGCGCGCGGTGGACGGAGCTGCAGCAAACTCCCCTCGCGTTCGAGACGACGACAGCGCGATCGGTCGACGCGAGACCGGCGTCCCGCCCGGGACGACCGGCCGCCACCACCCGCCTGCTGGCCCACGACCTCAGCTCAGACGAGAAGACCCGCTGAATTTGGGCTATATTACTAAGCGGAGGAAAGAAACCAACCGGGATTCCCAGTAGCGGCGAGCGAAGAGGAAAGTCCAGCGCCGAACCCCGCCCTCTGCCGAGGGCGAGGGACCTGTGGCGTACGGAGGGCCGCCTTCGGCGCGTGGCCAGGGCCAAAGTCCTTCTGATGGAGGCTTAGCCCGCGGACGGTGTGAGGCGTCGGTGTCGGCCCCCGCCCCGCCGGGGTGCGGTTCCTCCCGGAGTCGGGTTGTTTGGGAATGCGCCCAAAGTGGGTGGTAAACTCCATCTAAGGCTAAATACCGGCACGAGACCGATAGCGGACAAGTACCGTGAGGGAAAGTTGAAAAGAACTTTGAAGAGAGAGTTCAACAGGGCGTGAAACCGTTAAGAGGTAAACGGGTGGGGACCGCACCGTCCGCCCGGTGGATTCAGCCCGGCGGGGGCGGGGTCGGCCTGTCCGGTGCGCGCTCCCTTCGCCCCCTCATTCCTGGGGTGGCGCGGGGTTGACGCCCGGGCGAAGGTCTCGGCCGCCGCCGGGGCACTTCCGCCGCGGTGGAGCGCCGCGACCGGCTCCGGTTCGGCTTGGAAGGGTCAGGGGGGCGAAGGTGGCCCGCGTCGGTTCAGGCCGTCGGGCTTTACAGCGCCCTCCCGCCCCGACTTCGCCGCTTGCTCTCCGGGGCCGCGGGTGAGTGTCCTCCGCGCCTCTCTGCCCTCCCTCCCTGTGGGAGGTGGGACGGGGGTCCCCGCCCCCGGCGTGGCGCGACAGGTGGACTGTCCTCAGTCCGCCCACGGCCGGGTGCCAAGCGCTGCCCAGGCGGGGATCCGACCCACGTTCGGGCGCCTGAGGTCTGCGGCGACGCCGCCTCCCACCCGACCCGTCTTGAAACACGGACCAGTGAGTCCAACGCGCGCGAGTCAGAGGGTGGCTCGCGAGCCCTGCGGCGCAATGAAGGTGAGAACGGGGGTCTCCCTGGTGGATCCCCGCCCCGGCGGGGGCGCACCACCGGCCCGCCTCGCGACCCTCCGGGGCAGGAGTTGGAGCGCGCGCGATGGCACCCGAAAGATGCAGAACTATGCCTGGGCAGGGCGGTCAGGGGAAACTCTGGTGGAGGCCCGCCGCGTCCTGACGTGCAATCGGTCGTCCGACCTGGGCATAGCGAAAGACTAATCGAACCATCTAGTAGCTGGTTCCTCCGAAGTTTCCTCAGGATACTGGCGCTCACGATCACAGTTTTATCCGGTAAAGCCAATGACTAGAGGCCTGTAATGAAACGGCCTCAACCTATTCTCAAACTTTAAATGGTAAGAGCCCGGCTCGCTGGCGCTGGAGCCGGGCGTGGAATGTGACGCGCCTAGTGGGCCATTTTTGGTAAGCAGAACTGGTGCTGCGGGATGAACTGAACAGTTGGTTAAGGCGCCCGATGCCGACGCTCATCAGACCCCATGTAAAGGTGTTGGTTGATATAGACAGCAGGACGGTGGCCATGGAAGTCGGCACCCGCCAAGGAGTGTAACAACTCACCTGCCGAATCAACCAGCCCTGAAATGGATGGCGCTGGAGCGTCGGGCCCATACCCGGCCGTCGACGGCAAGGGACACGCGAGCGGTCGCGCGCTGCAAGCCTCGACGAGTAGGGAGGGCCGCCGCGGTGGCGCCGAAGCCCAGGGCGCGGGTCTGCGGGAGCGCTGCGGGCGCAGATCTTGGTGGTAGTAGCAAATATTCAAACGAGAGCTTAGCCGAAGTGGAGAAGGGTTCCAGGAACAGCAGTTGAACATGGGCAGCCGGTCCTATGACGGGCCTACGCTGTTCGGAGGCGATGGCTCGTCGCTCCCCGTTCGACCGAAAGGAGTCGGGCCAGATCCCCGAGCCCGGAGCGGTGGAGACGGCGCCGCGAGGCGCCCAGTGCGGTGACGCAAACGACCCGGAGATGCCGATGGTGCCCCGGGAAAGAGTTCTCTTTCTTTGTGAAGGGCAGGGCGCCCTGGAACGGGTTCGCCCCGAGAGAGGGCCCGCGCCTGAAAGCGCCGCGCTTCTGGCGGCGTCCGTGAGCCTTCGTCGGCCCTTGAAAACTGGGGGAGAAGGTGTAAATCTCGCGCCGGGCCGTACCCATATCCGGCAGCAGGTCTCCAAGGTGAATCACGCCTCTGGCAAAGCTGGAACAAGCGCGCAGTAAGGGAAGTCGGCGTCAGATCCGTAACTTCGGGATAAGGATTGGCTCTTAAGGGTTGAGCCGGTCGGGCTGAGGTGCGAAGCGGGCCTGGGCCCGAGCCGCGACTGGGGGGAGCGGCCGCCCGGAGGTGCTCCTGACCCCGTTCCCGAGCCGGTGCGCGCGGTGGCGCGCGGTCCTCTCTCCCCCGCCGCGTCCGGGCCTCAAGCCTTTAGGTTCCCGACTCCCCGACTCTTCCCTCCGCCCTTCCCCGTCCGCGGGGTCGGGCGGGGGTCTCGGAGGAGGGGGGGGTGGAGAGGCCGGGGAGAAAGGGGTGGTTCGTGGCGTCGGGGCAACGGGAGGTCCTCGCCTGCCGCGCGATGCCTCCGCTGGCCGGGGCTCCGGTGGGGCGGATGCGCGGCTGGCGGCGGCGACCCTGGGCGCGCGCCGCGCCCTTCCTGCGGATTCCGCAGCTACGGCCCCGCGCCGGGGCCTGCGTCCGCGCGTGCGCCTCTCCGGGGAGGGCGCGCGGCCCTCGGCTCCCCCGGTGCGGGCGCCTCGCCGGCGGCTACAGCCAGCTTAGAACTGTCGCGGACCAGGGGAATCCGACTGTTTAATTAAACAGCATCGCGAAGGCCTCGGCGGTGTTCGACGCGATGTGATTTCTGCCCAGTGCTCTGAATGTCAAAGAAGAAATTCAACGAAGCGCGGCAAACGGCGGAGTAACTATGACTCTCTCTTAAGGTAGCCAAATGCCTGTCATCTAATTAGTGACGCATGAATGGATGGAAAACGAGAGATTCCACTGTCCCTACCAAGTGCCCATGAAACCATGCAGCCAAGAAATGTCATAGACCTGAAGGGGGAAAGAAGACCCCCTGTTGAGCTGACTCTGAAGTCTGCTCAGGGAAGAGACATGAGGGGTGTAGAATAAGTGGGAGTCCTGGCCCTGGGCCGGCCGCCGGTGAAATACCACTTACTCTTATCGTTTCCTCACTTACCTGGTGAGGAAGGCGTCCGCCCGGCGGGGGCCGCTCTGCTTCTGGCGTCGGGAGCGCCCCGGGGCCGTGCGGGGTGGGGCAACCCCCCTTCCCCTCCCGGCCGCTGACCAGTGCGACCCGCCCCGGGGACAGCGTCAGGTGGGAGTTTGACTGGGGCGGTACACCTGTCAAACGGTAACGCAGGTGTCCTAAGGCGAGCTCAGGGGGACAGAAACCTCCTGTAGAGCAGAAGGGCAAAAGCTCGCTTGATCTTGATTTTCAGTATGAGTACGGACTGCGAAAGCGGGGCCTCACGATCCTGGCTTTGGGTTTTGGTGGAGGTGTCGTAGTGCTACCACAGGGATAACTGGCTTGTGGCGGCCAAGCGTTCATAGCGACGTCGCTTTTGATCCTTCGATGTCGGCTCTTCCTATCATTGTAGAAGTTAGAATTCCACCCGTTGTTGATTGTTCACCCAATTAACAGGAACGTGAGCTAATAACTGTCGAGACAGGTTAGTTCATCCACTGATGTGAGCCGTTGTTGCAATAGTAATCCCGCTCAGTACGAGGAACCGCGACAGACATTTGTCTGGGTATGCGCTTGGCTGAGGAGCCACTGGCGCGAAGCCACCATCTGCAGGGATTATGACTGAACGCCTCTAAGTCAGAATCCCGCCTAAAAGCAACGATACAGCAGCGCCGTCGGACTCGATAGGCCCTGCAACCGGGAAGCCCTCTCCGGGGGCCCCGGCGCGGAGAGCCATTCGTGACAGAACCGGGGCGTGGCTAGAACGAGCGCCGCCCTTTCCCAGTGACGCACCGCATGCTTCATAGGGGAACCCCGGCGCTAAATGACTCGTAGACGACCTGATTCTGGGTCGGGGTGTCGTGCGTGGCAGAGCAGCTCTGTTGCTGCGATCCATTGAAAGTCAGCCCCGATCCAAGTTTTGTCGGGCCGGACGCAGGCGACAGGGGCCACCGGTCCGACAGACCCGCGGCGTGCGAAGGCAACAGGTTGCAGTCGGTGAAGGCTTGCCATCCTCGCCACTTGCGCGCGCAGTACGAGGCACCAGCGGCGGAGCGTGGCGCGATCCCAATACACCTGGGCACCAGCGGCAGAGGGAGGCTCAGCCTCGGTCCGCACCCCACAGCAAAGCCCCTACTGGGGCACCAGCAGCACGCAGCACTCATGTGGGGTAACCAGCGGCAGAAGGGGTTCTGTCCTGACCCTAACCCACAGCTGAAGCGGCGCACGAAAAACGCGAAAGCTCTCACGATAAATGCGAAAACGGTTAAACACCCATCTGAAACTGAACGAAGCG
